# Tunable Bias Signaling of the Angiotensin II Type 1 Receptor for Inotropy via C-Terminal Peptide Modifications and Allosteric Site Targeting

**DOI:** 10.1101/2025.08.13.670122

**Authors:** Margot Hadjadj, Justin Martel, Marie-Frédérique Roy, Malihe Hassanzadeh, Hugo Giguère, Alexandre Murza, Brian J. Holleran, Yoon Namkung, Ulrike Froehlich, Richard Leduc, Mannix Auger-Messier, Stéphane A. Laporte, Pierre-Luc Boudreault

## Abstract

The angiotensin II (AngII) type 1 receptor (AT1R) is a key prototypical G protein-coupled receptor in cardiovascular regulation. Biased agonists that activate G protein or β-arrestin pathways provide promising therapeutic potential, but the molecular determinants for this signaling bias and its physiological implications remain poorly understood. This study profiles AngII analogs with modifications at the C-terminal Phe^8^, revealing that analogs **11**, **12**, and **29a** exhibit varying degrees of Gα_q_ engagement while maintaining potent β-arrestin recruitment. Notably, **12** enhances left ventricular ejection fraction with minimal pressor responses in normotensive rats, while other analogs with variable Gα_q_ activity do not promote inotropy. Molecular modeling indicates that the unique profile of **12** results from its flexible long side chain engaging a deep allosteric pocket within AT1R. This study demonstrates that engineering AngII’s C-terminus enables selective tuning of AT1R signaling to control arterial versus cardiac responses, providing strategies for developing improved cardiovascular therapeutics.

**GRAPHICAL ABSTRACT:** 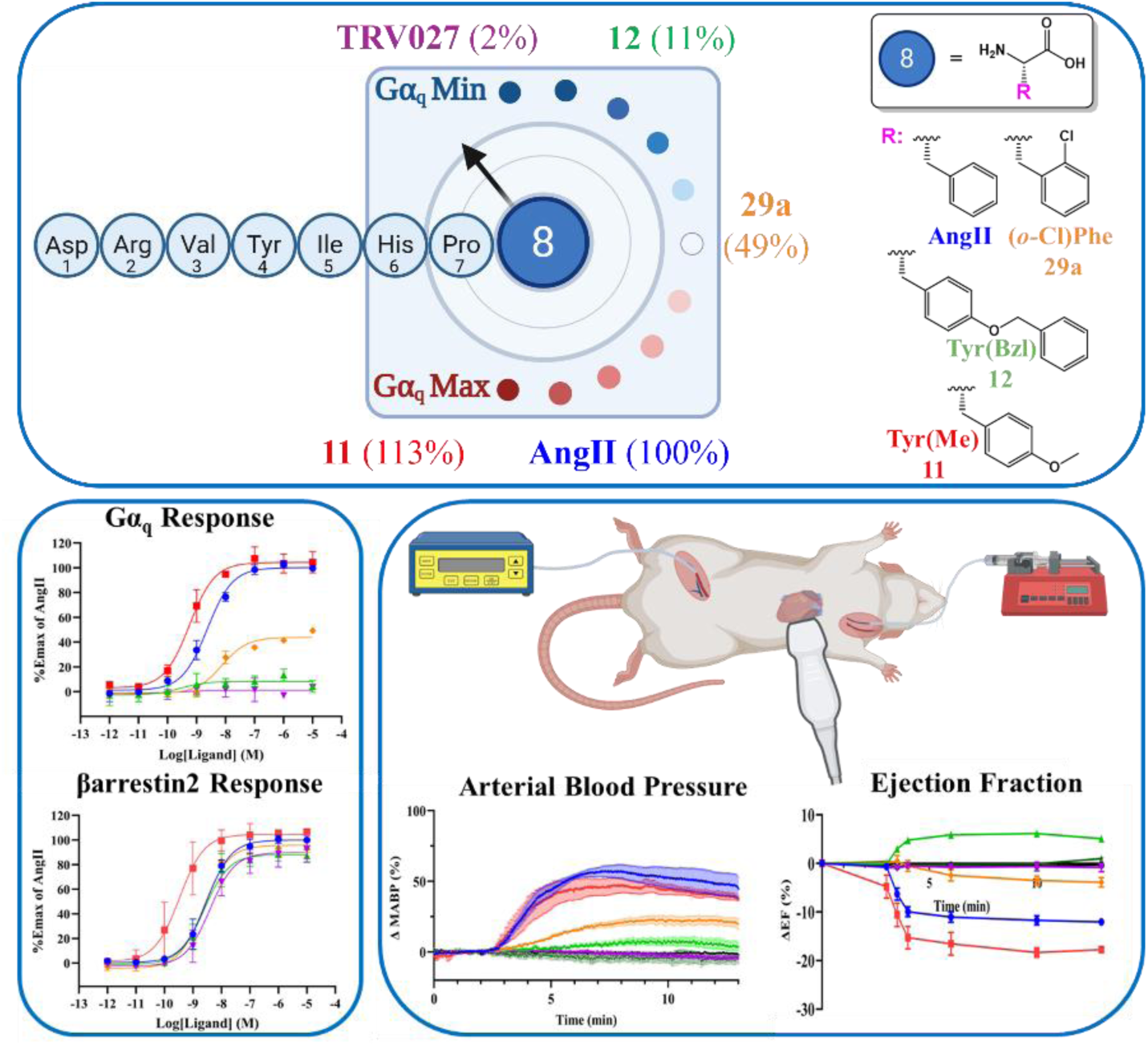

## INTRODUCTION

G protein-coupled receptors (GPCRs) are crucial targets for drug discovery. In fact, 30% of marketed medicines exert their therapeutic effects through interactions with one or more GPCRs.^1,2^ Among these, the Angiotensin II (AngII) Type 1 receptor (AT1R) has been particularly significant in developing therapeutics for cardiovascular disease.^3–5^ AT1R plays a crucial role in the renin-angiotensin-aldosterone system (RAAS), deemed the primary regulator of blood pressure.^6^ Its endogenous ligand, the octapeptide AngII, initiates both G protein-dependent (Gα_q/11_, Gα_i_, Gα_12/13_) and β-arrestin-dependent responses.^7–10^ The activation of Gα_q/11_ protein usually induces vasoconstriction and raises blood pressure, whereas the recruitment of β-arrestin to AT1R is associated with cardioprotective effects.^5,11^

AT1R blockers (ARBs, sartans) are commonly used to decrease blood pressure in hypertensive patients and function by inhibiting both G protein and β-arrestin signaling pathways.^12,13^ While this inhibition potentially contributes to their antihypertensive effects, it could also interfere with the protective mechanisms ostensibly mediated by β-arrestin.^14^ Therefore, in recent years, extensive efforts have been devoted to developing AngII ligands with functional selectivity (e.g., biased signaling properties). These ligands inhibit AT1R-mediated Gα_q/11_ protein activation while retaining β-arrestin coupling to the receptor.

Ligands with functional selectivity properties include the peptidergic AngII analog TRV027, modified at position 8, which exhibits bias to activate β-arrestin over Gα_q/11_.^13^ The benefit of these biased ligands would be to reduce afterload (e.g., vascular resistance) by inhibiting AT1R-mediated vasoconstriction while maintaining AT1R-driven inotropic effects in the heart and enhancing cardioprotection via β-arrestin signaling. This action may support cardiac output.^15,16^ Indeed, β-arrestin-biased AT1R ligands have proven to increase myocyte contractility *in vitro* and enhance cardiac contractility *in vivo*.^17^

While TRV027-related compounds exhibited promising results in preclinical models, they failed to deliver benefits in patients with acute heart failure (HF).^18^ The reason for this remains unclear but might be related to study design, the complexity of the disease, and/or our incomplete understanding of how the biased properties of AngII can be exploited *in vivo*. Nonetheless, post-analysis revealed that TRV027 reduced cardiovascular mortality in HF patients,^18^ spotlighting the potential benefits of leveraging new AngII ligands with optimized pharmacology for cardiovascular therapy.

Extensive evidence has emphasized the critical role of Phe⁸ in AngII-mediated AT1R activity.^19,20^ Alterations at this position significantly affect ligand binding affinity and receptor activation.^21^ Importantly, AngII analogs such as [Sar¹, Ile⁸]AngII and [Des-Phe⁸]AngII maintain AT1R binding but exhibit decreased receptor activation.^22^ Substitutions at Phe⁸ reduce the capability of AngII analogs to activate Gα_q/11_ but do not affect β-arrestin recruitment to AT1R.^8,23^ Such strategies have been used to develop β-arrestin biased agonists. Structural studies, in conjunction with molecular docking and molecular dynamics (MD) simulations, have also shown that Phe⁸ engages in hydrophobic and π-π stacking interactions with key AT1R residues, thereby stabilizing the active conformation.^24^ However, the high conformational heterogeneity seen in Phe⁸ substituted AngII analogs like TRV027, which relates to differential biased activity at AT1R,^24,25^ along with the discovery of a potential allosteric binding site beneath Phe⁸ that accommodates small-molecule binding,^26^ suggests that the chemical diversity at position 8 and the structural diversity of the receptor can be further utilized to regulate AT1R biased signaling and to enhance *in vivo* therapeutic effects.

Here, we comprehensively investigated the ability of AngII’s position 8 to modulate AT1R signaling by replacing Phe with a series of natural and unnatural amino acids (UAAs), each representing diverse chemical pharmacophores. We evaluated these analogs’ ability to activate AT1R’s Gα_q/11_ and β-arrestin signaling pathways *in vitro* and further assessed their *in vivo* properties for regulating blood pressure and left ventricular ejection fraction in rats. Our study identified new classes of β-arrestin-biased ligands with distinct Gα_q/11_ protein and β-arrestin pharmacological profiles. One exhibits partial Gα_q/11_ activity but does not show pressor effects *in vivo*, much like TRV027, while it improves heart ejection fraction. Molecular modeling of these ligands reveals an extension of the side chain and flexibility within a putative allosteric site in AT1R. We propose that such sites can be leveraged to design biased signaling ligands targeting AT1R, which could yield compounds with distinct *in vivo* properties beneficial for improved cardiovascular therapeutics.

## RESULTS AND DISCUSSION

### Synthesis and signaling of novel, position 8-substituted AngII analogs

We replaced AngII’s position 8 with amino acids of different physico-chemical properties such as lipophilic, polar, charged, and natural or unnatural aromatics (**Figure 1**). Incorporating UAAs not only fosters a broader chemical diversity, possibly promoting atypical ligand-dependent signaling, but it could also significantly improve their *in vivo* stability.^27^ To assess the AT1R-mediated signaling to Gα_q/11_ and β-arrestin of different AngII analogs compared to AngII, we expressed either the Gαβγ-Gα_q_-RlucII (GABY) or β-arrestin2-RLucII/rGFP-CAAX Bioluminescence Resonance Energy Transfer (BRET) sensors in the HEK293 cells, which were also transiently expressing the receptor, as previously done.^8,26^ Initially, we tested the 48 analogs based on their ability to engage Gα_q_ and recruit β-arrestin2, performing dose-response curves to determine the ligands’ efficacy (E_max_) and potency (log of the half-maximal effective concentration (EC_50_)), and compared them to the known β-arrestin-biased AngII analog TRV027.^13^ The substitution of Phe by aliphatic amino acids such as Ala (**1**), Nle (**2**), and Tle (**3**) resulted in β-arrestin engagement by AT1R similar to that observed with AngII, with EC_50_ values of 5.7, 0.86, and 2.3 nM, in comparison to 2.4 and 6.1 nM for AngII and TRV027, respectively (**Table 1**). Replacing Phe^8^ with polar/charged residues, as seen in analogs **5**, **6**, and **7**, impaired the ligands’ ability to activate Gα_q_ and substantially reduced the analogs’ potencies to promote β-arrestin’s recruitment to AT1R (100- to 1000-fold decrease), suggesting a lipophilic or aromatic residue is necessary at this position to engage these pathways.^24^ Interestingly, replacing position 8 with Cha (**4**) created an analog with partial activity on Gα_q_ (i.e., 52% of AngII) that retained high potency for both Gα_q_ and β-arrestin activation. Linear lipophilic substitutions led to minimal Gα_q_ engagement, indicating the importance of aromatic π-stacking-type interactions for preserving the AT1R engagement of this pathway.

**Figure 1.**
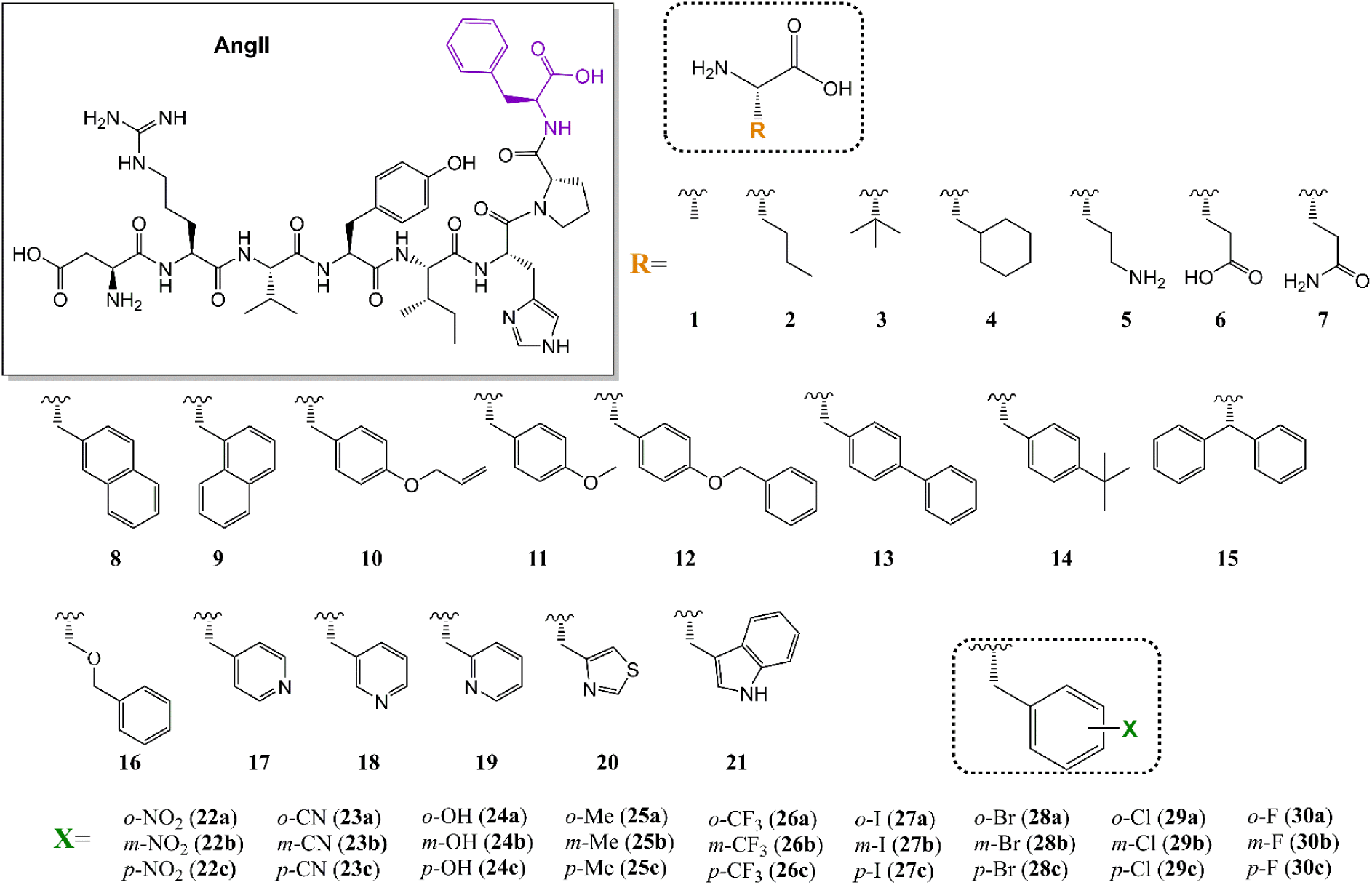
Analogs of AngII with various natural and unnatural amino acid substitutions at position 8.

**Table 1.** Potency (EC_50_) and relative efficacy (E_max_) of lipophilic, polar, or charged AngII analogs for Gα_q_ and βarr2 signaling pathways.

| Compounds | Peptide sequence | EC<sub>50</sub> Gα<sub>q</sub> (nM)<br>(E<sub>max</sub> %<br>AngII) | EC<sub>50</sub> βarr2 (nM)<br>(E<sub>max</sub> % AngII) |
| --- | --- | --- | --- |
| AngII | D-R-V-Y-I-H-P-F | 2.7±0.6 (100) | 2.4±0.3 (100) |
| TRV027 | **Sar**-R-V-Y-I-H-P-**a** | >10000 (2) | 6.1±1.1 (97) |
| 1 | D-R-V-Y-I-H-P-**A** | >10000 (4) | 5.7±1.0 (78) |
| 2 | D-R-V-Y-I-H-P-**Nle** | 3.0±3.6 (36) | 0.86±0.16 (92) |
| 3 | D-R-V-Y-I-H-P-**Tle** | 0.36±0.84 (19) | 2.3±0.4 (81) |
| 4 | D-R-V-Y-I-H-P-**Cha** | 5.5±3.9 (52) | 1.4±0.2 (95) |
| 5 | D-R-V-Y-I-H-P-**Orn** | >10000 | 4558±3877 (45) |
| 6 | D-R-V-Y-I-H-P-**E** | >10000 | 957±111 (63) |
| 7 | D-R-V-Y-I-H-P-**Q** | >10000 | 158±20 (56) |

E<sub>max</sub> was obtained from the response of the ligand at a concentration of 10 μM. EC<sub>50</sub> are presented as mean ± SEM of at least three to five independent experiments.

We then evaluated the roles of the aromatic moiety at position 8 in its ability to interact with AT1R and engage with Gα_q_ and β-arrestin. This was done by replacing Phe with a series of aromatic UAAs of different lengths and bulkiness (analogs **8** to **21**, **Table 2** and **Figure 1**). Most alterations did not negatively impact the recruitment of β-arrestin to AT1R, with the notable exceptions of compounds **13** (Bip) and **14** (*p*-tBuPhe). These compounds displayed an 8–10 fold decrease in potency (EC_50_ 15 nM and 26 nM, respectively) in comparison to AngII. Notably, the replacement of Phe^8^ with Tyr(Me) (**11**) resulted in a 6-fold increase in potency for β-arrestin recruitment relative to AngII (EC_50_ 0.36 vs 2.4 nM) and demonstrated increased potency and efficacy in activating Gα_q_ (EC_50_ 1.1 nM and 113%, respectively). The substitution of the methyl group in the *para* position with a benzyl group (**12**) invalidated Gα_q_ but preserved β-arrestin activation (2.5 nM vs 2.4 nM for AngII, **Table 2**). This substitution even slightly improved the response compared to TRV027 (6.1 nM). These results imply that the *para* position of the Phe^8^ residue can be extended to fully support AT1R-mediated β-arrestin responses, but not Gα_q_ activation. Unexpectedly, the rigid positioning of aromatic groups with the Dip substitute (**15**) was not detrimental to Gα_q_ activation by AT1R (EC_50_ 7.9 nM, E_max_ 99%). This was contrary to Ser(Bzl) (**16**) and Tyr(Bzl) (**12**) residues that both lost their ability to engage that pathway (i.e., EC_50_ >10 000 nM). Taken together, these findings suggest that the receptor’s orthosteric binding pocket, which interacts with the C-terminus of AngII, can accommodate extended and flexible side chains while maintaining β-arrestin selective responses.

**Table 2.** Potency (EC_50_) and relative efficacy (E_max_) of aromatic AngII analogs for Gα_q_ and βarr2 signaling pathways.

| Compounds | Peptide sequence | EC<sub>50</sub> G$\alpha_q$ (nM)<br>(E<sub>max</sub> % AngII) | EC<sub>50</sub> βarr2 (nM)<br>(E<sub>max</sub> % AngII) |
| --- | --- | --- | --- |
| AngII | D-R-V-Y-I-H-P-F | 2.7±0.6 (100) | 2.4±0.3 (100) |
| TRV027 | **Sar**-R-V-Y-I-H-P-**a** | >10000 (2) | 6.1±1.1 (97) |
| 8 | D-R-V-Y-I-H-P-**1Nal** | 8.3±5.1 (61) | 2.1±0.4 (101) |
| 9 | D-R-V-Y-I-H-P-**2Nal** | 3.9±1.9 (69) | 1.0±0.3 (97) |
| 10 | D-R-V-Y-I-H-P-**Tyr(All)** | 17±10 (48) | 7.4±1.9 (100) |
| **11** | D-R-V-Y-I-H-P-**Tyr(Me)** | **1.1±0.4 (113)** | 0.36±0.10 (105) |
| **12** | D-R-V-Y-I-H-P-**Tyr(Bzl)** | **>10000 (11)** | 2.5±0.4 (89) |
| 13 | D-R-V-Y-I-H-P-**Bip** | 16±12 (45) | 15±4 (98) |
| 14 | D-R-V-Y-I-H-P-***p*-tBuPhe** | 45±29 (40) | 26±4 (104) |
| 15 | D-R-V-Y-I-H-P-**Dip** | 7.9±1.7 (99) | 6.7±0.8 (109) |
| 16 | D-R-V-Y-I-H-P-**Ser(Bzl)** | 2.3±4.0 (26) | 3.0±0.5 (87) |
| 17 | D-R-V-Y-I-H-P-**4Pyridyl** | 7.5±6.3 (37) | 5.2±1.0 (105) |
| 18 | D-R-V-Y-I-H-P-**3Pyridyl** | 22±11 (31) | 3.7±0.7 (90) |
| 19 | D-R-V-Y-I-H-P-**2Pyridyl** | 6.9±2.6 (48) | 2.2±0.4 (101) |
| 20 | D-R-V-Y-I-H-P-**4ThzAla** | 14±13 (29) | 1.9±0.3 (97) |
| 21 | D-R-V-Y-I-H-P-**W** | 4.7±1.1 (91) | 2.2±0.4 (107) |

E<sub>max</sub> was obtained from the response of the ligand at a concentration of 10 μM. EC<sub>50</sub> are presented as mean ± SEM of at least three to five independent experiments.

We next aimed to explore the relationship between the C-terminal aryl group’s electronicity^28^ and Gα_q_/β-arrestin engagement. The inclusion of an electron-richer pyridyl group instead of phenyl (analogs **17** to **19**) led to a minor decrease in Gα_q_ activation (EC_50_ 6.9 to 22 nM) with a more prominent efficacy decrease (E_max_ <50%). The β-arrestin response from these ligands suffered fewer effects, with only 1.5-2-fold potency decreases compared to AngII, without any impact on their efficacy. This observation was corroborated with the 4ThzAla analog (**20**), which also resulted in a notable decrease in Gα_q_ activation (EC_50_ 14 nM, E_max_ 29%), but had no impact on the β-arrestin response. Lastly, the integration of an H-bond donor aromatic residue, like Trp (**21**) at position 8, did not hinder either the Gα_q_ or β-arrestin AT1R-mediated responses (EC_50_ 4.7 nM, E_max_ 91%) compared to AngII. In summary, these results emphasize the essential role of the lipophilic/aromatic interactions at position 8 in managing Gα_q_ and β-arrestin engagement by AT1R. Additionally, they show how the electronic and steric properties of this residue can be selectively adjusted to modulate pathway activation differently, allowing for the design of ligands with distinct signaling profiles.

### Probing *ortho*, *meta*, and *para* substitutions of Phenylalanine at position 8

To better understand whether the position of the substituent could also act as a chemical switch, we used atoms/groups of various sizes such as halides, H-bond donors/acceptors, and methylene. Substituting either the *ortho* (analogs **22a** to **30a**), *meta* (**22b** to **30b**), or *para* (**22c** to **30c**) positions with different pharmacophores size (CF_3_ >NO_2_ >Me >CN >I >Br >Cl >OH >F >H) led to a β-arrestin response similar to that of AngII (**Table 3**). However, for the *ortho* position, significant drops in Gα_q_ efficacies were observed (ranging from 23% to 57% for **23a** and **25a**, respectively), except for the hydroxyl (*o*-OH, **24a**) and fluorine (*o*-F, **30a**) derivatives, which exhibited respective EC_50_ values of 2.1 nM (100%) and 1.0 nM (101%). This full activation, akin to that of the endogenous ligand, appears achievable only with smaller substituents, suggesting that steric hindrance is likely the most important factor. The comparison between cyano (*o*-CN, **23a**) and hydroxyl (*o*-OH, **24a**) groups provides compelling evidence supporting the hypothesis that the size and the electronic nature of the substituents influence the orientation of the aromatic ring, thereby affecting the Gα_q_ signaling.

**Table 3.**
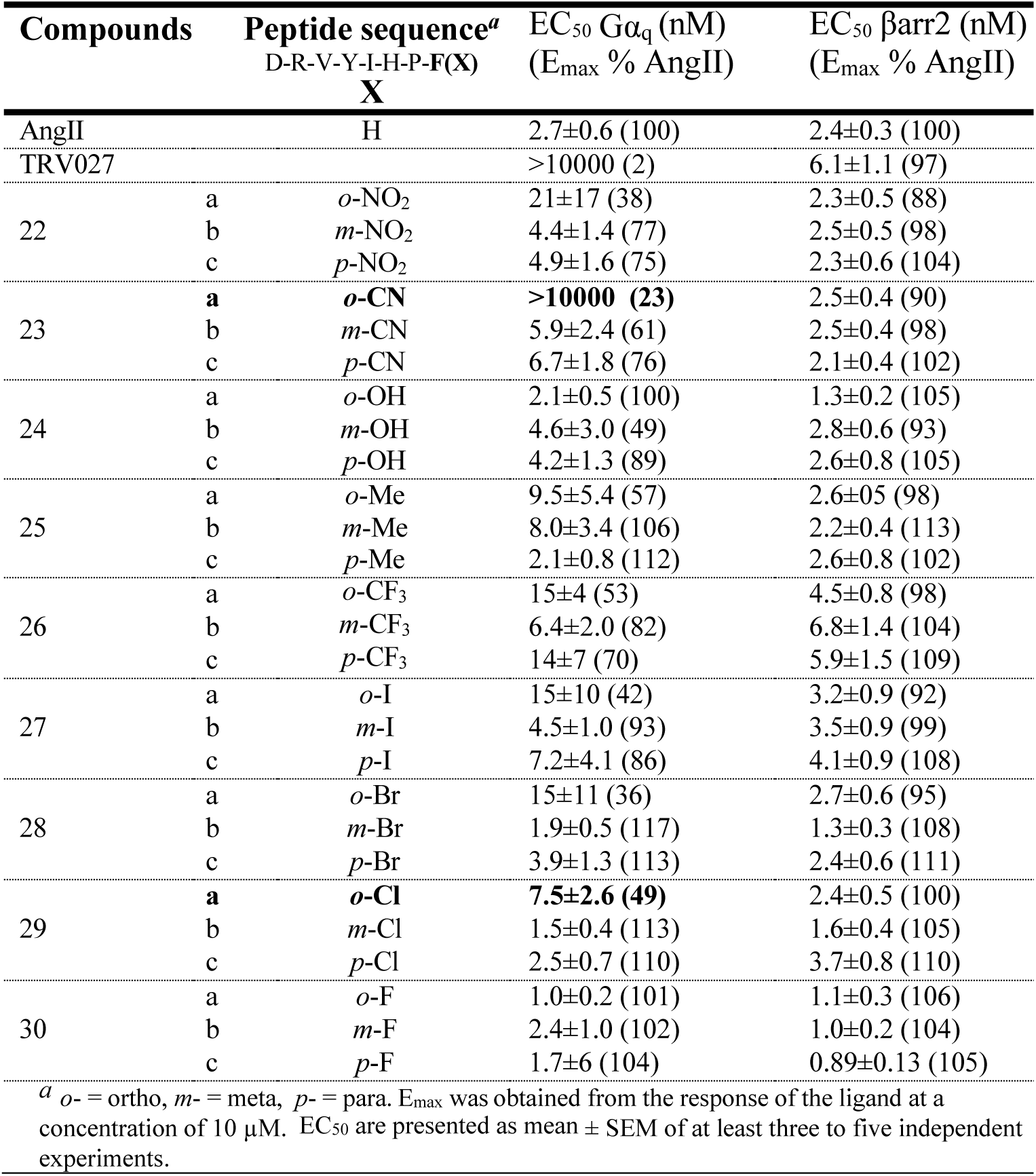
Potency (EC_50_) and relative efficacy (E_max_) of *ortho*-, *meta*-, *para*-substituted aromatic AngII analogs for Gα_q_ and βarr2 signaling pathways.

Conversely, the *meta* position appears to be more accommodating, as halides of varying sizes fully activate Gα_q_ protein engagement (EC_50_ 4.5, 1.9, 1.5, and 2.4 nM) for (*m*-I, **27b**), (*m*-Br, **28b**), (*m*-Cl, **29b**), and (*m*-F, **30b**), respectively. Electron-donating or -withdrawing groups, such as nitro (*m*-NO_2_, **22b**), cyano (*m*-CN, **23b**), and hydroxyl (*m*-OH, **24b**), caused a reduction in efficacies (E_max_ of 77%, 61%, and 49%, respectively) and potencies (EC_50_ of 4.4 nM, 5.9 nM, and 4.6 nM, respectively), which might be attributed to detrimental H-bond donor/acceptor properties. Lastly, as expected (**Table 2**), minor substituents at the *para* position did not hinder Gα_q_ engagement. Substituents such as methylene (**25c**) or halides (**27c** to **30c**) were well tolerated, and only analog **26c** (*p*-CF3) exhibited a 5- to 6-fold loss of potency relative to AngII. Furthermore, the presence of small substituents at the *para* position did not affect Gα_q_ (e.g., *p*-Cl, **29c**, EC_50_ 2.5 nM; E_max_ 110% or *p*-F, **30c**, EC_50_ 1.7 nM; E_max_ 104%), in contrast to polar groups (*p*-NO_2_, **22c**), or (*p*-CN, **23c**) that led to a slight reduction of E_max_ to ∼75%.

### Assessing the pharmacological signaling profiles of AngII analogs

Next, we compared different analogs’ signaling profiles and assessed how they diverged from AngII when expressed through AT1R in a heterologous system. The Gα_q_ and β-arrestin BRET sensors, as well as PKC and Rho BRET sensors, were employed to discern downstream Gα_q_ responses from the analogs, which would be intensified by signaling cascades to these effectors (**Figure 2**).

**Figure 2.**
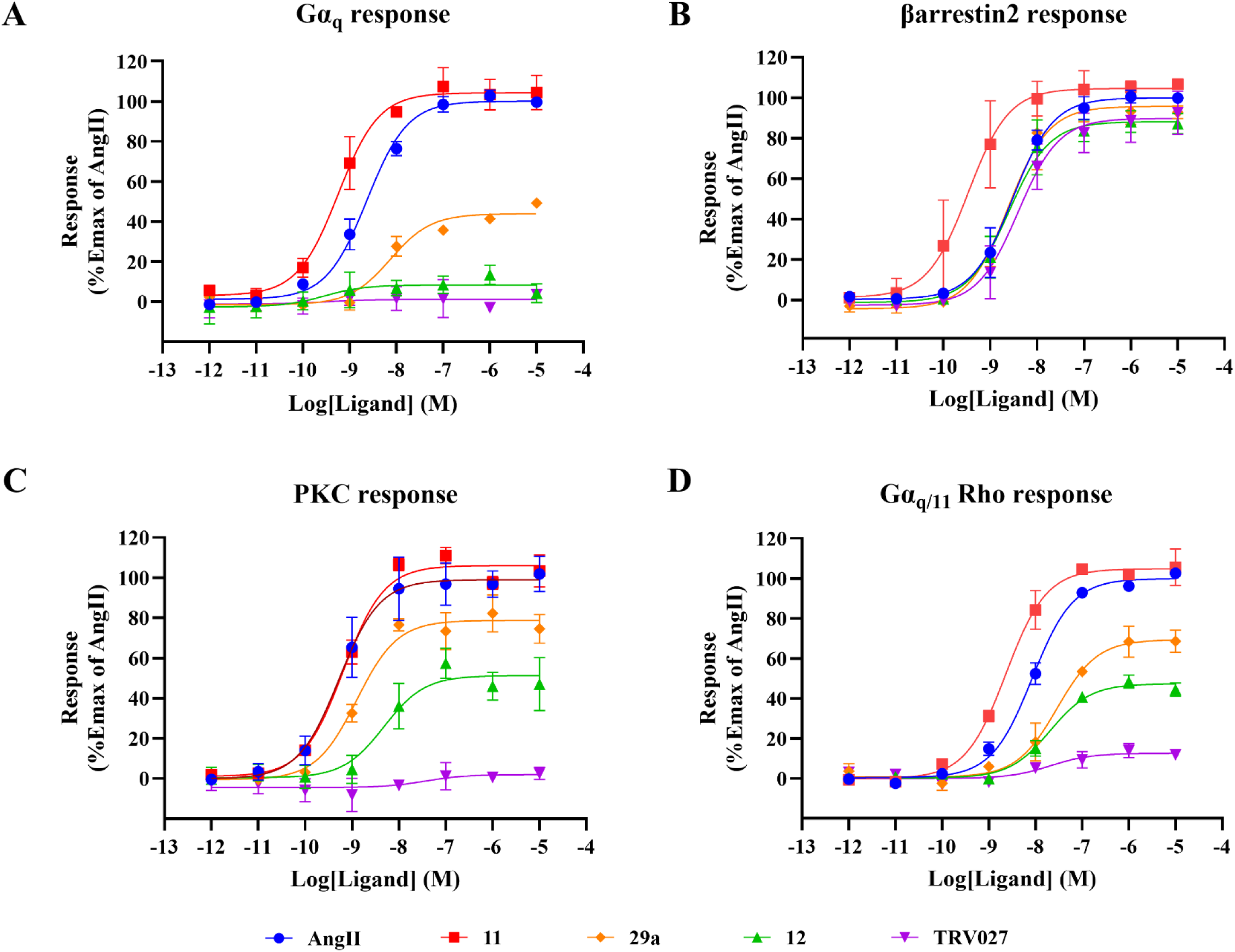
Dose-response curves of selected AngII analogs for activating AT1R downstream pathways. HEK293 cells expressing either the Gα_q_ (**A**), βarrestin2 (**B**), PKC (**C**), or Rho (**D**) BRET sensor along with AT1R were stimulated with increasing concentrations of AngII or analogs. Cells were transfected and assayed as indicated in the Experimental Section. BRET measurements were recorded and normalized to the response of AngII in the same experiment and expressed as %Emax of AngII. Data are means ± SEM of at least three to five independent experiments.

As before,^29^ the Rho BRET sensor was expressed in HEK293 cells, which lack Gα_12/13_, alongside the AT1R to solely measure Gα_q_’s activity, because both G protein subtypes can engage Rho via AT1R.^8^ We honed in on analogs **11** and **12** – the former demonstrated high potency and efficacy in engaging both Gα_q_ and β-arrestin through AT1R, while the latter maintained high efficacy and potency relative to AngII for β-arrestin activation, but showed no discernible Gα_q_ activity with the GABY sensor. We also chose analog **29a** to stand for a partial agonist on Gα_q_ and a full agonist with high potency for β-arrestin engagement. Results were gauged against TRV027, which appeared to have no noticeable activity on Gα_q_ but showed full activity on β-arrestin. We began by determining the binding affinities (*K*_i_) of all analogs and found they were in the low nanomolar range, varying from 0.23 nM to 1.4 nM, with TRV027 showing the lowest apparent affinity of all AngII analogs (6.6 nM, **Table 4**).

**Table 4.** Apparent affinity of AngII analogs and their potency (EC_50_) and relative efficacy.

| Compounds | Peptide sequence | $K_i$ (nM) | $EC_{50}$ $G\alpha_q$ (nM)<br>($E_{max}^a$ % AngII) | $EC_{50}$ βarr2 (nM)<br>($E_{max}$ % AngII) | $EC_{50}$ PKC (nM)<br>($E_{max}$ % AngII) | $EC_{50}$ Rho $G\alpha_{q/11}$ (nM)<br>($E_{max}$ % AngII) |
| --- | --- | --- | --- | --- | --- | --- |
| AngII | D-R-V-Y-I-H-P-F | 0.47±0.07 | 2.7±0.6 (100) | 2.4±0.3 (100) | 0.53±0.11 (100) | 8.6±1.0 (100) |
| 11 | D-R-V-Y-I-H-P-Tyr(Me) | 0.23±0.18 | 1.1±0.4 (113) | 0.36±0.10 (105) | 0.66±0.13 (105) | 2.3±0.5 (105) |
| 29a | D-R-V-Y-I-H-P-(o-Cl)F | 1.4±0.4 | 7.5±2.6 (49) | 2.4±0.5 (100) | 1.3±0.4 (79) | 29±10 (68) |
| 12 | D-R-V-Y-I-H-P-Tyr(Bzl) | 1.4±0.2 | >10000 (11) | 2.5±0.4 (89) | 4.8±3.4 (51) | 21±5 (47) |
| TRV027 | Sar R-V-Y-I-H-P-a | 6.6±3.6 | >10000 (2) | 6.1±1.1 (97) | >10000 (3) | >10000 (13) |

<sup>a</sup> $E_{max}$ was obtained from the response of the ligand at a concentration of 10 μM. Data are presented as mean ± SEM of at least three to five independent experiments.
( $E_{max}$ ) for $G\alpha_q$ , βarr2, PKC, and $G\alpha_{q/11}$ Rho signaling pathways.

Interestingly, although we did not register Gα_q_ activity with the GABY sensor for **12**, it acted as a partial agonist when assessed using PKC or Rho sensors, which amplify signals originating from Gα_q_. (**Figure 2** and **Table 4**). Similarly, while the GABY sensor detected no clear Gα_q_ activity for TRV027, a marginal response from these Gα_q_ downstream effectors was recorded by the PKC and Rho sensors. However, this response was significantly dampened compared to **12**. Analog **29a** persistently exhibited robust partial agonist activity on Gα_q_ when evaluated using PKC or Rho sensors.

We determined the relative activity (RA) of each analog compared to AngII, applying the Δlog(τ/KA) method for Gα_q_ and β-arrestin activation, as previously done (**Figure 3A** and **Table 5**).^8^

**Figure 3.**
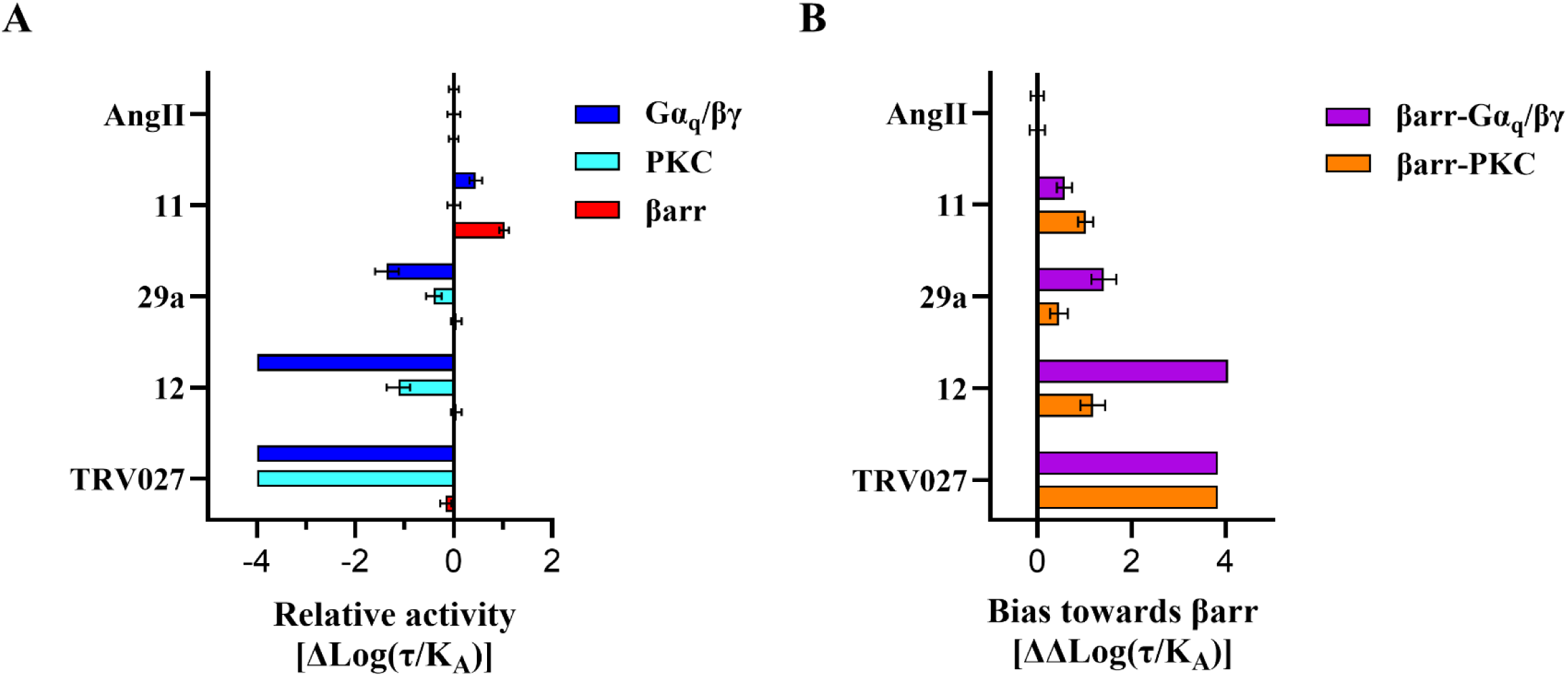
Relative activity (Δlog(τ/*K*_A_) and bias of AngII analogs for activating AT1R downstream pathways. (**A**) Relative activities [ΔLog(τ/*K*_A_)] of each ligand to AngII, the reference ligand. ΔLog(τ/*K*_A_) were calculated by subtracting the log(τ/*K*_A_) value of AngII in each pathway. (**B**) Bias of the ligand toward the β-arrestin pathway was determined by subtracting each ligand’s Δlog(τ/K_A_) value in Gα_q_/βγ or PKC pathways from that of the βarr2 pathway (ΔΔlog(τ/*K*_A_). Data represent means ± SEM of at least three to five independent experiments.

**Table 5.** Relative activities (ΔLog(τ/K_A_) and bias factors of AngII and analogs.

| Compounds | Log($\tau/K_A$) | | | Relative activity, $\Delta\log(\tau/K_A)$ | | | Bias ($\Delta\Delta\log(\tau/K_A)$) | |
| --- | --- | --- | --- | --- | --- | --- | --- | --- |
| | Gq | PKC | βarr2 | Gq | PKC | βarr2 | βarr2-Gq | βarr2-PKC |
| AngII | 8.69±0.07 | 9.24±0.09 | 8.51±0.07 | 0.00±0.10 | 0.00±0.13 | 0.00±0.10 | 0.00±0.14 | 0.00±0.16 |
| 11 | 9.13±0.11 | 9.24±0.09 | 9.54±0.07 | 0.45±0.13 | 0.00±0.13 | 1.03±0.10 | 0.58±0.16 | 1.03±0.16 |
| 29a | 7.32±0.23 | 8.83±0.13 | 8.56±0.09 | -1.36±0.24 | -0.41±0.16 | 0.05±0.11 | 1.41±0.27 | 0.46±0.19 |
| 12 | n.d. | 8.11±0.22 | 8.57±0.08 | -4<sup>a</sup> | -1.12±0.24 | 0.05±0.11 | 4.05 | 1.18±0.26 |
| TRV027 | n.d. | n.d. | 8.35±0.09 | -4<sup>a</sup> | -4<sup>a</sup> | -0.17±0.12 | 3.83 | 3.83 |

n.d., not determined due to lack of response; <sup>a</sup>, assigned an arbitrary RA of -4, because EC<sub>50</sub> or Emax could not be determined.

To calculate biases (ΔΔlog(τ/KA) between the Gα_q_ and β-arrestin pathways, we used either the Gα_q_ GABY or the amplified PKC responses (**Table 5** and **Figure 3B**), and compared these to AngII and its analogs. Analog **11** exhibited the strongest increase in RA for the β-arrestin response, while **12**, **29a**, and TRV027 showed varied decreases in RAs for Gα_q_. The rank order from the lowest to the highest activity for the Gα_q_ pathway was TRV027, **12**, and **29a**. Interestingly, even though **11** and **12** exhibited very different RAs for Gα_q_ and β-arrestin activation, they both demonstrated a similar bias towards the β-arrestin pathway when using the PKC sensor, albeit to a lesser extent than TRV027 (**Figure 3**). These results suggest that while ligands may exhibit similar signaling biases, they influence downstream pathways differently, potentially leading to distinct *in vivo* outcomes.

We next assessed the analogs’ ability to engage Gα_q_ by assessing either the PKC or Rho responses in rat primary vascular smooth muscle cell (VSMC) culture, expressing low levels of endogenous AT1R (between 25–50 fmol/mg) (**Figure 4**, **Table 6**). The Rho response in VSMC is solely dependent on Gα_q_ activity, as it is completely blocked by a selective Gα_q_ inhibitor, YM-254890 (**Figure 4C**). Using either sensor, **11** acted as a full agonist with similar potency and efficacy to AngII (**Figures 4A-B**). Analog **29a** behaved as a partial agonist, akin to responses in HEK293 cells, albeit with reduced potency (4.1 nM up to 138 nM) (**Figure 4**, **Table 6**). Analog **12** retained some partial Gα_q_ activity, while TRV027 showed no activity on that pathway when examined with either the PKC or Rho sensor. Strikingly, the analogs elicited comparable responses across both sensors, indicating that the Gα_q_-PKC and Gα_q_-Rho pathways are equally and efficiently engaged in VSMCs. This suggests that both signaling pathways contribute to the vascular cell response,^30,31^ even in the context of low AT1R expression. Notably, the rank order of ligand ability in engaging Gα_q_ was preserved between VSMCs and HEK293 cells. However, in HEK293 cells, the potency and/or efficacy of many ligands were enhanced, likely reflecting the higher receptor expression.

**Figure 4.**
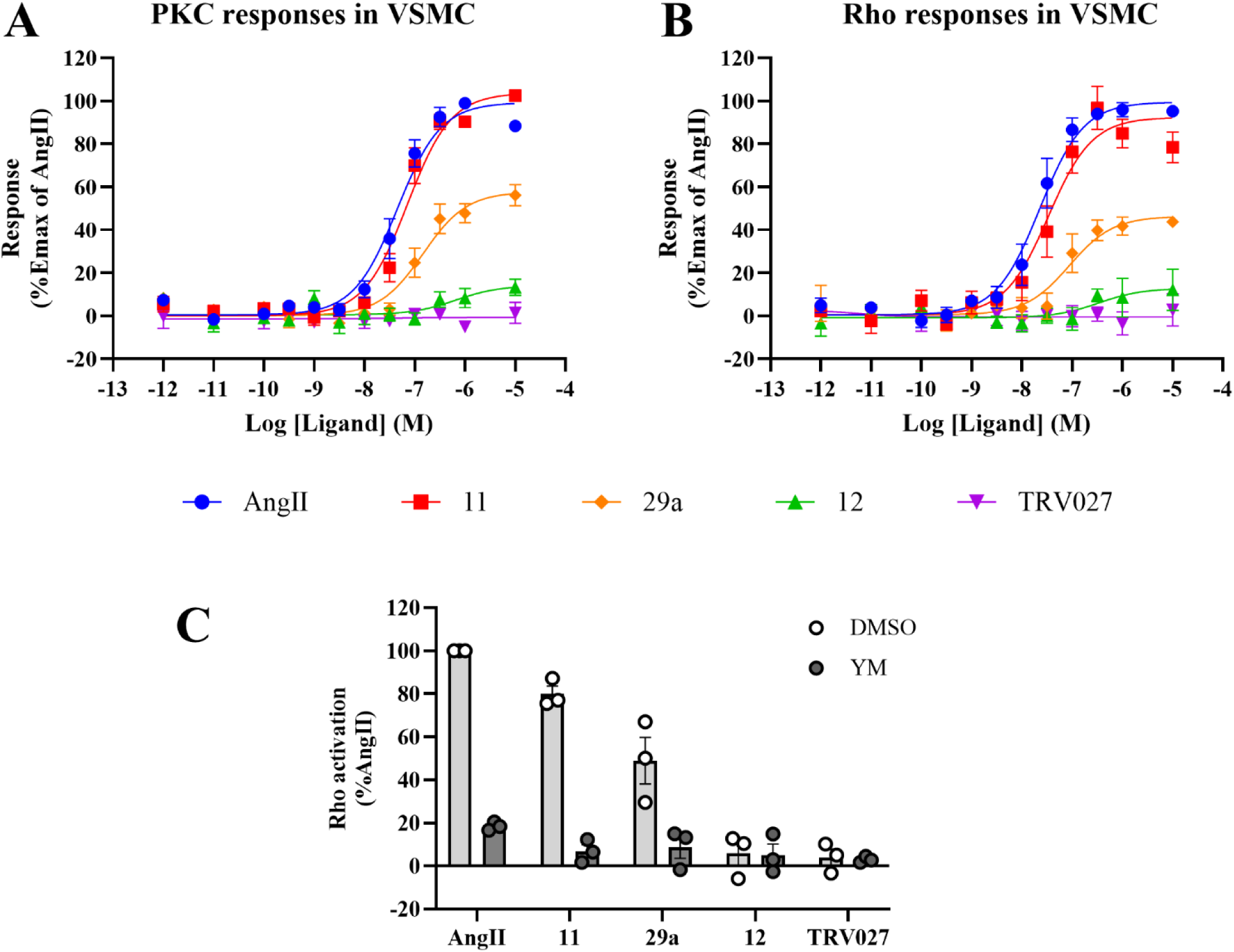
Dose-response curves for PKC and Rho activation in rats. Cells were transduced with either PKC- (**A**) or PKN-RlucII/rGFPcaax (**B**) adenoviral expressing sensors and assayed as indicated in the Experimental Section. BRET measurements were recorded and normalized to the response of AngII in the same experiment and expressed as %Emax of AngII. Data are means ± SEM of at least three independent experiments. (**C**) Rho sensor transduced VSMC cells were pretreated with either 400 nM of YM-254890 or DMSO (final 0.4%) for 30 min, then cells were stimulated with 1 μM of the indicated ligands for 1 min 30 s prior to BRET measurements. Ligand-mediated BRET changes were normalized to that of AngII in DMSO (%AngII). Data were expressed as means ± SEM with individual values.

**Table 6.** Apparent affinity of AngII analogs and their potency (EC_50_) and relative efficacy (E_max_) for PKC and Rho signaling pathways in VSMCs.

| Compounds | EC<sub>50</sub> PKC (nM)<br>(E<sub>max</sub> % AngII) | EC<sub>50</sub> Rho (nM)<br>(E<sub>max</sub> % AngII) |
| --- | --- | --- |
| AngII | 52.4±13.5 | 25.8±8.8 |
| 11 | 79.8±17.7 (104) | 39.1±11.0 (92) |
| 29a | 188.3±53.6 (58) | 118.5±65.6 (46) |
| 12 | >10000 (13<sup>a</sup>) | >10000 (12<sup>a</sup>) |
| TRV027 | >10000 (1<sup>a</sup>) | >10000 (3<sup>a</sup>) |

<sup>a</sup> E<sub>max</sub> was obtained from the response of the ligand at a concentration of 10 μM. Data represent means ± SEM of at least three to five independent experiments.

### Assessing AngII analogs’ effect on blood pressure and cardiac inotropy

Given the distinct signaling profiles and biases of the AngII analogs, some of which exhibited similar biases (e.g., **11** and **12**), we next investigated their acute cardiovascular effects *in vivo*. AngII is known to rapidly alter both parameters in animal models, so we focused on the analogs’ blood pressure effects and regulation of cardiac ejection fraction.^32–35^ We compared **11, 29a, 12**, and TRV027 effects to Olmesartan, a known AT1R small-molecule antagonist with antihypertensive effects, but with no inotropic functions.^16,36^ All ligands were administered intravenously at the same dose (0.96 nmol kg^−1^ min^−1^), by instrumentalizing adult Sprague-Dawley rats for simultaneous recordings of arterial blood pressure and cardiac function (**Figure 5A**). Plasma half-lives of **11**, **12**, and **29a** were similar to AngII (∼30 min) (see the Supporting Information), suggesting that any differences from *in vivo* effects are unlikely to result from inherent stability discrepancies. As expected from previous findings in rats,^37–39^ constant infusion of AngII provoked a rapid and sustained increase in arterial blood pressure (**Figures 5B-C**), together with a significant decrease in cardiac ejection fraction of the left ventricle (**Figures 5D-E**). In these conditions, infusion of Olmesartan did not alter the cardiovascular parameters measured in normotensive rats (**Figures 5B-E**), consistent with previous findings.^40^ Although **11** increased blood pressure as rapidly and to the same extent as AngII (**Figures 5B-C**), it further reduced the left ventricular ejection fraction by approximately 1.5-fold compared to AngII (**Figures 5D-E**). Analog **29a** partially increased blood pressure to about half that of AngII (**Figures 5B-C**), which aligns with its partial effectiveness in activating Gα_q_ (**Table 4**). Additionally, it consistently reduced cardiac contractility, although this reduction did not reach statistical significance compared to the control vehicle administration (**Figures 5D-E**). Analog **12** negligibly increased blood pressure, while TRV027 and Olmesartan did not significantly affect either blood pressure or inotropic responses.

**Figure 5.**
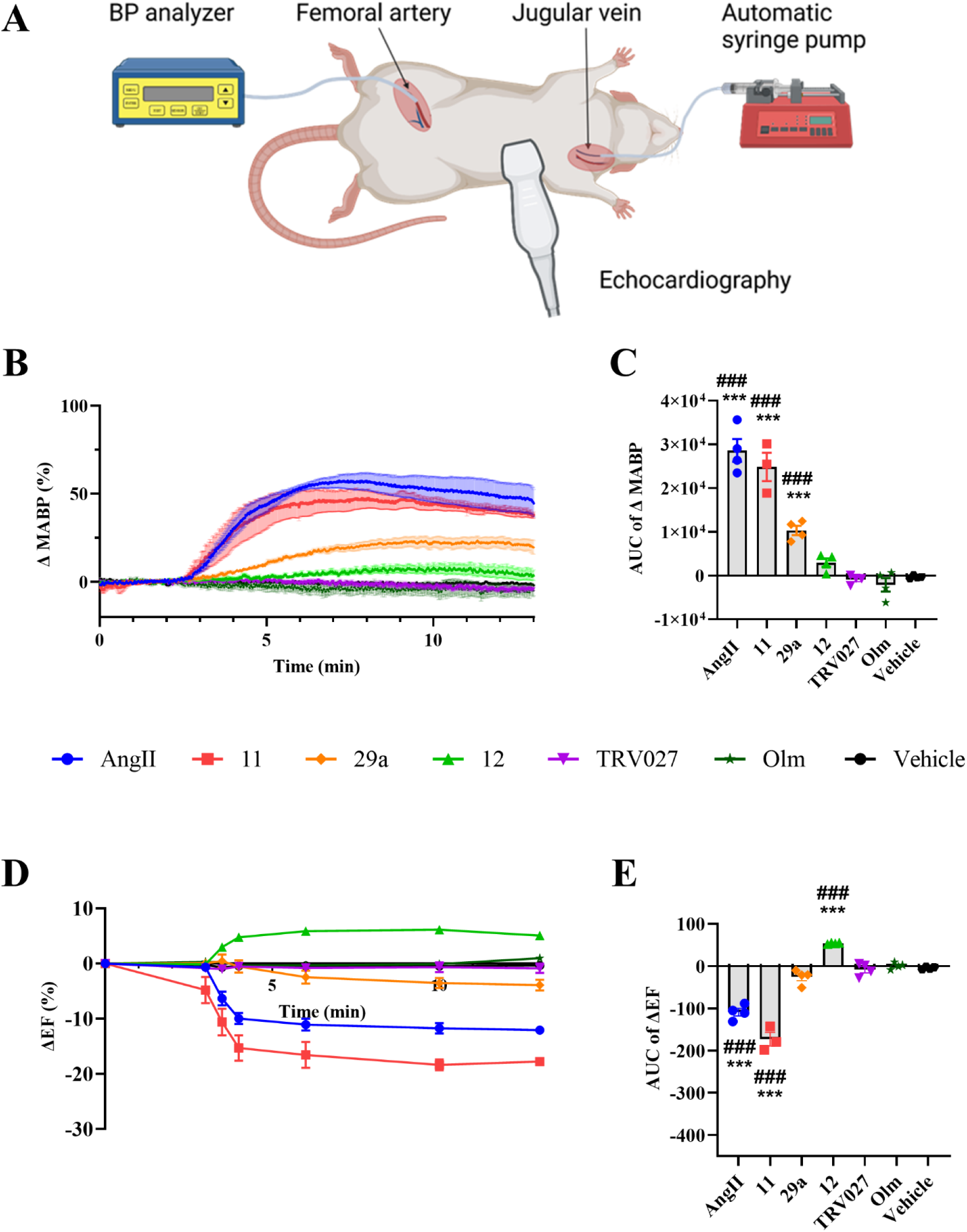
Simultaneous *in vivo* cardiac ultrasound imaging and blood pressure monitoring of adult Sprague-Dawley rats following constant administration of AT1R ligands. (**A**) The acquisition setup for each isoflurane-anesthetized rat includes a catheter in the right femoral artery connected to a TDX300 Micro-med transducer and BPA-100c blood pressure (BP) analyzer for continuous BP measurement, while drug administration is performed via a catheter inserted in the left jugular vein and linked to an automatic syringe pump set at a flow rate of 30 µl*min^−1^. Effect of AngII, selected analogs, and Olmesartan (Olm) on the changes in mean arterial blood pressure (ΔMABP; **B, C**) and ejection fraction (ΔEF; **D, E**), acquired simultaneously on isoflurane-anesthetized rats during intrajugular perfusion (0.96 nmol*kg^− 1^*min^−1^) of each ligand. The syringe pump is initiated at 0 s, and the catheter’s saline dead volume is flushed in approximately 180 s. Each data point (**B, D**) and bar (**C, E**; area under curve (AUC) of the corresponding administration) represent the mean ± SEM with n = 4 rats per group. Statistical analyses were performed with standard ANOVA followed by a Tukey’s multiple comparisons test. * p < 0.05, ** p < 0.01 and ***p<0.005 VS vehicle; # p < 0.05, ## p < 0.01 and ### p<0.005 VS TRV027 were considered significantly different.

Remarkably, **12** was the only β-arrestin-biased analog showing positive cardiac contractility at this equimolar dose (**Figures 5D-E**). These results suggest that a partial, low Gα_q_ activity, as seen with **12**, improves cardiac performance in physiological conditions. Therefore, we next challenged rats with a 10-fold increased dose (i.e., 9.6 nmol kg^−1^ min^−1^) of **12** and TRV07 to assess the impact of potentially increasing Gα_q_ activity or any other potential differences in the ligands’ cardiovascular effects under the same experimental conditions (**Figure 6**). The higher dose of analog **12** only marginally and transiently increased the mean arterial blood pressure, which returned to normotensive levels in less than 10 min (**Figure 6A**). This is likely due to a sealing pressor effect of **12** via the Gα_q_ pathway having reached its maximal intrinsic efficacy, combined with the rapid and efficient engagement of the autonomic baroreflex response, which helps maintain arterial blood pressure within a narrow range under physiological conditions.

**Figure 6.**
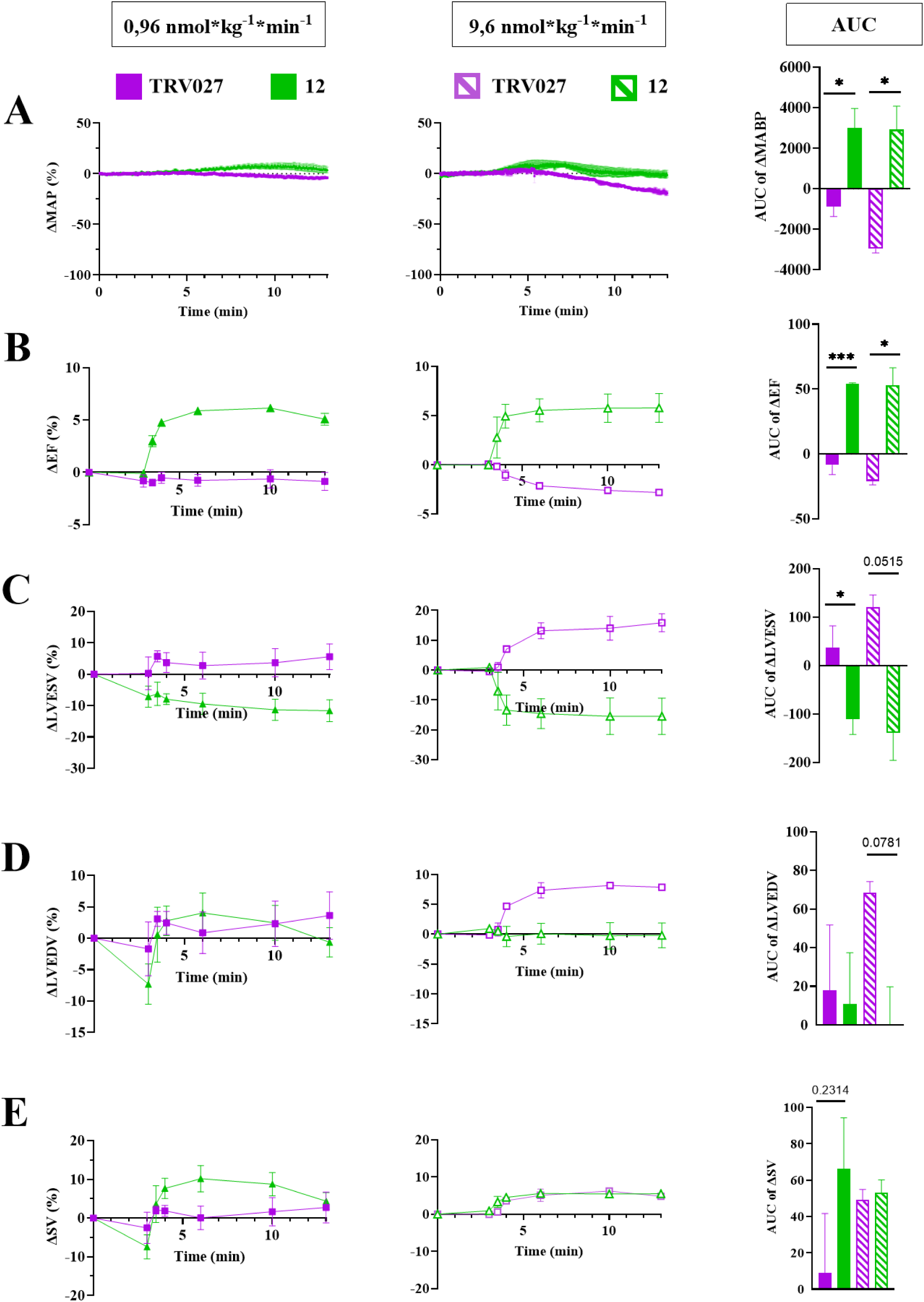
Comparison of the cardiovascular responses to constant administration of analog **12** and TRV027 at two different doses in adult Sprague-Dawley rats. Changes in mean arterial blood pressure (ΔMABP; **A**), ejection fraction (ΔEF; **B**), left ventricular end-systolic (ΔLVES; **C**) and end-diastolic (ΔLVEDV; **D**) volumes, and stroke volume (ΔSV; **E**) were measured from simultaneous *in vivo* cardiac blood pressure monitoring and ultrasound imaging. The acquisition setup for each isoflurane-anesthetized rat includes a catheter in the right femoral artery connected to a TDX300 Micro-med transducer and BPA-100c blood pressure (BP) analyzer for continuous BP measurement, while AngII analogs administration at two different doses (0.96 VS 9.6 nmol*kg^−1^*min^−1^; left and middle panels, respectively) is performed via a catheter inserted in the left jugular vein and linked to an automatic syringe pump set at a flow rate of 30 µl*min^−1^. The syringe pump is initiated at 0 s, and the catheter’s saline dead volume is flushed in approximately 180 s. Each data point (left and middle panels) and bar (area under the curve (AUC); right panels) of the corresponding administration represents the mean ± SEM with n = 2-4 rats per group. Statistical analyses were performed with an unpaired standard t-test. * p < 0.05, ** p < 0.01 and ***p<0.005.

Cardiac responses to **12** supported an increase in ejection fraction (**Figure 6B**), characterized by a reduction in left ventricular end-systolic volume without changes in end-diastolic volume (**Figures 6C–D**), and an associated increase in stroke volume (**Figure 6E**) observed at the lower dose (0.96 nmol kg^−1^ min^−1^). Unlike **12**, higher-dose TRV027 perfusion caused a progressive drop in arterial pressure (**Figure 6A**) that was not effectively compensated by the baroreflex. A similar decrease in mean arterial pressure was reported at doses as low as 0.096 nmol kg^−1^ min^−1^ in pentobarbital-anesthetized Sprague-Dawley rats, although escalating doses of TRV027 increased cardiac performance.^13^ Surprisingly, high-dose TRV027 in isoflurane-anesthetized rats increased stroke volume to a level similar to that seen with **12** (**Figure 6E**), but reduced ejection fraction (**Figure 6B**) and elevated both end-systolic and end-diastolic left ventricular volumes (**Figures 6C–D**). These divergent cardiac effects may stem from the differing anesthesia protocols used – sodium pentobarbital in the previous study vs. isoflurane in ours.^13^ Nonetheless, these findings highlight that ligands such as **12**, which retain β-arrestin signaling while exhibiting minimal peripheral vasopressor effects through optimized partial Gα_q/11_ activity, may help sustain or even enhance stroke volume and cardiac output.

### Molecular dynamics simulations reveal a putative deep allosteric binding pocket in AT1R

We conducted MD simulations to explore the structural dynamics of AT1R in association with varied selected ligands. Our focus was on how alterations at position 8 of AngII influence receptor conformation and dynamics and therefore cause diverse signaling results via Gα_q_ and β-arrestin. Specifically, we investigated how the structural extension of AngII (^EC^_50_ Gα_q_ 2.6 nM; E_max_ 100 %), attained through the addition of the Bzl group in Tyr(Bzl) (**12**), modulates Gα_q_ activation (^EC^_50_ Gα_q_ >10000 nM; E_max_ 11%) and how a subtle modification by introducing an *ortho*-cyano group (**23a**, (*o*-CN)Phe; ^EC^_50_ Gα_q_ >10000 nM; E_max_ 23%) significantly decreases Gα_q_ activation compared to AngII and analog **11** (^EC^_50_ Gα_q_ 1.1 nM; E_max_ 113%), which carry a Phe and a Tyr(Me) residue at position 8, respectively with minimal effect on β-arrestin recruitment (all ^EC^_50_ ^βarr2 ≤ 2.5 nM,^ E_max_ ≥ 89%). The MD simulation suggested that AngII produces a minor conformational shift in AT1R, which led to the formation of two crucial intramolecular key interactions (**Figures 7A-B**). Tyr^4^ interacted with Phe^8^ through an H-bond interaction, while it also engaged with His^6^ via an NH-π interaction. In the receptor’s binding site, AngII is stabilized through various hydrogen bonds and salt bridges with critical amino acids covering distinct structural regions, similar to Sar^1^-AngII. In AT1R’s N-terminal, residues Asp17^N-term^, Gln15^N-term^, and Arg2^N-term^, along with Arg167^4.64^, Phe182^ECL2^, and Gln187^ECL2^ found in ECL2, significantly contribute to ligand stabilization. In the transmembrane domains, Tyr87^2.64^, Ser105^3.32^, Lys199^5.42^, Asp263^6.51^, and Gln267^6.55^, along with Asp281^7.32^, further strengthen ligand binding and are probably key in mediating downstream signaling. In addition, we observed that the interaction with Arg272^ECL3^ was missing after the simulation compared to the initial frame, likely because of a minor conformational change in AngII’s N-terminal region (**Figure 7A**).

**Figure 7.**
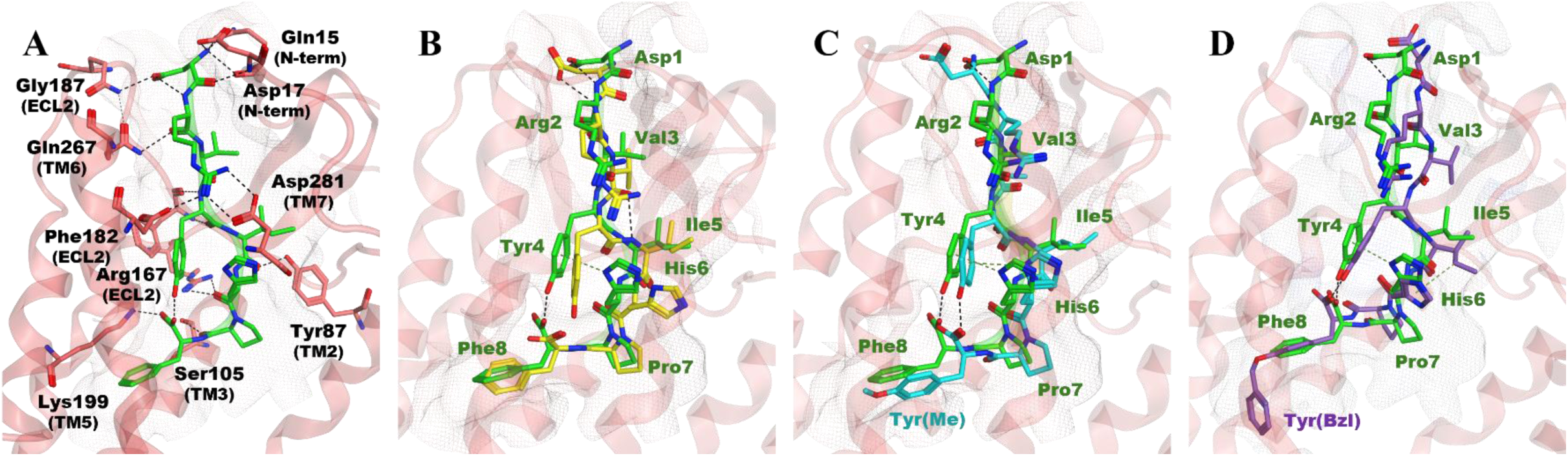
Extended AngII analogs can reach a cryptic allosteric site of the AT1R binding pocket. (**A**) Three-dimensional interactions of AngII within the binding site of AT1R. Structural comparison of the final frame of AngII (green) with its initial frame (yellow; **B**), **11** (cyan; **C**), and **12** (purple; **D**), all bound to AT1R. The distinct orientations of analogs’ key residues, including Phe⁸, Pro⁷, His⁶, and Tyr⁴, underscore structural differences that may contribute to variations in receptor activation and downstream signaling.

We identify key structural determinants within **11**, **12**, and **23a** in AT1R, which solely differ in their substitutions on the aromatic ring of Phe^8^ (AngII: Phe, **11**: Tyr(Me), **12**: Tyr(Bzl), and **23a**: Phe(*o*-CN)) and may govern their signaling profile (**Figure 7 and 8**). The MD trajectories reveal fundamental differences in each ligand’s ability to stabilize the key transmembrane helices that mediate G-protein activation and, crucially, show that opening of the orthosteric pocket exposes a putative cryptic allosteric site capable of accommodating the Bzl motif of compound **12** – consistent with our earlier predictions.^26^ Major differences arose from Trp253^6.48^ from the “toggle-switch” motif, which rotates by ±55°, allowing the Bzl to sink deeply into an allosteric cavity (**Figure 8E**). ^41^ This shifts Phe249^6.44^, thereby pushing TM6 outward, a known movement involved in GPCR activation.^6,42^ Other conformational changes, albeit more minor, also contribute to the opening of the cavity surrounding the Tyr(Bzl) side chain, which involve: Ser107^3.31^, Val108^3.32^, Leu112^3.36^, Tyr113^3.37^, and Val116^3.40^ in TM3; Lys199^5.42^ and Gly203^5.46^ in TM5; His256^6.51^, Gln257^6.52^ and Thr260^6.55^ in TM6; Met284^7.35^, Ile288^7.37^ and Ala291^7.40^ in TM7 (**Figure 8E**).

**Figure 8.**
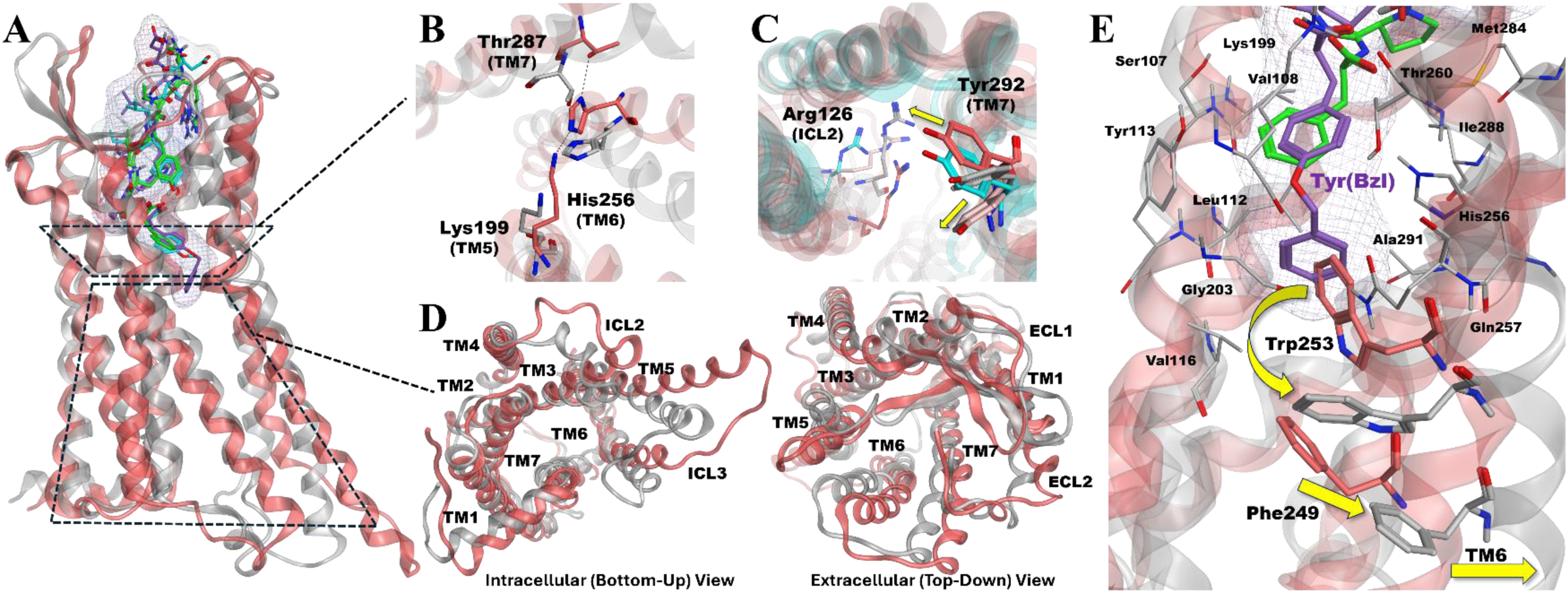
Structural comparison of analogs **11**, **12**, **23a**, and AngII in the AT1R binding pocket. (**A**) Superimposed receptor-ligand binding poses of **11** (cyan, pink receptor), **12** (purple, grey receptor), and AngII within AT1R, illustrating similar binding modes but distinct receptor conformations. The ligand-binding pocket is visualized with a mesh surface, showing how each ligand fits within the receptor. (**B**) Close-up view of the critical Lys^199^ (TM5) - His^256^ (TM6) - Thr^287^ (TM7) interaction network, which is preserved in **11** (pink receptor) but disrupted in analog **12** (grey receptor), indicating structural determinants for differential signaling outcomes. Additionally, **B**) highlights the different orientations of His^256^ in **11** and **12**, further contributing to their distinct receptor activation mechanisms. (**C**) Structural comparison of Tyr^292^ (TM7) and Arg126 (ICL2) in AngII (pink), **11** (cyan), **12** (grey), and **23a** (light pink). Tyr^292^ adopts distinct orientations toward TM3 or TM2, depending on the ligand, which may influence Gα_q_ versus β-arrestin signaling bias. Similarly, Arg^126^ (ICL2) exhibits ligand-specific positioning, impacting the signaling profile. (**D**) Intracellular and extracellular views of AT1R in the presence of **11** (pink ribon) and **12** (grey ribon). The intracellular view shows how ligand-induced conformational shifts propagate through the receptor core, affecting transmembrane helices. (**E**) Opening of a cryptic allosteric pocket induced by Tyr(Bzl) insertion in analog **12** (purple) compared to the Phe^8^ in AngII (green), facilitated by Trp^253^ rotation and Phe^249^ shift pushing TM6 outward. Thin sticks (grey) represent residues near Tyr(Bzl) with minimal conformational change, while thick sticks indicate those undergoing significant structural rearrangement.

In the **11**-bound complex, Lys199^5.42^, His256^6.51^, and Thr287^7.38^ form an interaction network between TM5, TM6, and TM7, preserving a conformation essential for Gα_q_ engagement (**Figure 8B**). Conversely, these stabilizing interactions are absent in **12**, resulting in a more compact receptor conformation that obstructs Gα_q_ coupling. This open-closed switch, seen in other GPCRs,^6^ is a critical element in AT1R signaling bias, with **11** promoting full Gα_q_ activation, and **12** favoring a β-arrestin-biased state by disrupting the TM5–TM6–TM7 network.^24^ We also noticed that the orientation of His256^6.51^ on AT1R seems to impact the signaling profile of the analogs. His256^6.51^’s orientation in both **23a** (not displayed) and **12** remained consistent with its positioning in AT1R-bound agonists (**Figure 8B**). This finding supports the idea that the unique dynamic of His256^6.51^ plays a pivotal role in hampering efficient Gα_q_ activation while biasing the receptor toward β-arrestin signaling. Notably, changing the orientation of His256^6.51^ impedes the creation of a key interaction network, leading to TM6 compaction and an inward rotation of TM7. This pattern is also observed in analogs such as **23a**, **12**, and some TRV derivatives, and was previously identified as crucial for ligand-dependent activation of AT1R.^24,43,44^

Our data, supported by a recent study,^25^ show that Tyr292^7.43^ adopts distinct orientations in AngII-bound AT1R compared with analogs **12** and **23a**, but more closely matches AngII’s orientation in **11** (**Figure 8C**). In the AngII- and **11**-bound receptors, Tyr292^7.43^ points toward TM2, stabilizing the canonical active conformation needed for efficient Gα_q_ signaling. Conversely, in **23a** and **12**, Tyr292^7.43^ pivots toward TM3, shifting the receptor into a state less favorable for Gα_q_ activation yet still suitable for β-arrestin engagement. Finally, Arg126^3.50^ of the DRY motif also adopted distinct conformations depending on the ligand (**Figure 8C**). In complexes with **12** and **23a**, Arg126^3.50^ shifted away from its canonical position, potentially weakening receptor stabilization and reducing its ability to engage Gα_q_ efficiently. This observation also agrees with the known importance of the conserved DRY motif for Gα_q_ protein activation by AT1R.^45^

These findings collectively suggest that modifications at position 8 of AngII reconfigure the conformational topology of AT1R. This reconfiguration exposes hidden allosteric sites and rearranges essential transmembrane interactions that gradually govern the signaling bias between Gα_q_ and β-arrestin.

## CONCLUSIONS

Although the eighth position of AngII has been recognized for its antagonistic properties on Gα_q/11_ responses via the AT1R, its role in governing biased signaling and *in vivo* cardiac regulation has been understudied.^8,19,21^ We have systematically explored the pharmacological properties of this position using diverse non-proteinogenic amino acid pharmacophores. This has allowed us to uncover nuanced variations that fine-tune AT1R signaling and bias profiles. Our molecular dynamic simulation analyses strongly suggest that modifications in AngII’s eighth position reshape the AT1R’s binding pocket, allowing to expose a cryptic allosteric cavity, and alter distinctly the receptor’s conformational landscape. This discovery is consistent with the receptor’s high conformational heterogeneity.^24,25^ We demonstrate that this newly uncovered cavity accommodates the large Bzl addition of analog **12**, pushing key transmembrane residues into orientations enabling β-arrestin engagement, while variably limiting Gα_q/11_ activation in a rheostat-like modulating fashion. Moreover, the open-closed switch between TM5, TM6, and TM7 is sensitive to modifications of the Phe^8^ aromatic ring. This feature highlights how even subtle yet distinct substitutions in AngII can modulate receptor variable conformations to govern distinct signaling and bias profiles. Our findings also reiterate the pivotal roles of Trp253^6.48^, His256^6.51^, Arg126^3.50^, and Tyr292^7.43^, as identified in prior studies,^4,26,41,46^ in the modulation of Gα_q/11_ and β-arrestin signaling. We also uncovered an allosteric site within the AT1R that not only improves ligand binding affinity and signaling efficiency but also specifically modulates biased signaling.

Our findings reveal that diverse β-arrestin-biased AngII analogs, which exhibit a spectrum of Gα_q/11_ signaling efficacies, correlate with distinct *in vivo* outcomes and can be rationally developed. This refines the oversimplified notion of the “binary bias” concept, where one pathway needs to be entirely suppressed while another remains fully active, as the optimal strategy for developing more effective drugs. Notably, the differential Gα_q/11_ signaling abilities among these analogs result in distinct hemodynamic and inotropic responses *in vivo*. For instance, compound **12** shows partial Gα_q/11_ activity, resulting in modest and transient vasopressor responses, yet engages β-arrestin potently and efficiently, causing pronounced and sustained inotropic effects. This *in vivo* response is contrasted with analogs like **11** and **29a**, which respectively display full and robust partial Gα_q/11_ activation, leading to more pronounced systemic vasopressor effects that may limit the inotropic response. The unique signaling properties and *in vivo* effects of compound **12** may offer therapeutic benefits in conditions such as refractory vasoplegic and cardiogenic shock, where standard inotropes and vasopressors like catecholamines turn ineffective due to receptor desensitization or resistance.^47^ In the broader context of acute HF management, compound **12** may complement the cardioprotective effects of β-arrestin-biased ligands like TRV027 by enhancing positive inotropy while maintaining hemodynamic stability without causing systemic hypotension, thereby broadening therapeutic options for difficult-to-treat cardiovascular conditions. Therefore, AngII analogs with partial Gα_q/11_ activity, like compound **12**, may provide sufficient cardiovascular support (i.e., increased cardiac contractility) while avoiding excessive vasoconstriction or hypotension, thus preserving or even enhancing tissue and organ perfusion.

While our findings propose new strategies for the modulation of AngII signaling via AT1R, limitations require consideration. First, though our results suggest that partial Gα_q/11_ agonism, coupled with full β-arrestin engagement seen in certain ligands such as **12**, might benefit inotropic function improvement and normal systemic vascular pressure preservation, the “threshold” of necessary Gα_q/11_ signaling level required for optimal clinical benefit remains to be established. Furthermore, it remains ambiguous as to how such partial Gα_q/11_ signaling could affect other key RAS-regulated responses like renal function, and to what extent analogs with partial Gα_q/11_ but full β-arrestin activity may be beneficial in pathophysiological conditions, frequently characterized by dysregulated RAAS and altered AT1R responses.^4^ Secondly, while our discoveries strongly support partial AT1R-Gα_q/11_ signaling’s role in the positive inotropic response, the role of the β-arrestin or other pathway-specific effects that may preferentially regulate cardiac contractility and inotropy induced by the analogs was not investigated. Consequently, these signaling pathways could differentially influence their *in vivo* effects on the vasculature versus the heart, potentially limiting or enhancing therapeutic efficacy and side-effect profiles. Despite these limitations, our findings establish a foundation for the design of new peptidic and non-peptidic AT1R ligands, whether orthosteric or small-molecule targeting this allosteric site, with tailored Gα_q/11_ versus β-arrestin signaling profiles, thereby expanding the AngII-based cardiovascular therapeutic portfolio.

## EXPERIMENTAL SECTION

### Reagents

Natural amino acids, HATU, and 2-Chlorotrityl chloride resin were purchased from Matrix Innovation (Québec, Canada). DIPEA was ordered from Chem-Impex (Illinois, USA). Piperidine was obtained from A&C (Québec, Canada). UAAs were obtained from Chem-Impex (Illinois, USA), Combi-blocks (California, USA), and Aaron chemicals (Jiangsu, China). Other solvents and reagents were purchased from Fischer Scientific (Ontario, Canada) and used as received.

### Solid Phase Synthesis

Peptides were synthesized on solid phase at a 0.05 mmol scale using Fmoc-based chemistry on Mettler Toledo MiniBlocks. The first amino acid was loaded onto the resin using a unimolecular nucleophilic substitution (SN1). An amino acid (1.5 eq, 0.075 mmol) was added to the 2-Cl resin (170 mg) with DIPEA (3 eq, 0.15 mmol, 27 µL) in a minimal amount of dichloromethane (DCM). The mixture was stirred overnight. Excess reagents were removed by washing twice with 2 mL DCM. The amino acid loading was quantified by measuring the UV absorbance of the dibenzofulvene–piperidine adduct resulting from Fmoc deprotection. The loading usually ranged from 0.25 - 0.35 mmol/g. Once the desired loading was achieved, the resin was capped with 2 mL of DCM/MeOH/DIPEA (17/2/1) for 15 min. The resin was washed five times with DCM and twice with ether. The subsequent amino acids were added to the sequence in two steps: 1) Fmoc deprotection, and 2) amide coupling. The resin was consistently rinsed with DMF-DMF-DCM-DCM-DMF between these two steps. Fmoc deprotection was achieved by treating the resin twice with 2 mL of 20% piperidine/DMF for 15 min. For the coupling steps, HATU (5 eq, 0.25 mmol, 97 mg) and the amino acid (5 eq, 0.25 mmol) were dissolved in 2 mL DMF, transferred to the resin, and then DIPEA (6 eq, 0.3 mmol, 54 μL) was added to initiate the coupling reaction. The reaction was allowed to run between 30 min and overnight, and the excess reagents were removed by filtration. The deprotection and coupling steps were repeated to synthesize the linear peptide. Final cleavage from the resin and simultaneous protecting groups removal was accomplished using a cocktail of trifluoroacetic acid (TFA)/triisopropylsilane (TIPS)/water (95:2.5:2.5). The cleavage reaction was run for 2 h. The mixture was filtered through a glass wool plug to remove solid particles, and the solution was slowly poured into 30 mL of pre-cooled ether (at 0°C) to precipitate the product. The crude peptide was isolated by centrifugation (3000 rpm, 15 min), and then dried under an air flow before being dissolved in acetonitrile (ACN)/water (40/60) for lyophilization.

Peptides were purified on an HPLC-MS system from Waters (Milford, USA), using a column XSELECTTM CSHTM Prep C18 (19 × 100 mm), packed with 5 µm particles, a UV detector 2998, an MS SQ Detector 2, a Sample Manager 2767, and a binary gradient module. The binary solvent system used was acetonitrile/water + 0.1% formic acid. The pure fragments, confirmed by UPLC-MS, were combined and lyophilized to form a white solid. The purity of these peptides was measured using a UPLC/MS system from Waters (Milford, USA). This system employed an Acquity UPLC® CSH™ C18 column (2.1 × 50 mm) packed with 1.7 µm particles and the gradient used was acetonitrile and water with 0.1% HCOOH (0→0.2 min: 5% acetonitrile; 0.2→1.5 min: 5%→95%; 1.5→1.8 min: 95%; 1.8→2.0 min: 95%→5%; 2.0→2.5 min: 5%). The purity of all peptides was found to be >95%.

Compounds 6, 23, 26, and 29 were purified on a HPLC-MS system from Waters (Milford, USA) using a column XSELECT™ CSH™ Prep C18 (19 × 100 mm) packed with 5 µm particles, a UV detector 2998, an MS SQ Detector 2, a Sample Manager 2767, and a binary gradient module. A binary solvent system (acetonitrile/water + 0.1% NH_4_OH) was utilized. Pure segments (endorsed by UPLC-MS) were consolidated and lyophilized with TFA drops to create a white solid. The purity of peptides was assessed via a UPLC/MS system from Waters (Milford, USA). An Acquity UPLC® CSH™ C18 column (2.1 x 50 mm) packed with 1.7 µm particles was used with the subsequent gradient: acetonitrile and water with 0.1% HCOOH (0→0.2 min: 1% acetonitrile; 0.2→1.5 min: 1%→50%; 1.5→1.8 min: 99%; 1.8→2.0 min: 99%→1%; 2.0→2.5 min: 1%).

### Cell culture and transfections

We utilized a clonal cell line HEK293, referred to hereafter as HEK293 cells, which has been previously described.^7,8^ As a gracious gift from A. Inoue (Tohoku University, Sendai, Miyagi, Japan), CRISPR Gα_q/11_ and Gα_12/13_ knockout cell lines (ΔG_q/11_ and ΔG_12/13_, respectively), were derived as previously explained.^48^ In brief, a CRISPR-Cas9 system was used to target either *GNAQ* and *GNA11* or *GNA12* and *GNA13* genes in HEK293T cells simultaneously to generate the Gα_q/11_ and Gα_12/13_ knockout cell lines. For the production of adenovirus, we utilized HEK293A cells. All cells were grown in DMEM supplemented with 10% FBS and Penicillin-Streptomycin-L-Glutamine (10000 IU penicillin, 10000 µg/mL Streptomycin, 200 mM L-Glutamine) at 37°C in T175 flasks under a 5% CO_2_ humidified chamber. Periodic testing was carried out for mycoplasma contamination (ABM™, Mycoplasma PCR Detection kit). For all biosensors used, cells underwent transfection with the PEI reagent, employing a PEI:DNA ratio of 3:1. For the Gα_q_ biosensor, HEK293 cells were transfected with the plasmids coding for Gα_q_-RLucII (120 ng), Gβ_1_ (3000 ng), Gγ_1_-GFP_10_ (3000 ng), human AT_1_ (3000 ng) and Salmon sperm DNA (SSD) (880 ng). For the β-arrestin2 recruitment assay, HEK293 cells were transfected with the plasmids coding for 120 ng β-arrestin-2-RLucII, 3000 ng AT_1_, 3000 ng rGFP-CAAX, and 3880 ng SSD. For the measurement of PKC, HEK293 cells were transfected with 3000 ng AT_1_, 600 ng PKC-c1b, and 6400 ng SSD. A mixture of 30 μg of PEI, which was incubated for 15 min, and 2×10^6^ cells was generated. 100 µL of the cell/DNA mixture was added to white 96-well plates and incubated at 37_°_C at 5% CO_2_ for 48 h. For the Rho assay, ΔG_12/13_ cells were seeded at a density of 9,000 cells per well in poly-ornithine-coated white 96-well plates and transfected the following day. In brief, 1 μg of total DNA in 100 μL of PBS was mixed with 100 μL of PBS containing 3 μL of 1 mg/mL PEI. For Rho signaling, 150 ng of AT1R DNA with 25 ng of PKN-RBD-RlucII ^8^, and 100 ng of rGFP-CAAX were used. Empty pcDNA was used to reach 1 µg of the total DNA amount in 100 μL of PBS. The DNA was then mixed with 100 μL of PBS containing 3 μL of 1 mg/mL PEI. After a 20-min incubation, the DNA/PEI complexes were dispensed into cells in 96-well plates (15 µL/well), and the experiment was carried out 48 h post-transfection.

### Adenovirus generation and VSMC transduction

The coding regions of the PKC-c1b sensor, PKN-RBD-RlucII, and rGFPcaax ^8^ were PCR-amplified and subcloned into the KpnI/XhoI sites of the pENTR3C vector (Invitrogen), respectively. These were then transferred into the adenoviral expression vector pAd/CMV/V5-DEST (Invitrogen), via gateway cloning (pAD-CMV-PKC-c1b, pAD-CMV-PKN-RBD-RLucII, and pAD-CMV-rGFPcaax). Viruses containing the sensor codes were generated following a previously outlined process.^49^ Briefly, HEK293A cells were seeded at a density of 1 × 10^6^ cells on 100-mm dishes. The following day, cells were transfected with 4 µg of PacI-digested pAd5-CMVsensors using Lipofectamine 2000 (Invitrogen). After 7–10 days post-transfection, mature virus particles were gleaned via the collection of cells and media, which then underwent multiple cycles of freezing and thawing, followed by centrifugation. The harvested virus was amplified by re-infection on 100-mm dishes of confluent HEK293A cells, and harvested 48-72 h later. Viral titers were determined by serial dilution of adenovirus stock and subsequent infection of HEK293A cells in 96-well plates. 36 h after infection, cells were stained using an anti-hexon antibody (ab8249; Abcam), and hexon-expressing cells were tallied via fluorescence microscopy. The virus was then stored at −80°C. After this, rat VSMCs (between the 13th to 15th passages) were seeded at a density of 20,000 cells in each well of white 96-well plates 24 h before transduction with the viral vector. The cells were then transduced with the PKC sensor-adenovirus (3 μL per well) or Rho sensor (1 μL each per well of PKN-RBD-RLucII-adnovirus and rGFP-caax adenovirus). After 24 h, the medium was replaced. 48 h post-viral transduction, the cells were thoroughly washed with Tyrode’s buffer. Finally, BRET assays were carried out as detailed below.

### BRET assays

On the day of the experiment, each well was washed with 80 µL of Tyrode’s buffer (140 mM NaCl, 2.7 mM KCl, 1 mM CaCl_2_, 12 mM NaHCO_3_, 5.6 mM D-glucose, 0.5 mM MgCl_2_, 0.37 mM NaH_2_PO_4_, 25 mM Hepes (pH 7.4)). This was followed by incubation for 1 h in Tyrode’s buffer at room temperature. For the Gα_q_, β-arrestin-2, and PKC biosensors used in HEK293 cells, coelenterazine 400A (at a final concentration of 5 μM) was added 10 min before stimulation. Cells were then stimulated with increasing doses of the indicated ligand, and BRET signals were measured using a Tristar2 (Berthold) microplate reader 10 min later. For the PKC and Rho biosensors in VSMCs, and the Rho biosensor in ΔG_12/13_ cells, cells were similarly stimulated with ligands for 1 min, 1 min 30 s, and 3 min, respectively, before BRET measurement. Coelenterazine 400a was added 3–4 min before initiating the measurements, at a final concentration of 2.5 μM. BRET signals were then measured using a Synergy2 (BioTek) microplate reader. The applied filter set was 410/80 nm and 515/30 nm, designed to detect the light emissions of RlucII (*Renilla luciferase*) (donor) and rGFP (acceptor), respectively.

### Data analysis of signaling profiles

BRET ratios were analyzed in GraphPad Prism 10 using nonlinear regression and then normalized as a percentage of the E_max_, with AngII serving as the reference ligand for obtaining the EC_50_ and E_max_.

### Binding assays

HEK293 cells, transiently expressing the human AT1R (3 million cells transfected with 4 μg of AT1R plasmid), were washed once with PBS and submitted to one freeze-thaw cycle. The fractured cells were gently scraped into the washing buffer (20 mM Tris-HCl, 5 mM MgCl_2_, 150 mM NaCl, pH 7.4), then centrifuged at 2500×g for 15 min at 4°C, and resuspended in binding buffer (washing buffer with 0.1% BSA, pH 7.4). Binding assays were carried out in 96-well plates. Dose displacement experiments were performed by incubating the fractured cells (20–40 μg of protein) at room temperature for 1 h with 0.8 nM ^125^I-AngII (2200 ci/mmol) as a tracer, alongside increasing concentrations of AngII or analogs. The incubation mixtures were filtered through binding plates (MultiScreen HTS, Merck Millipore, Cork, Ireland) and preincubated with the washing buffer for 1 h at 4°C to remove unbound ligands. The filtered membranes were washed thrice with 200 µL cold binding buffer (4°C). The filters with membranes were retrieved with punch tips (Filter Plates Punch Tips, Merck Millipore, Cork, Ireland). The γ emission was quantified using a 2470 Wizard^2^ automatic γ-counter from PerkinElmer (Waltham, USA) (82% efficiency). IC_50_ values, indicating the concentration of the tested ligand displacing 50% of the radiolabeled ligand from the receptor, were calculated from binding results using GraphPad Prism 10. A Kd of 0.8 nM for AngII was established by saturation binding experiments. The dissociation constant (*K*_i_ value) was determined from the IC_50_ using the Cheng-Prusoff equation and presented as a mean ± SEM of three independent experiments, each conducted in duplicate. VSMC cells were seeded at 80,000 cells per well in poly-ornithine-coated 24-wells. After 2 days, the cells were washed with ice-cold PBS and incubated with 0.12 nM 125I-AngII (2200 ci/mmol) in the binding buffer at 4°C, overnight. Nonspecific binding was measured in the presence of 1 µM AngII. The next day, cells were washed thrice with ice-cold PBS, solubilized in 0.5 mL of 0.2 M NaOH/0.05% SDS, and bound radioactivity was counted using a PerkinElmer Wizard 1470 automatic γ-counter. Protein quantities were evaluated using a Bradford assay. Receptor binding capacity was calculated using the equation: B/Bmax=[L]/(Kd+[L]) with a Kd of AngII determined at 0.8 nM. Bmax for AT1R expression in VSMC was projected at 50 fmol/mg.

### Plasma stability

In a 96-well plate, 6 μL of compound (1 mM aqueous solution, 10% DMSO) was incubated with 27 μL of plasma (from a male Sprague−Dawley rat) at 37°C for 0, 1, 2, 4, 7, and 24 h. The degradation was quenched by adding 140 μL of 1% formic acid in an acetonitrile-ethanol (1:1) solution containing 0.25 mM of *N,N*-dimethylbenzamide (used as an internal standard). This mixture was initially transferred to an Impact Protein Precipitation filter plate (Phenomenex, California), the filtrates from which were subsequently collected in a second 96-well UPLC plate. Both plates were centrifuged at 500 g for 10 min while maintaining a temperature of 4°C. The filtrates were then diluted with 30 μL of distilled water and processed for analysis using an Acquity UPLC-MS system class H (using an Acquity UPLC protein BEH C18 column that measured 2.1 mm × 50 mm and had 1.7 μm particles with pores 130 Å in width). The peptide half-life, derived from the degradation curve, was calculated using the exponential one-phase decay function in GraphPad Prism 9. Degradation was estimated by comparing the mass spectrum at each point with those at the zero-minute mark. The data were presented as a mean ± SEM of at least three unique experiments, each performed in singlicate.

### Simultaneous *in vivo* echocardiography and blood pressure measurements

Male Sprague-Dawley rats (230–350 g, 7–9 weeks of age) were initially anesthetized with 5% isoflurane (mixed with 100% oxygen at a flow rate of 2.5 L/min). The animal was placed in a supine position on a thermostatic pad set at 37°C, and anesthesia was maintained with 2% isoflurane, confirmed by the absence of reflexes to toe pinch. The right femoral artery was cannulated with a PE 50 catheter filled with heparinized saline (10 units/mL) and connected to a Micro-Med transducer (TDX-300) and a Micro-Med blood pressure analyzer (BPA-100c). The left jugular vein was also cannulated with a PE 50 catheter filled with heparinized saline (10 units/mL) and connected to a syringe operated by an automatic pump with a flow rate of 30 µL min-1. After a 5-min stabilization period, the catheter was flushed of the heparinized saline dead volume for 3 min, allowing for basal blood pressure measurement before compounds were perfused at two different dosages (i.e., 0.96 and 9.6 nmol kg^−1^ min^−1^, equivalent to 1 and 10 g kg^−1^ min^−1^, respectively, of AngII reference ligand) for an additional 10 min. Blood pressure parameters were continuously obtained throughout the experiment, and heart function parameters were simultaneously captured at various time points using a Vevo3100 echocardiography system equipped with an MX250 ultrasound probe (FUJIFILM VisualSonics). The ejection fraction was determined from a parasternal long-axis view of the left ventricle in B-mode imaging and analyzed using Vevo LAB 5.7.1 (FUJIFILM VisualSonics). Data were presented as the mean ± SEM of four different rats per group. Statistical analyses were conducted with standard analysis of variance (ANOVA) followed by Tukey’s multiple comparisons test. Detection and elimination of outliers from datasets were based on the interquartile range method, using a multiplier of 1.5 for determining the threshold.

### Computational modeling

#### Molecular Modeling - System Preparation

MD simulations were conducted to examine the structural dynamics of AT1R when bound to different ligands. The Protein Data Bank provided the original receptor structure (PDB ID: 7F6G, https://www.rcsb.org/structure/7F6G), corresponding to an active-state AT1R configuration. The Molecular Operating Environment (MOE) modeled any missing side chains and loops to ensure the structure was complete.^50^ Using AngII as a template, ligand structures were built in MOE, and then the energy was minimized and parameterized using the CHARMM General Force Field (CGenFF). In both the upper and lower leaflets, the receptor was embedded in a symmetric lipid bilayer with an evenly distributed 650 lipids per leaflet, using CHARMM-GUI.^51^ The orientation of the complex was determined using the PPM 2.0 tool, and the complex-lipid system was solvated with the assistance of TIP3P explicit water molecules. The addition of 150 mM NaCl neutralized the system. The final simulation box dimensions were set at 201.02 Å × 201.02 Å × 198.94 Å, with a complete system size of 778,675 atoms. The CHARMM36m force field was applied to proteins and lipids, while the CHARMM General Force Field (CGenFF) parameterized the ligands.^52^ The Verlet cutoff scheme was used for nonbonded interactions, with a 1.2 nm cutoff applied to both van der Waals and Coulomb interactions. The Particle Mesh Ewald (PME) method was engaged to manage long-range electrostatics.^53^

#### Molecular Modeling - Energy Minimization and Equilibration

The system underwent initial energy minimization using the steepest descent algorithm to alleviate steric clashes and relax the structure. Positional restraints were incorporated for the backbone (4000 kJ mol⁻¹ nm⁻²), side chains (2000 kJ mol⁻¹ nm⁻²), and lipids (1000 kJ mol⁻¹ nm⁻²), with dihedral restraints applied at a force constant of 1000 kJ mol⁻¹ nm⁻². The system was minimized until the strongest force affecting any atom went below 1000 kJ mol⁻¹ nm⁻¹. Following minimization, six equilibration steps were carried out to gradually alleviate the system while preserving positional and dihedral restraints. The first couple of equilibration steps took place in the NVT ensemble, using a 1 fs integration time step and v-rescale thermostat holding the temperature at 303.15 K. Amid these steps, the backbone, side chain, and lipid restraints remained at 4000, 2000, and 1000 kJ mol⁻¹ nm⁻² appropriately, then progressively reduced in a stepwise mode. The subsequent equilibration steps occurred in the NPT ensemble with a C-rescale barostat, using semi-isotropic pressure coupling at 1 bar and a compressibility of 4.5 × 10⁻⁵ bar⁻¹. The integration time step rose to 2 fs, and the restraints were gradually lessened to 50 kJ mol⁻¹ nm⁻² for the backbone, with side chains and lipids entirely unrestrained in the last equilibration stage. The LINCS algorithm was implemented to constrain all bonds, enabling a stable equilibration process. The PME method was executed for long-range electrostatics, and a Verlet cutoff scheme was used with 1.2 nm cutoffs for van der Waals and Coulomb interactions. After equilibration, the system was prepped for the production MD simulation, where all restraints were lifted, and the simulation was carried out under NPT ensemble conditions for 500 ns using a 2 fs time step. The final equilibrated structure offered a stable launch point for production simulations.

#### Molecular Modeling - Production MD Simulations

Production MD simulations were performed using GROMACS 2023 for 100 ns, under NPT conditions,^54^ with a 2 fs integration time step. All bonds involving hydrogen atoms were constrained using the LINCS algorithm, which allowed for an increased time step. The system’s temperature and pressure were controlled using the v-rescale thermostat and C-rescale barostat, respectively. Nonbonded interactions were handled with a 1.2 nm cutoff, applying the force-switch modifier for van der Waals interactions. For analysis, trajectory snapshots were recorded every 50 ps.

#### Molecular Modeling - Trajectory Analysis

The post-simulation analysis was conducted using MOE and GROMACS. All AT1R residues were annotated according to the numbering scheme proposed by Ballesteros and Weinstein.^55^

#### Corresponding compound numbers related to future publications

1= HAD001 ; **2**= HAD002 ; **3**= HAD003 ; **4**= HAD004 ; **5**= HAD005 ; **6**= HAD006 ; **7**= HAD007 ; **8**= HAD008 ; **9**= HAD009 ; **10**= HAD118 ; **11**=HAD011 ; **12**=HAD012 ; **13**= HAD126 ; **14**= HAD157 ; **15**= HAD155 ; **16**= HAD013 ; **17**= HAD125 ; **18**= HAD159 ; **19**= HAD158 ; **20**= HAD156 ; **21**= HAD154 ; **22a**= HAD160 ; **22b**= HAD161 ; **22c**= HAD117 ; **23a**= HAD149; **23b**= HAD150 ; **23c**= HAD151 ; **24a**= HAD152 ; **24b**= HAD153 ; **24c**= HAD127 ; **25a**= HAD128 ; **25b**= HAD129 ; **25c**= HAD130 ; **26a**= HAD119 ; **26b**= HAD120 ; **26c**= HAD121 ; **27a**= HAD162 ; **27b**= HAD163 ; **27c**= HAD124 ; **28a**= HAD164 ; **28b**= HAD165 ; **28c**= HAD123 ; **29a**= HAD166 ; **29b**= HAD167 ; **29c**= HAD122 ; **30a**= HAD168 ; **30b**= HAD169 ; **30c**= HAD010

## ASSOCIATED CONTENT

### Supporting Information

HRMS and analytical UPLC-MS compound spectra are described in this article’s Supporting Information section.

## AUTHOR INFORMATION

### Corresponding Authors

**Pierre-Luc Boudreault**

Département de Pharmacologie-Physiologie, Institut de Pharmacology de Sherbrooke, Centre de Recherche du CHUS, Université de Sherbrooke, Sherbrooke, J1H 5N4, Québec, Canada.

**Stéphane A. Laporte**

Department of Medicine, Department of Pharmacology and Therapeutics, Research Institute of the McGill University Health Center (RI-MUHC), McGill University, Montréal, H4A 3J1, Québec, Canada.

**Mannix Auger-Messier**

Département de Médecine – Service de cardiologie, Institut de Pharmacologie de Sherbrooke, Centre de Recherche du CHUS, Université de Sherbrooke, Sherbrooke, J1H 5N4, Québec, Canada.

## ACKNOWLEDGMENTS

This work was supported by the Canadian Institutes of Health Research – CIHR (PJT-173504) and the Éric Marsault Research Chair in Medicinal Chemistry and Peptidomimetics (P-L.B), Université de Sherbrooke, Québec, Canada; J.M. received the Louis Gendron and Mylène Côté Excellence Scholarship from Fondation de l’Université de Sherbrooke (FUS) and a Canada Graduate Scholarships – Master’s (CGS M) award from the Canadian Institutes of Health Research (CIHR). M-F.R. was supported by a CIHR fellowship (699305). M.A.-M. was supported by a FRQS–Senior research scholarship-career award (313286). The authors have no other relevant affiliations or financial involvement with any organization or entity with a financial interest in or financial conflict with the subject matter or materials discussed in the manuscript apart from those disclosed.

## ABBREVIATIONS

AngII: Angiotensin II
AT1R: Angiotensin II Type 1 Receptor
BRET: Bioluminescence Resonance Energy Transfer
BSA: Bovine Serum Albumin
DBC: Deep Blue Coelenterazine
DCM: Dichloromethane
DIPEA: *N,N*-diisopropylethylamine
DMEM: Dulbecco’s Modified Eagle Medium
DMF: *N,N*-dimethylformamide
EC_50_: Half-maximal effective concentration
E_max_: maximum efficacy
FBS: Fetal Bovine Serum
GPCR: G Protein-Coupled Receptor
HATU: *O*-(7-azabenzotriazol-1-yl)-1,1,3,3-tetramethyluronium hexafluorophosphate
HEK: Human embryonic kidney
HRMS: High-Resolution Mass Spectrometry
Ovn: overnight
PBS: Phosphate-Buffered Saline
PEI: Polyethylenimine
PKC: Protein Kinase C
RAAS: renin-angiotensin–aldosterone system
rGFP: Green Fluorescent Protein Recombinant
RlucII: *Renilla luciferase*
SAR: Structure-Activity Relationship
SPPS: Solid-Phase Peptide Synthesis
SSD: Single-stranded DNA
UPLC: Ultra-Performance Liquid Chromatography

