## Supporting information for "Tunable Bias Signaling of the Angiotensin II Type 1 Receptor for Inotropy via C-Terminal Peptide Modifications and Allosteric Site Targeting"

**Supp Figure 1.** Dose-response curves for G$\alpha_q$ signaling by AT1R for all AngII analogs.

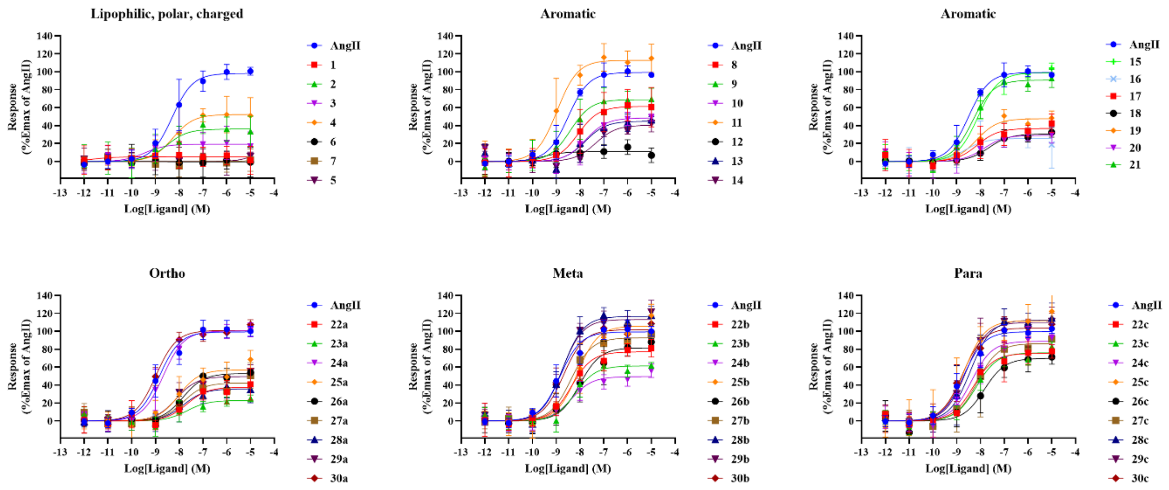


**Supp Figure 2.** Dose-response curves for β-arrestin engagement to AT1R for all AngII analogs.

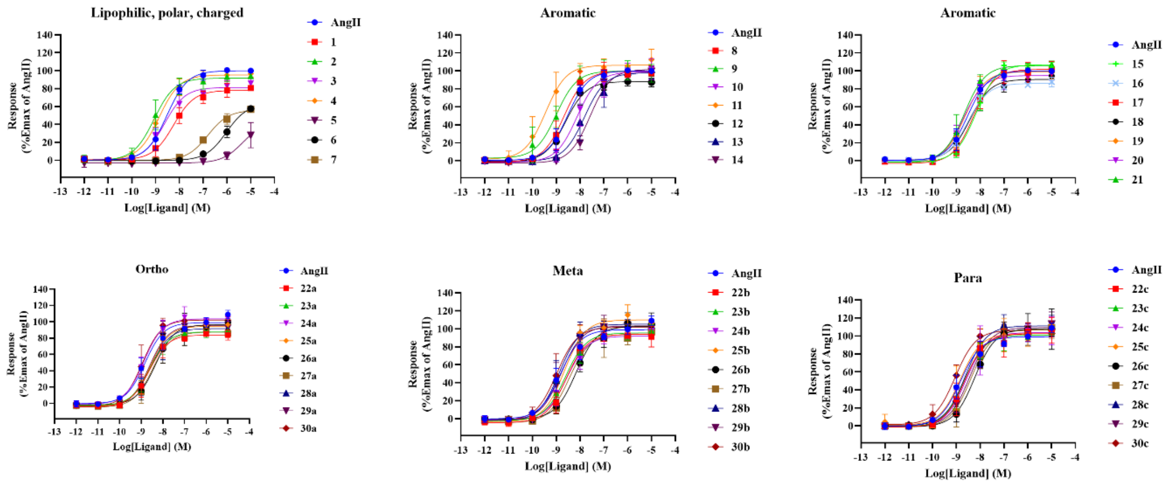


**Supp Table 1.** Plasmatic half-life of AngII analogs.

| **Compound** | **Peptide sequence** | **Half-life $t_{1/2}$(min) (±SEM)** |
|---|---|---|
| AngII | D-R-V-Y-I-H-P-F | 24±3 |
| 11 | D-R-V-Y-I-H-P-**Tyr(Me)** | 33±4 |
| 29a | D-R-V-Y-I-H-P-**(*o*-Cl)F** | 32±3 |
| 12 | D-R-V-Y-I-H-P-**Tyr(Bzl)** | 32±3 |
| TRV027 | **Sar** R-V-Y-I-H-P-**a** | 964±72 |

### Compound characterization (HRMS, purity)

| Compounds | Peptide sequence [a] | Mol formula | Purity (%) | Theoretical m/z | | HRMS |
|---|---|---|---|---|---|---|
| 1 | D-R-V-Y-I-H-P-A | $C_{44}H_{67}N_{13}O_{12}$ | 98,41 | $[M+2H]^{2+}$ | 485.7589 | 485.7612 |
| 2 | D-R-V-Y-I-H-P-Nle | $C_{47}H_{73}N_{13}O_{12}$ | 100 | $[M+2H]^{2+}$ | 506.7824 | 506.7822 |
| 3 | D-R-V-Y-I-H-P-Tle | $C_{47}H_{73}N_{13}O_{12}$ | 98.88 | $[M+2H]^{2+}$ | 506.7824 | 506.7833 |
| 4 | D-R-V-Y-I-H-P-Cha | $C_{50}H_{77}N_{13}O_{12}$ | 100 | $[M+2H]^{2+}$ | 526.7980 | 526.7984 |
| 5 | D-R-V-Y-I-H-P-Orn | $C_{46}H_{72}N_{14}O_{12}$ | 96.97 | $[M+2H]^{2+}$ | 507.2800 | 507.2814 |
| 6 | D-R-V-Y-I-H-P-E | $C_{46}H_{69}N_{13}O_{14}$ | 99.23 | $[M+2H]^{2+}$ | 514.7616 | 514.7616 |
| 7 | D-R-V-Y-I-H-P-Q | $C_{46}H_{70}N_{14}O_{13}$ | 98.82 | $[M+2H]^{2+}$ | 514.2696 | 514.2713 |
| 8 | D-R-V-Y-I-H-P-1Nal | $C_{54}H_{73}N_{13}O_{12}$ | 98.63 | $[M+2H]^{2+}$ | 548.7824 | 548.7826 |
| 9 | D-R-V-Y-I-H-P-2Nal | $C_{54}H_{73}N_{13}O_{12}$ | 100 | $[M+2H]^{2+}$ | 548.7824 | 548.7818 |
| 10 | D-R-V-Y-I-H-P-Tyr(All) | $C_{53}H_{75}N_{13}O_{13}$ | 100 | $[M+2H]^{2+}$ | 551.7876 | 551.7890 |
| 11 | D-R-V-Y-I-H-P-Tyr(Me) | $C_{51}H_{73}N_{13}O_{13}$ | 100 | $[M+2H]^{2+}$ | 538.7798 | 538.7808 |
| 12 | D-R-V-Y-I-H-P-Tyr(Bzl) | $C_{57}H_{77}N_{13}O_{13}$ | 100 | $[M+2H]^{2+}$ | 576.7955 | 576.7957 |
| 13 | D-R-V-Y-I-H-P-Bip | $C_{56}H_{75}N_{13}O_{12}$ | 100 | $[M+2H]^{2+}$ | 561.7902 | 561.7897 |
| 14 | D-R-V-Y-I-H-P-*p*-tBuPhe | $C_{54}H_{79}N_{13}O_{12}$ | 100 | $[M+2H]^{2+}$ | 551.8058 | 551.8072 |
| 15 | D-R-V-Y-I-H-P-Dip | $C_{56}H_{75}N_{13}O_{12}$ | 100 | $[M+2H]^{2+}$ | 561.7902 | 561.7911 |
| 16 | D-R-V-Y-I-H-P-Ser(Bzl) | $C_{51}H_{73}N_{13}O_{13}$ | 100 | $[M+2H]^{2+}$ | 538.7798 | 538.7804 |
| 17 | D-R-V-Y-I-H-P-4Pyridyl | $C_{49}H_{70}N_{14}O_{12}$ | 100 | $[M+2H]^{2+}$ | 524.2722 | 524.2715 |
| 18 | D-R-V-Y-I-H-P-W | $C_{52}H_{72}N_{14}O_{12}$ | 100 | $[M+2H]^{2+}$ | 543.2800 | 543.2809 |
| 19 | D-R-V-Y-I-H-P-4ThzAla | $C_{47}H_{68}N_{14}O_{12}S$ | 95.23 | $[M+2H]^{2+}$ | 527.2504 | 527.2513 |
| 20 | D-R-V-Y-I-H-P-2Pyridyl | $C_{49}H_{70}N_{14}O_{12}$ | 100 | $[M+2H]^{2+}$ | 524.2722 | 524.2736 |
| 21 | D-R-V-Y-I-H-P-3Pyridyl | $C_{49}H_{70}N_{14}O_{12}$ | 100 | $[M+2H]^{2+}$ | 524.2722 | 524.2727 |
| 22a | D-R-V-Y-I-H-P-F(2$NO_2$) | $C_{50}H_{70}N_{14}O_{14}$ | 100 | $[M+2H]^{2+}$ | 546.2671 | 546.2676 |
| 22b | D-R-V-Y-I-H-P-F(3$NO_2$) | $C_{50}H_{70}N_{14}O_{14}$ | 100 | $[M+2H]^{2+}$ | 546.2671 | 546.2684 |
| 22c | D-R-V-Y-I-H-P-F(4$NO_2$) | $C_{50}H_{70}N_{14}O_{14}$ | 98.6 | $[M+2H]^{2+}$ | 546.2671 | 546.2679 |
| 23a | D-R-V-Y-I-H-P-F(2CN) | $C_{51}H_{70}N_{14}O_{12}$ | 96.93 | $[M+2H]^{2+}$ | 536.2722 | 536.2732 |
| 23b | D-R-V-Y-I-H-P-F(3CN) | $C_{51}H_{70}N_{14}O_{12}$ | 100 | $[M+2H]^{2+}$ | 536.2722 | 536.2734 |
| 23c | D-R-V-Y-I-H-P-F(4CN) | $C_{51}H_{70}N_{14}O_{12}$ | 100 | $[M+2H]^{2+}$ | 536.2722 | 536.2735 |
| 24a | D-R-V-Y-I-H-P-F(2OH) | $C_{50}H_{71}N_{13}O_{13}$ | 98.81 | $[M+2H]^{2+}$ | 531.7720 | 531.7733 |
| 24b | D-R-V-Y-I-H-P-F(3OH) | $C_{50}H_{71}N_{13}O_{13}$ | 100 | $[M+2H]^{2+}$ | 531.7720 | 531.7741 |
| 24c | D-R-V-Y-I-H-P-F(4OH) | $C_{50}H_{71}N_{13}O_{13}$ | 98.32 | $[M+2H]^{2+}$ | 531.7720 | 531.7721 |
| 25a | D-R-V-Y-I-H-P-F(2Me) | $C_{51}H_{73}N_{13}O_{12}$ | 98.16 | $[M+2H]^{2+}$ | 530.7824 | 530.7833 |
| 25b | D-R-V-Y-I-H-P-F(3Me) | $C_{51}H_{73}N_{13}O_{12}$ | 99.4 | $[M+2H]^{2+}$ | 530.7824 | 530.7836 |
| 25c | D-R-V-Y-I-H-P-F(4Me) | $C_{51}H_{73}N_{13}O_{12}$ | 100 | $[M+2H]^{2+}$ | 530.7824 | 530.7826 |
| 26a | D-R-V-Y-I-H-P-F(2$CF_3$) | $C_{51}H_{70}F_3N_{13}O_{12}$ | 100 | $[M+2H]^{2+}$ | 557.7682 | 557.7683 |
| 26b | D-R-V-Y-I-H-P-F(3$CF_3$) | $C_{51}H_{70}F_3N_{13}O_{12}$ | 98.68 | $[M+2H]^{2+}$ | 557.7682 | 557.7668 |
| 26c | D-R-V-Y-I-H-P-F(4$CF_3$) | $C_{51}H_{70}F_3N_{13}O_{12}$ | 100 | $[M+2H]^{2+}$ | 557.7682 | 557.7680 |
| 27a | D-R-V-Y-I-H-P-F(2I) | $C_{50}H_{70}IN_{13}O_{12}$ | 98.81 | $[M+2H]^{2+}$ | 586.7229 | 586.7243 |
| 27b | D-R-V-Y-I-H-P-F(3I) | $C_{50}H_{70}IN_{13}O_{12}$ | 100 | $[M+2H]^{2+}$ | 586.7229 | 586.7242 |
| 27c | D-R-V-Y-I-H-P-F(4I) | $C_{50}H_{70}IN_{13}O_{12}$ | 98.32 | $[M+2H]^{2+}$ | 586.7229 | 586.7246 |
| 28a | D-R-V-Y-I-H-P-F(2Br) | $C_{50}H_{70}BrN_{13}O_{12}$ | 100 | $[M+2H]^{2+}$ | 562.7298 | 562.7312 |
| 28b | D-R-V-Y-I-H-P-F(3Br) | $C_{50}H_{70}BrN_{13}O_{12}$ | 99.37 | $[M+2H]^{2+}$ | 562.7298 | 562.7312 |
| 28c | D-R-V-Y-I-H-P-F(4Br) | $C_{50}H_{70}BrN_{13}O_{12}$ | 100 | $[M+2H]^{2+}$ | 562.7298 | 562.7318 |
| 29a | D-R-V-Y-I-H-P-F(2Cl) | $C_{50}H_{70}ClN_{13}O_{12}$ | 100 | $[M+2H]^{2+}$ | 540.7550 | 540.7566 |
| 29b | D-R-V-Y-I-H-P-F(3Cl) | $C_{50}H_{70}ClN_{13}O_{12}$ | 100 | $[M+2H]^{2+}$ | 540.7550 | 540.7562 |
| 29c | D-R-V-Y-I-H-P-F(4Cl) | $C_{50}H_{70}ClN_{13}O_{12}$ | 100 | $[M+2H]^{2+}$ | 540.7550 | 540.7568 |
| 30a | D-R-V-Y-I-H-P-F(2F) | $C_{50}H_{70}FN_{13}O_{12}$ | 100 | $[M+2H]^{2+}$ | 532.7698 | 532.7704 |
| 30b | D-R-V-Y-I-H-P-F(3F) | $C_{50}H_{70}FN_{13}O_{12}$ | 100 | $[M+2H]^{2+}$ | 532.7698 | 532.7702 |
| 30c | D-R-V-Y-I-H-P-F(4F) | $C_{50}H_{70}FN_{13}O_{12}$ | 100 | $[M+2H]^{2+}$ | 532.7698 | 532.7700 |

### UPLC-MS spectra

Compound 1: D-R-V-Y-I-H-P-A

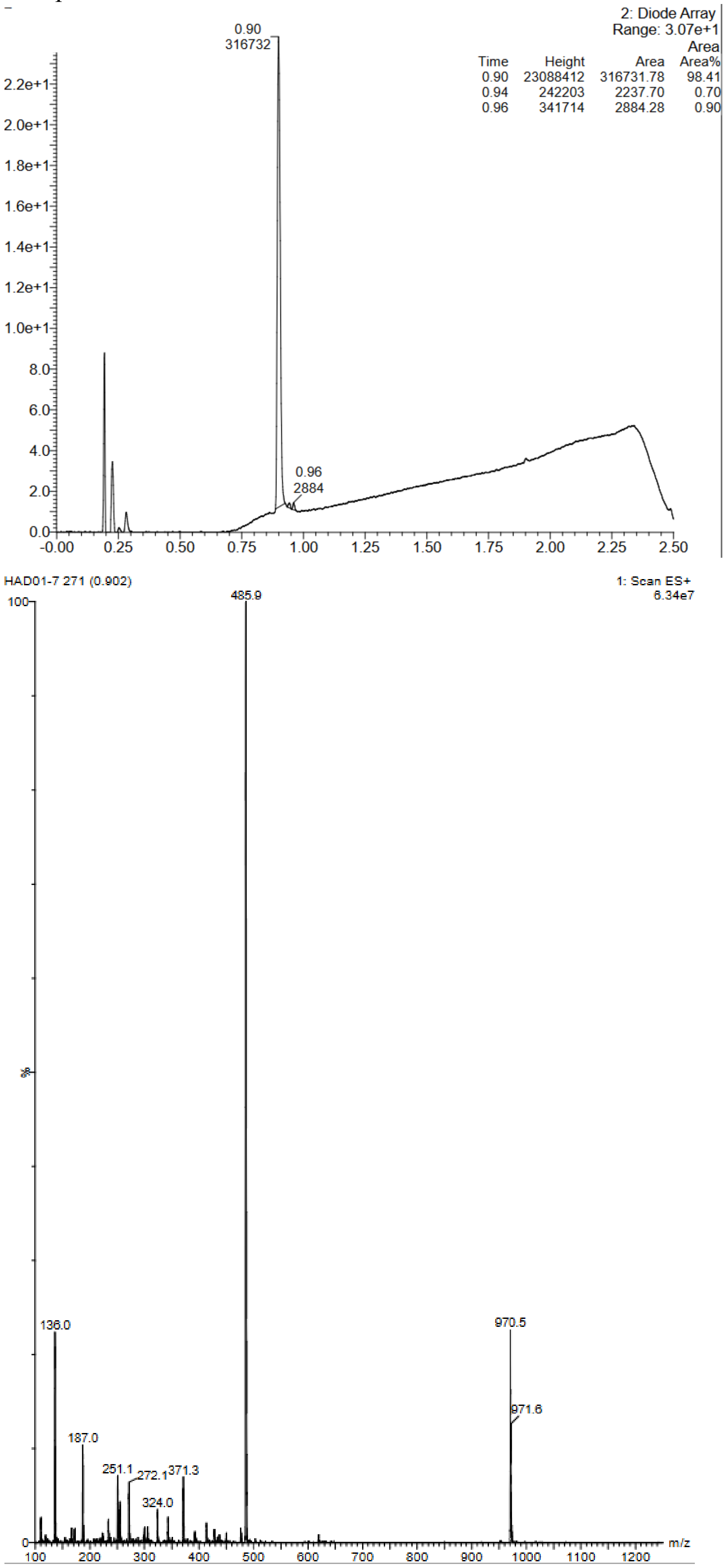

Compound 2: D-R-V-Y-I-H-P-Nle

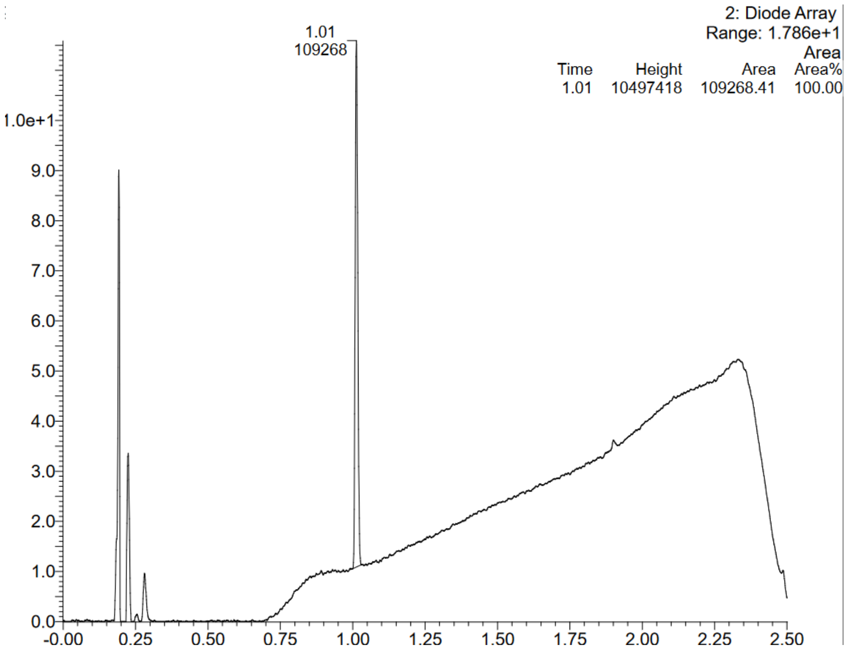


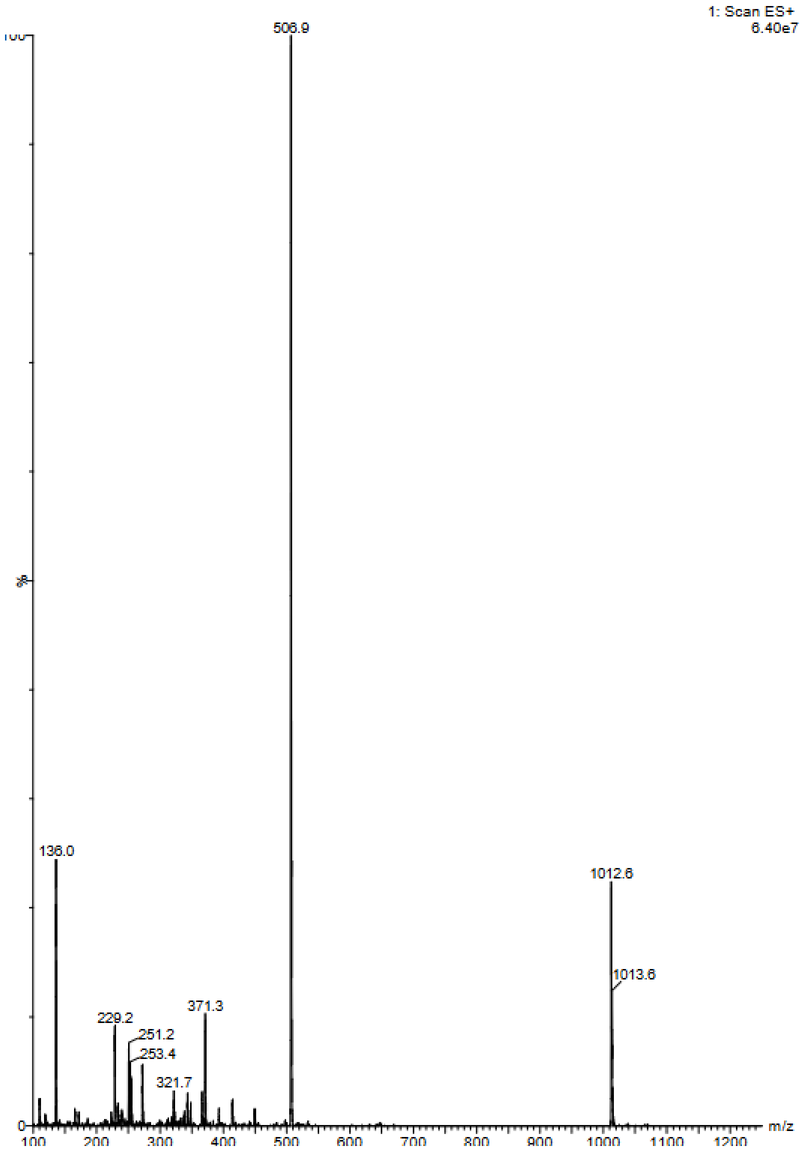

Compound 3: D-R-V-Y-I-H-P-Tle

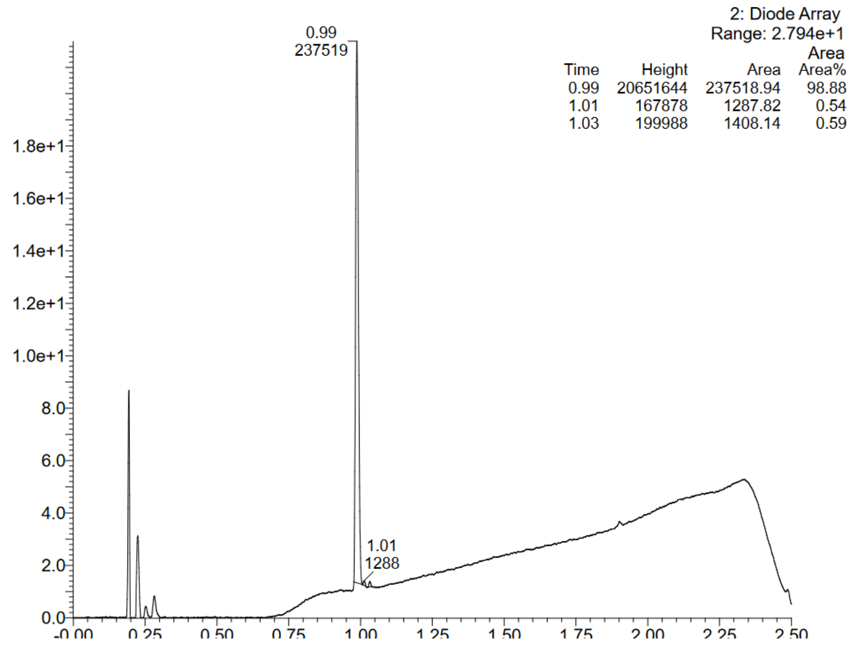


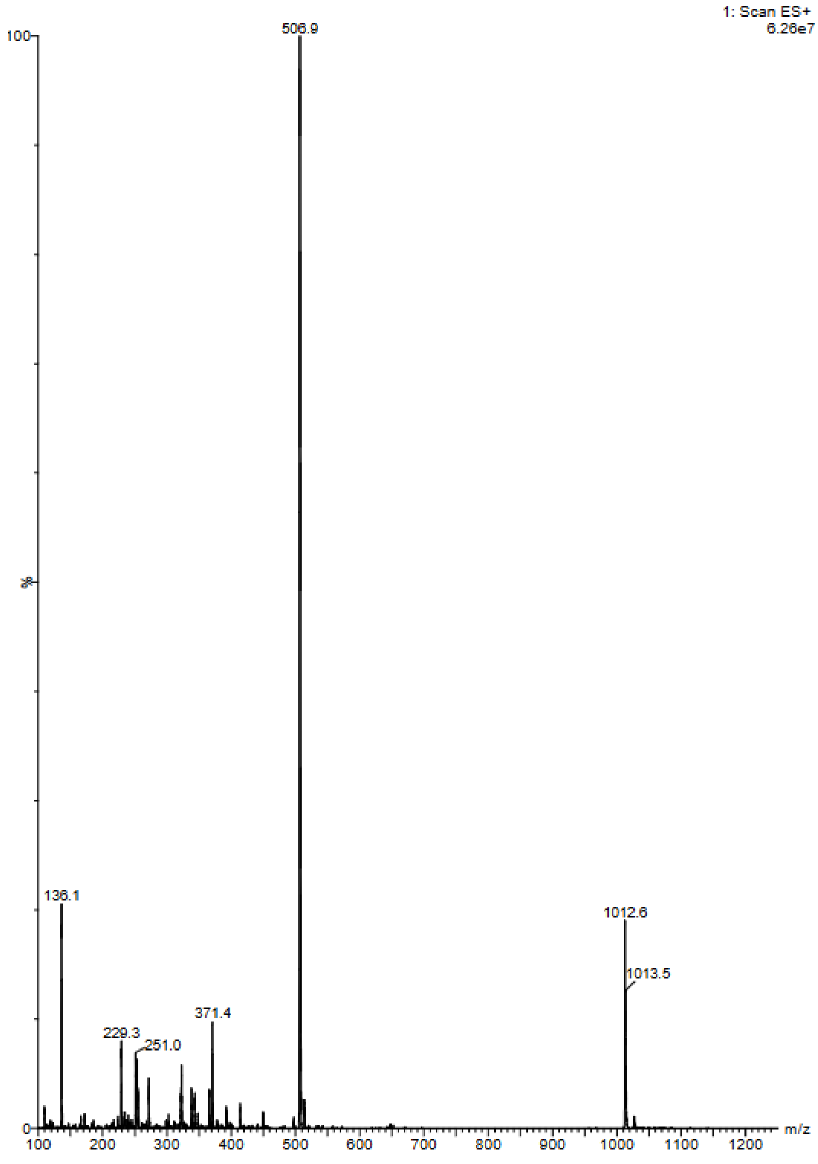

Compound 4:    D-R-V-Y-I-H-P-Cha

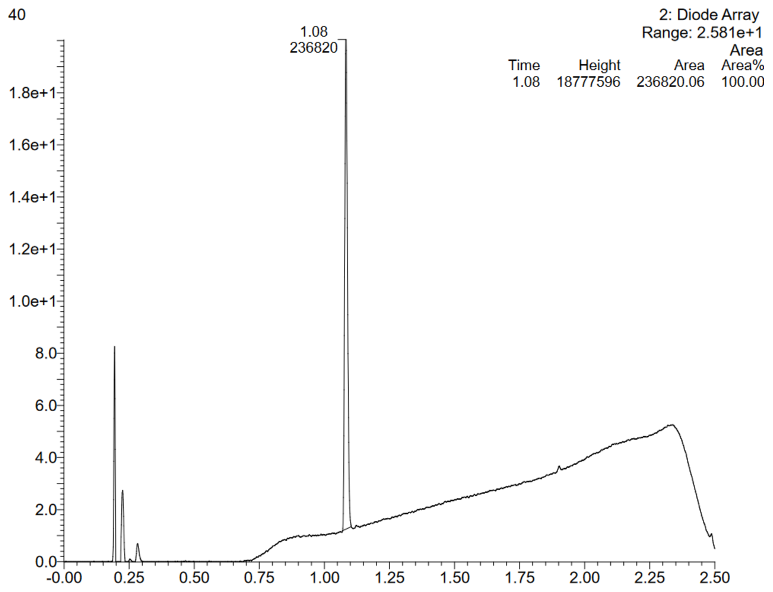


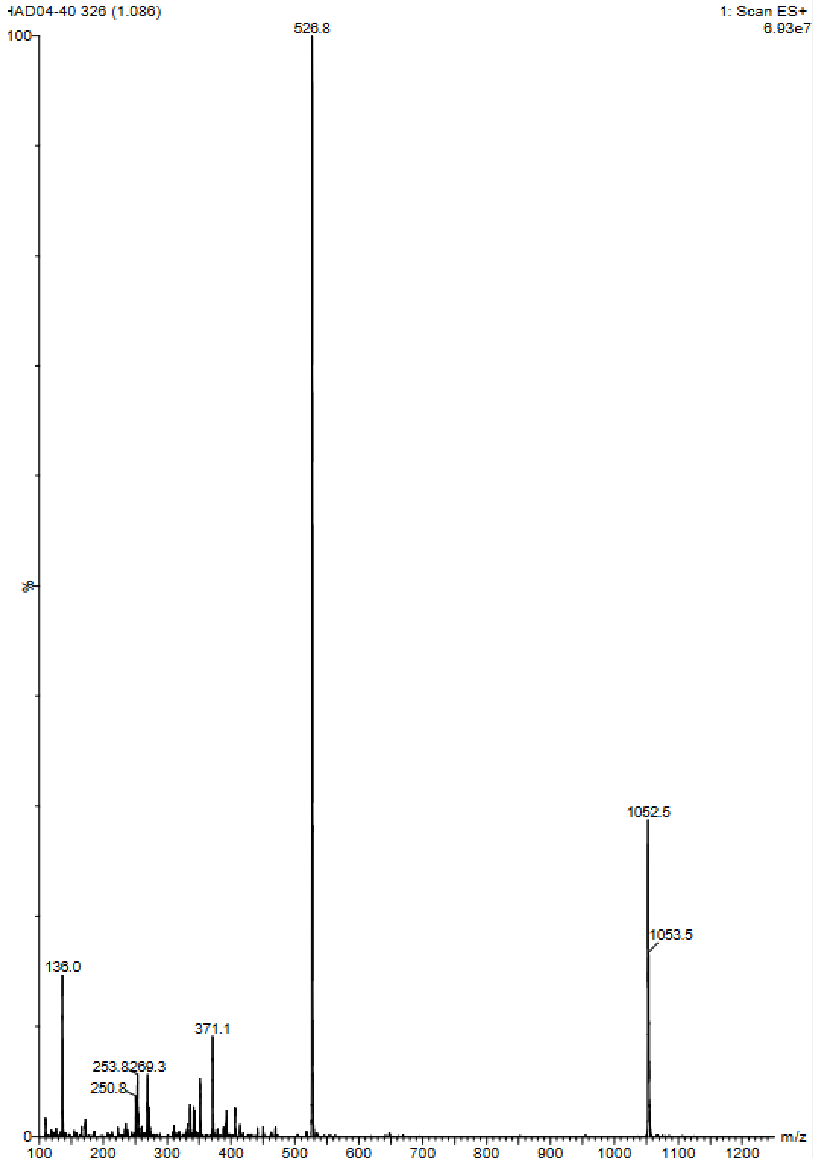

Compound 5: D-R-V-Y-I-H-P-Orn

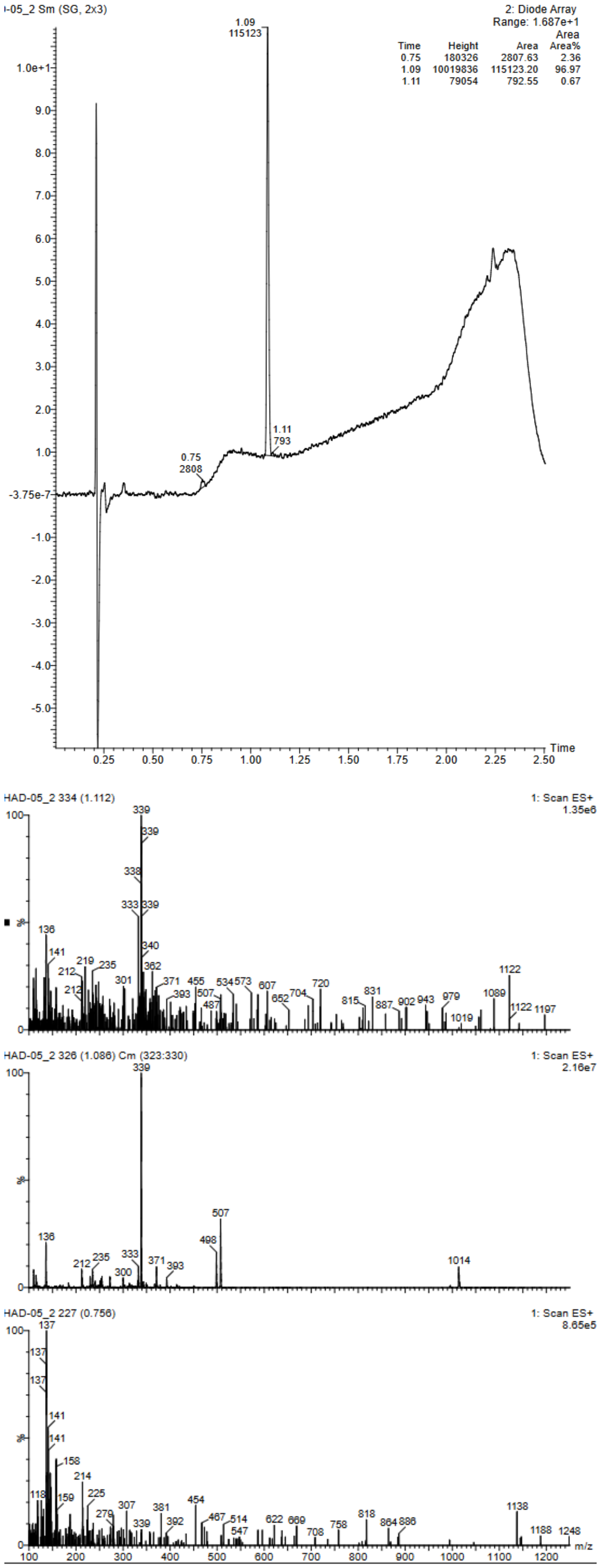

Compound 6:    D-R-V-Y-I-H-P-E

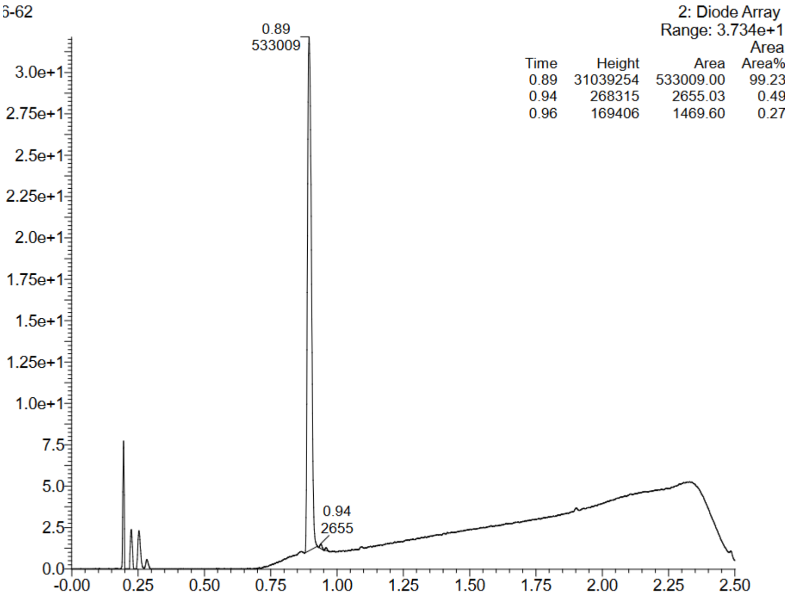


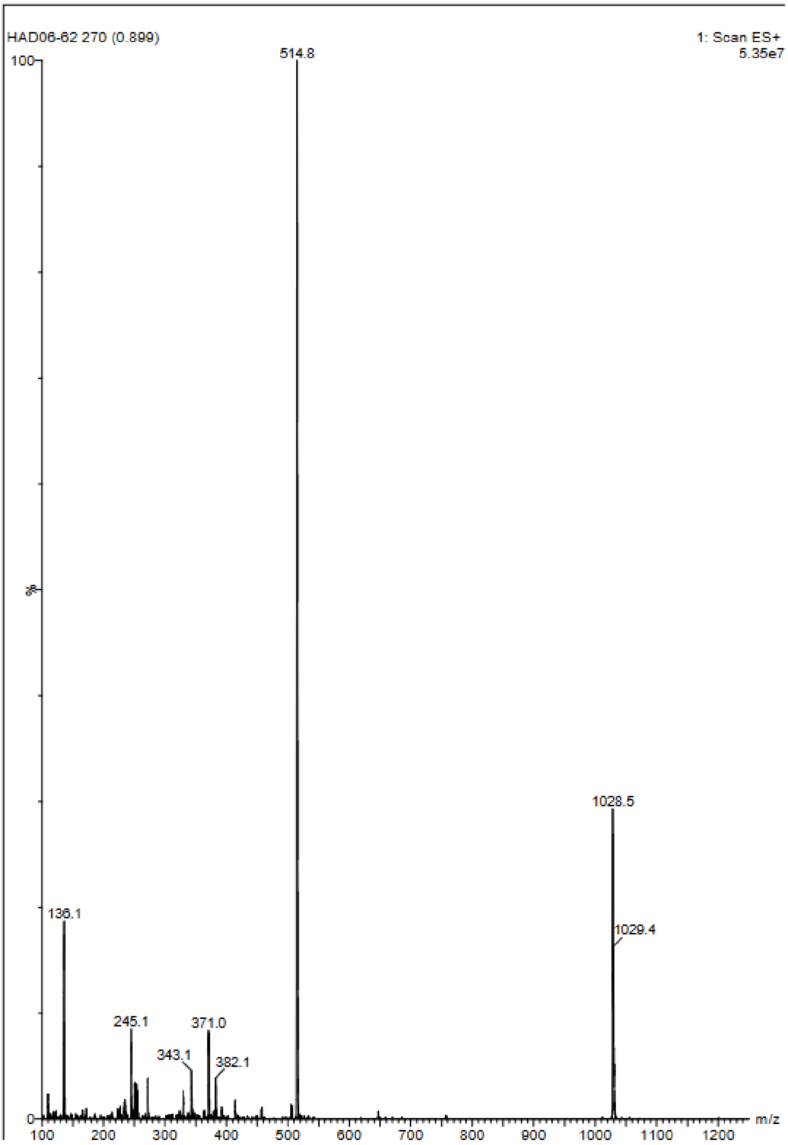

Compound 7:    D-R-V-Y-I-H-P-Q

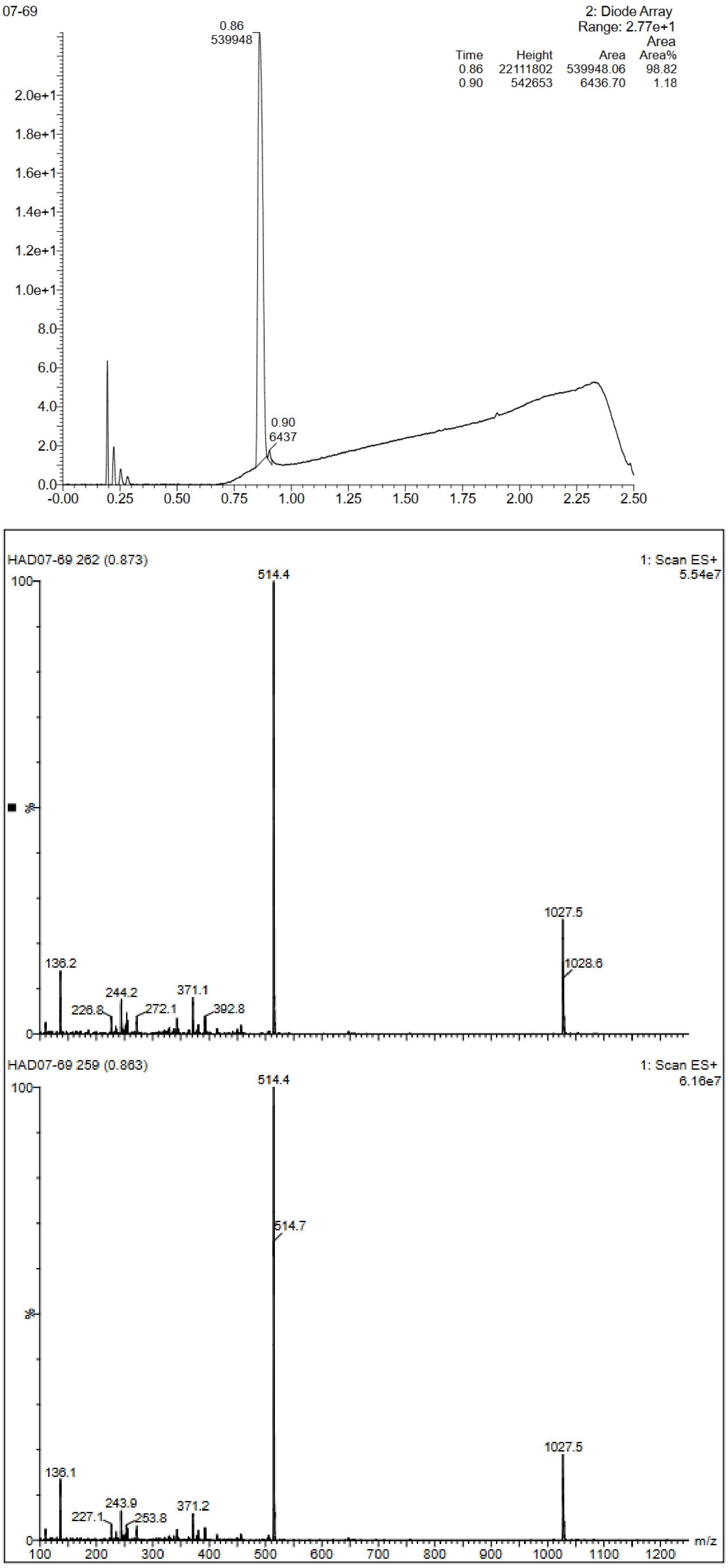

Compound 8:  D-R-V-Y-I-H-P-1Nal

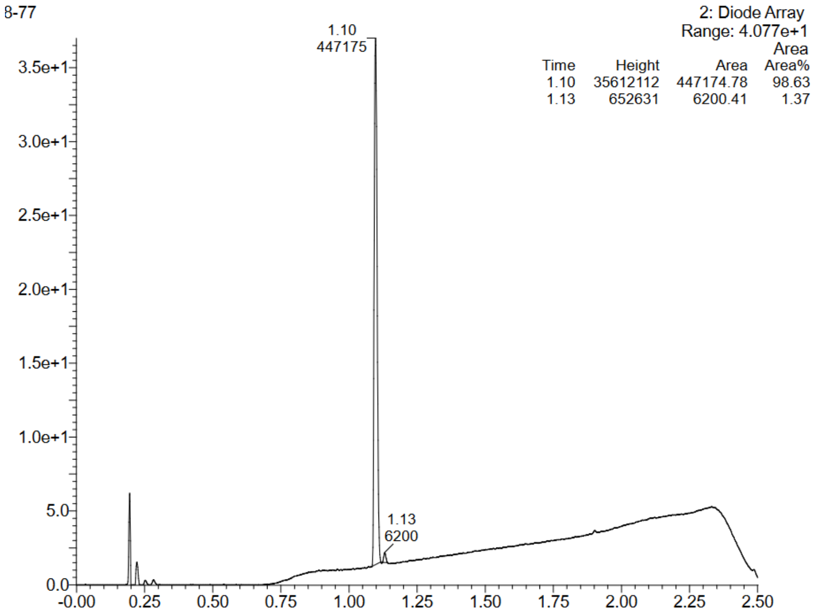


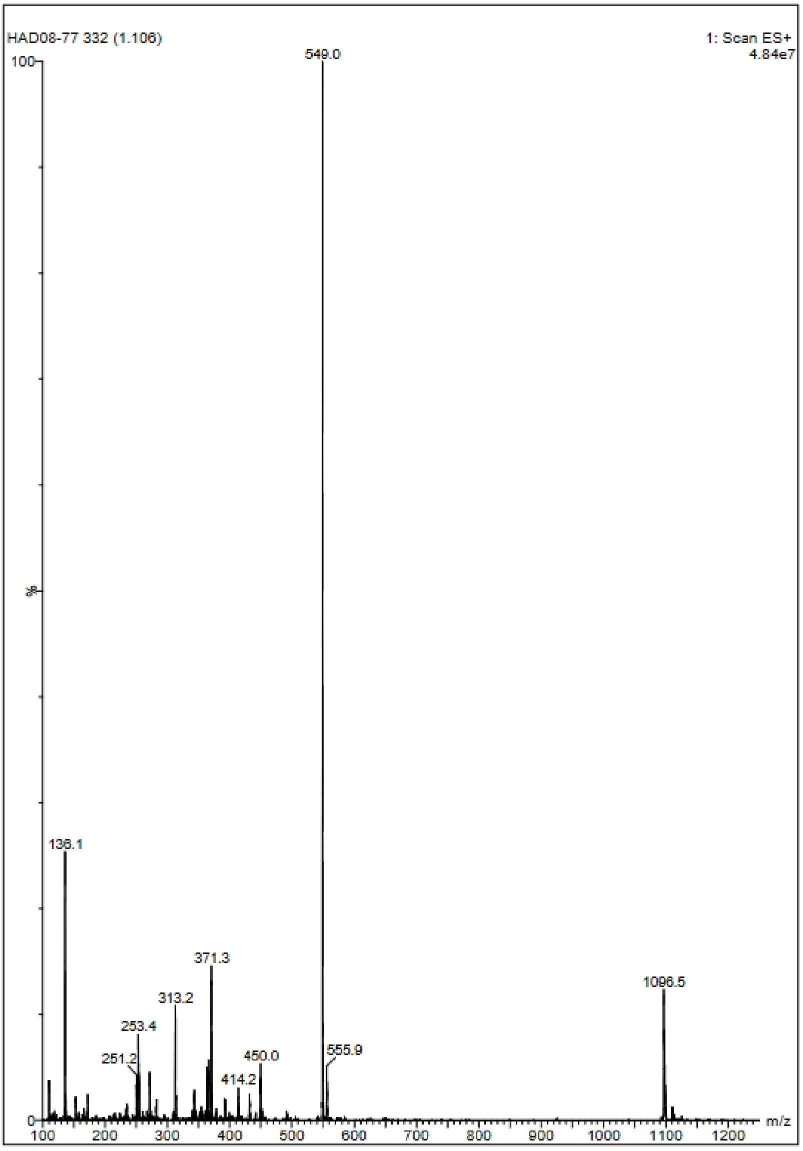

Compound 9: D-R-V-Y-I-H-P-2Nal

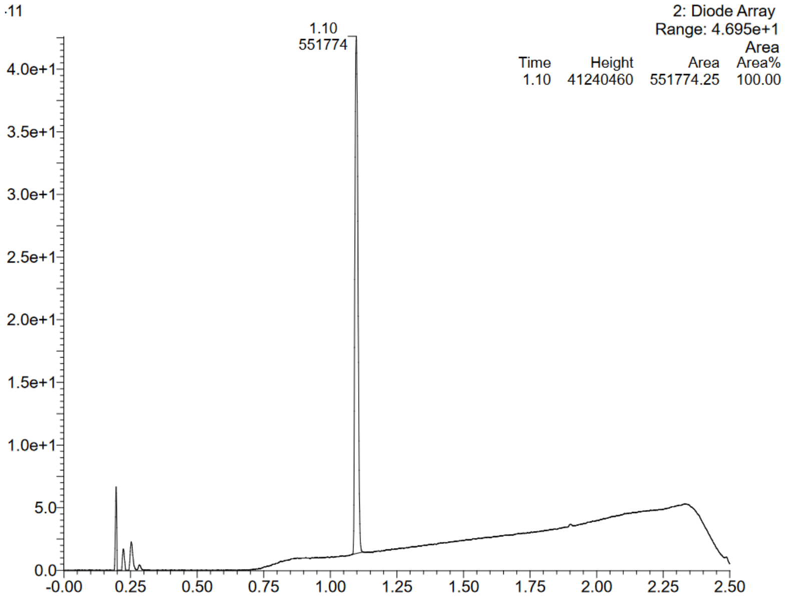


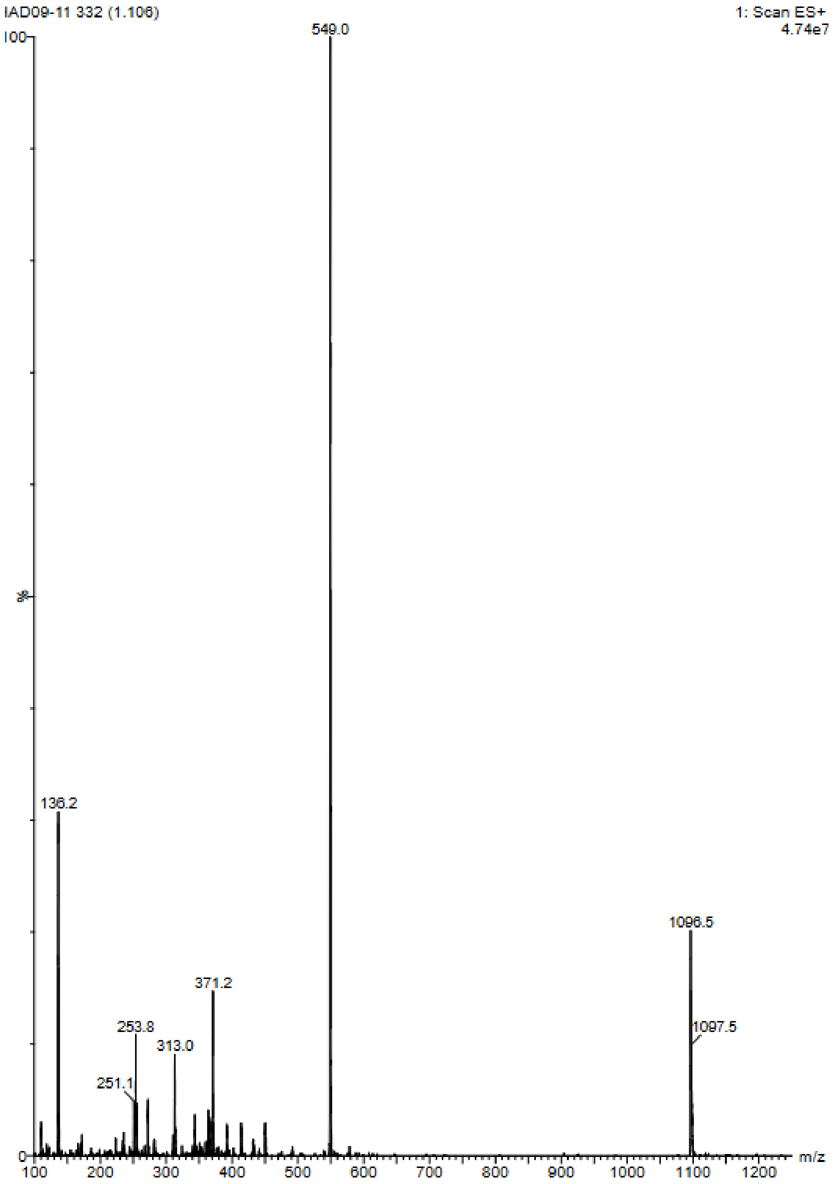

Compound 10:  D-R-V-Y-I-H-P-Tyr(All)

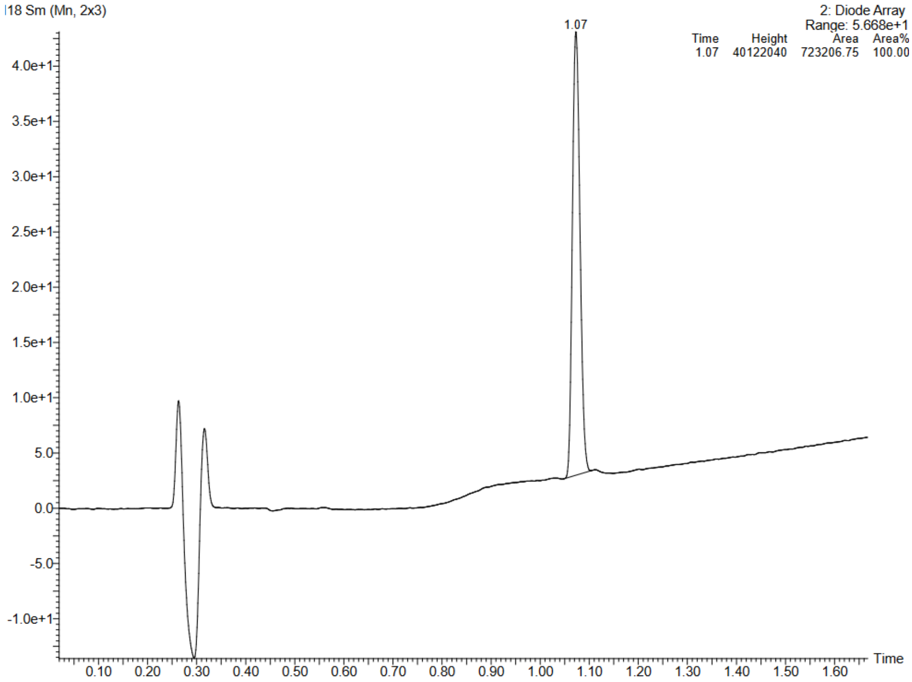


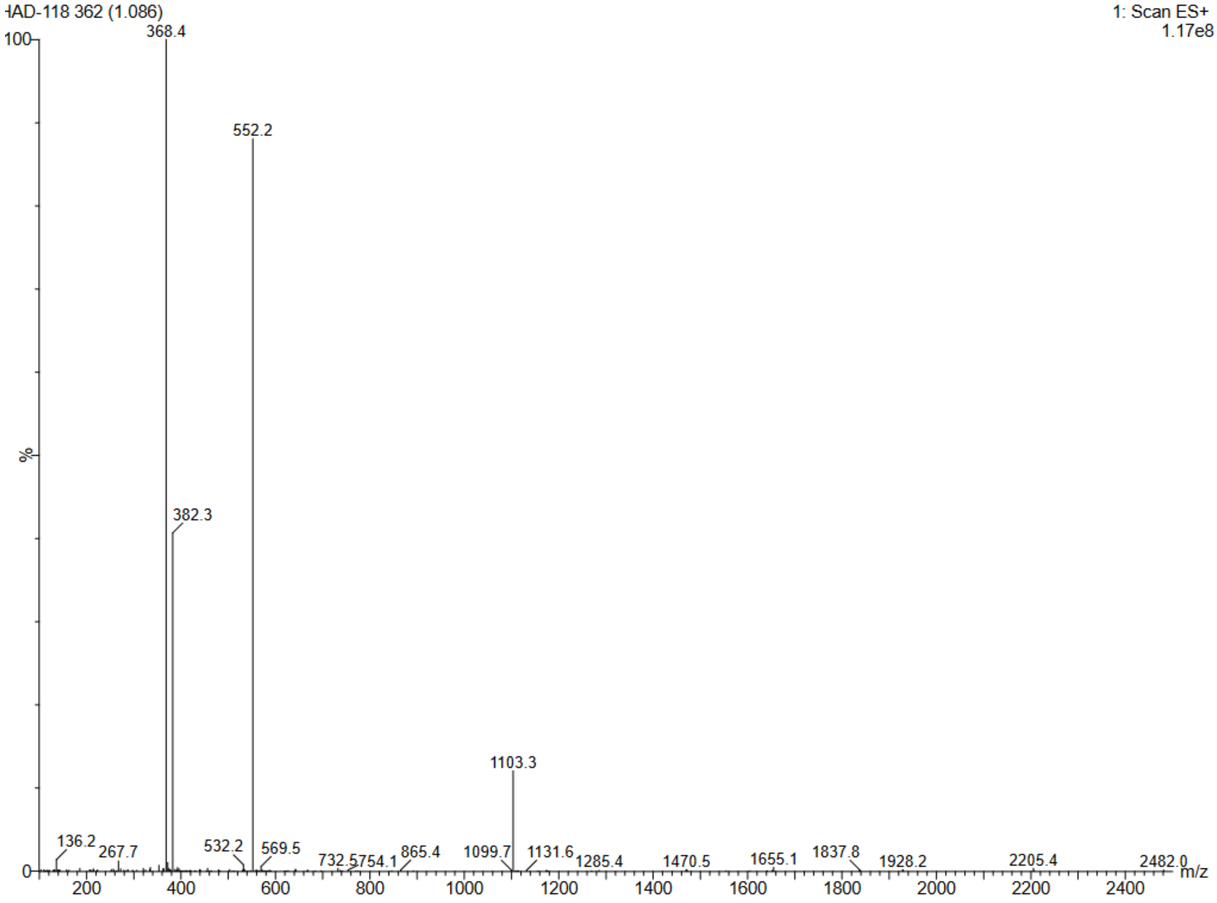

Compound 11: D-R-V-Y-I-H-P-Tyr(Me)

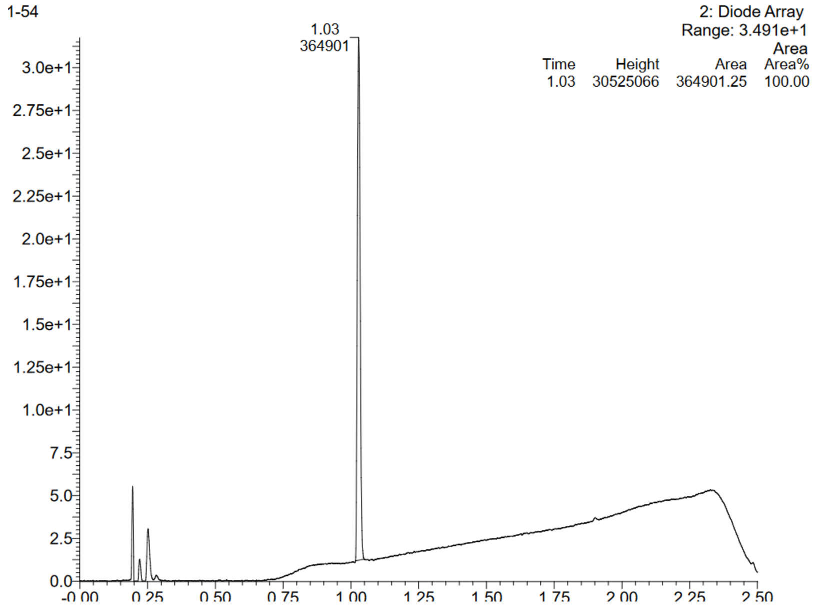


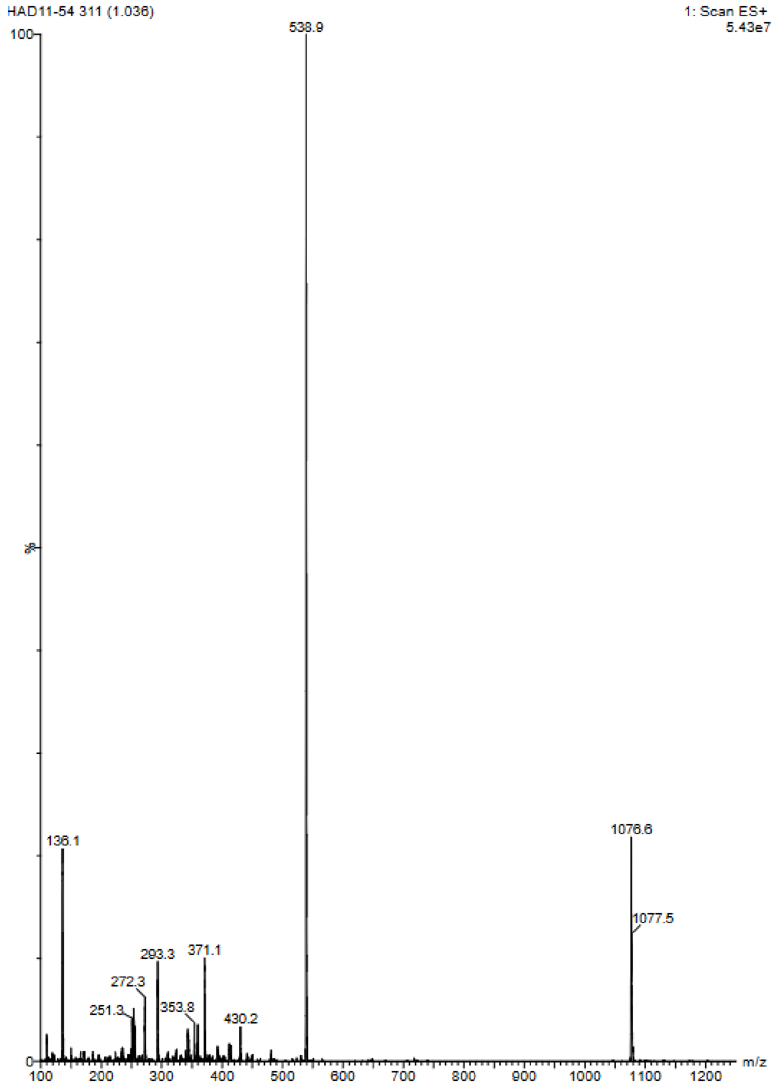

Compound 12: D-R-V-Y-I-H-P-Tyr(Bzl)

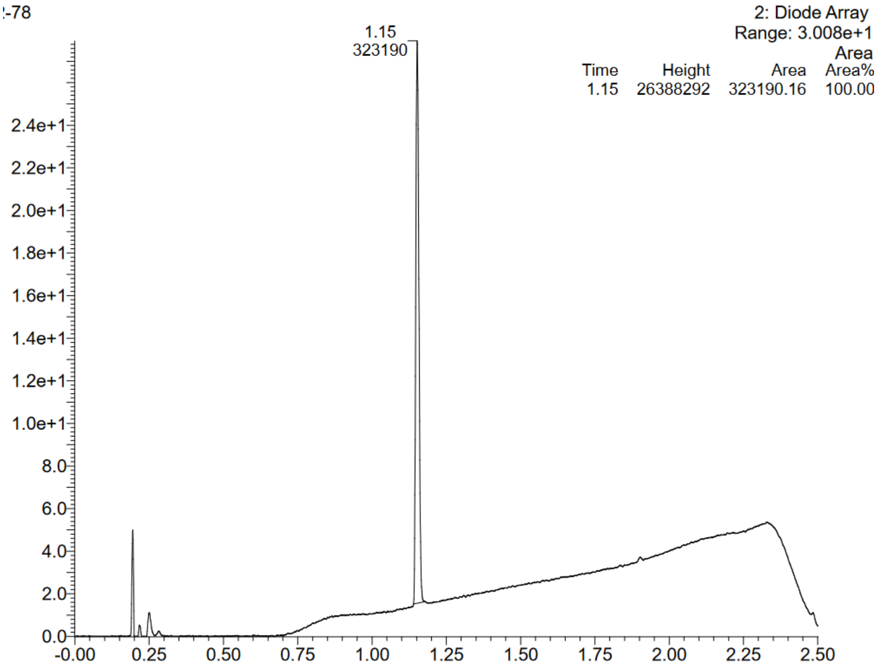


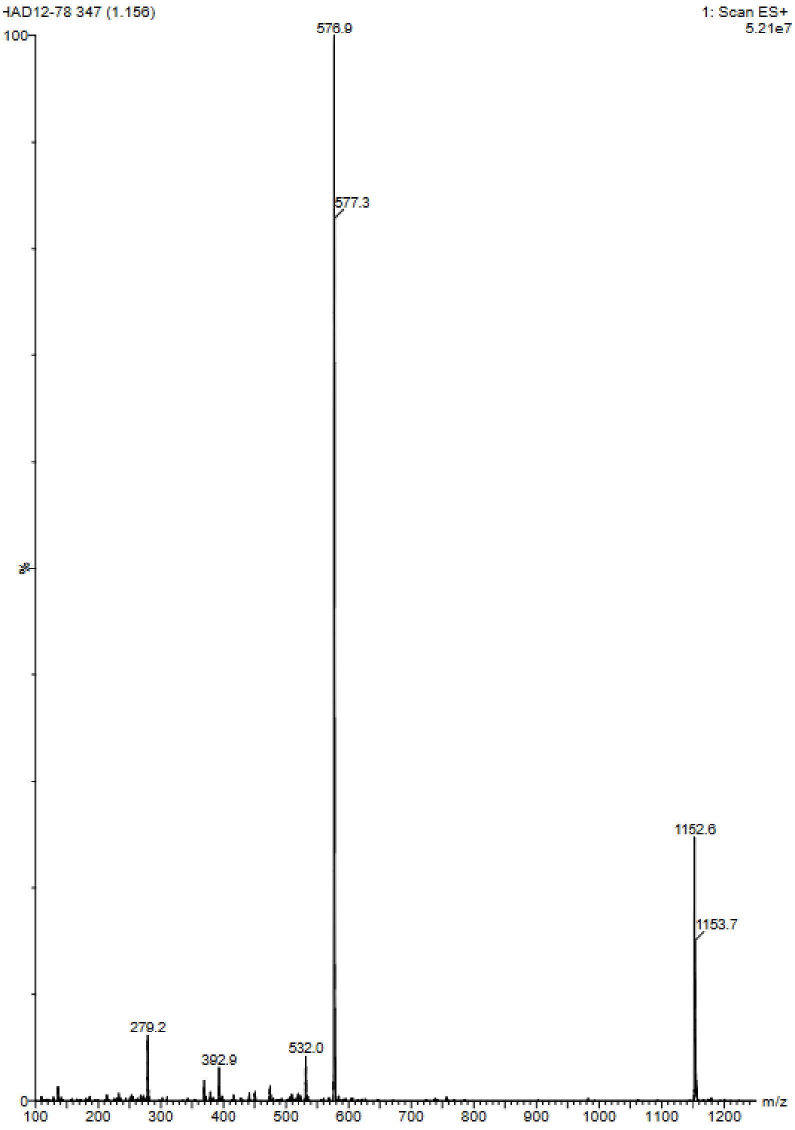

Compound 13: D-R-V-Y-I-H-P-Bip

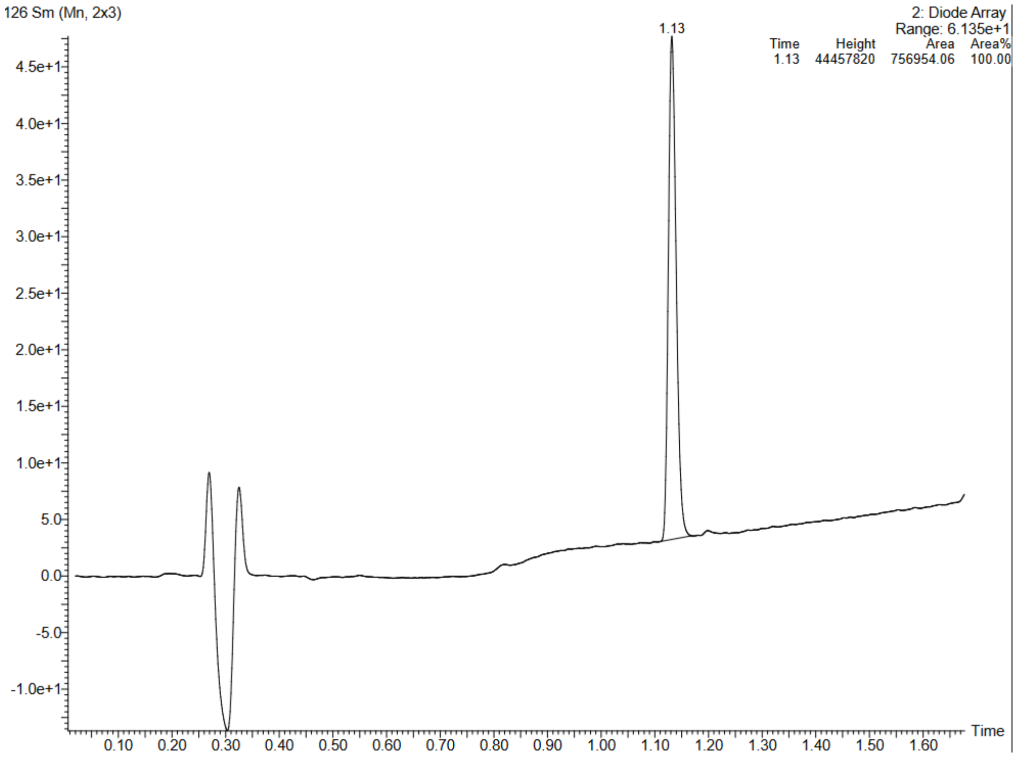


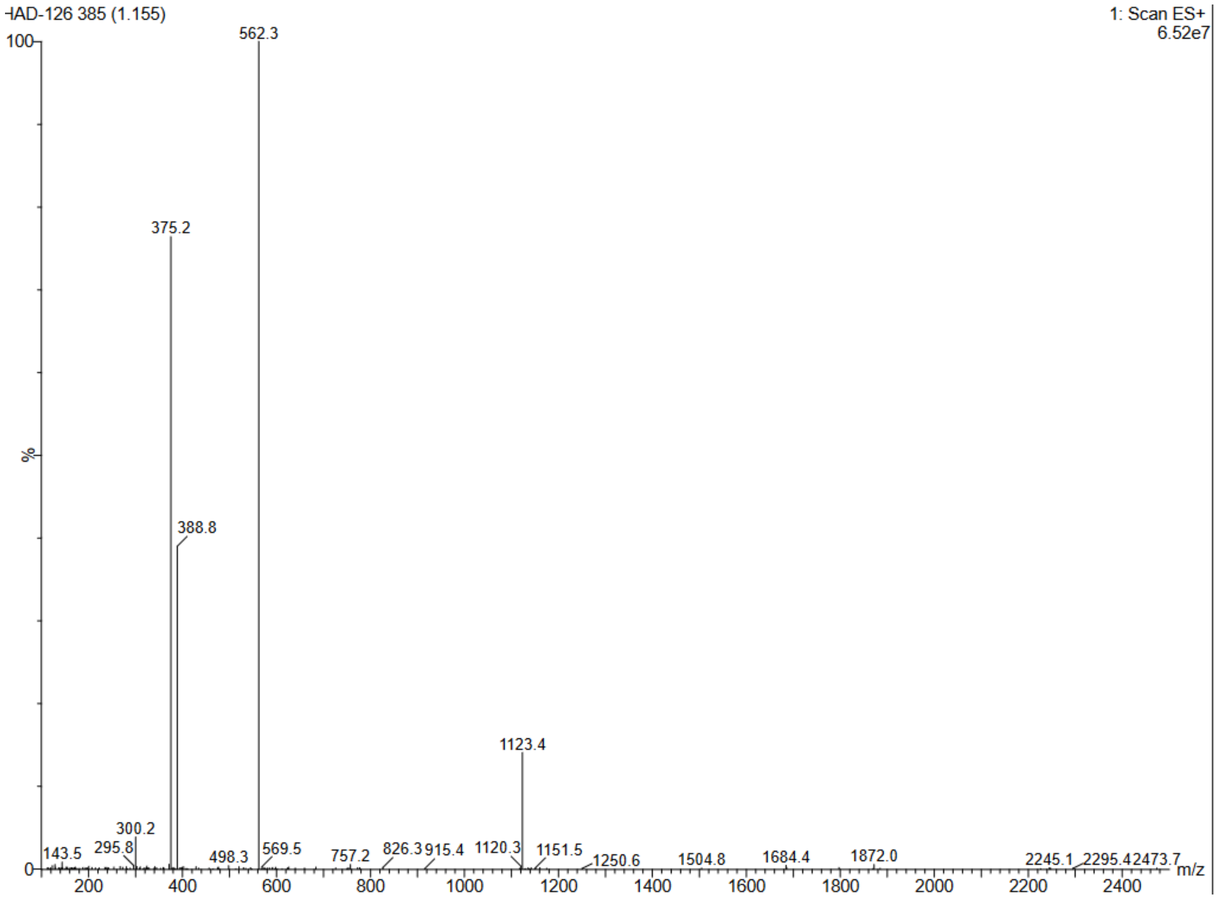

Compound 14:  D-R-V-Y-I-H-P-p-tBuPhe

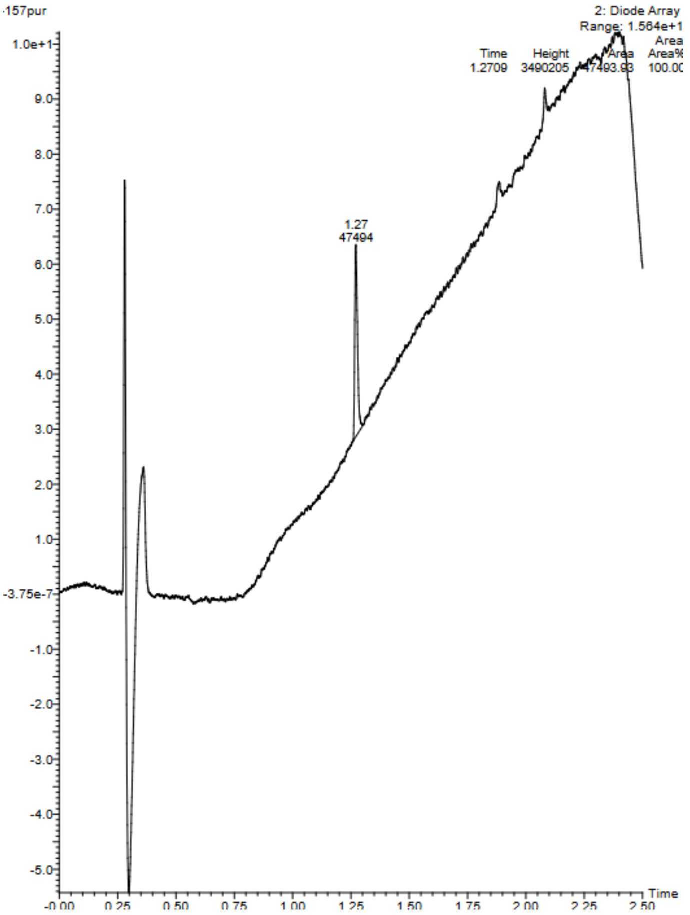


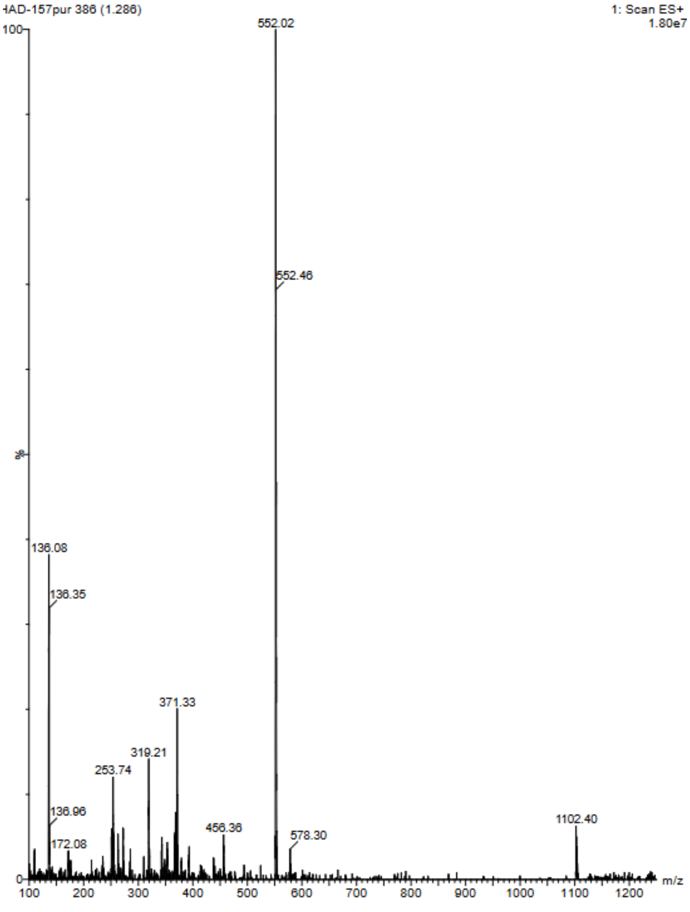

Compound 15: D-R-V-Y-I-H-P-Dip

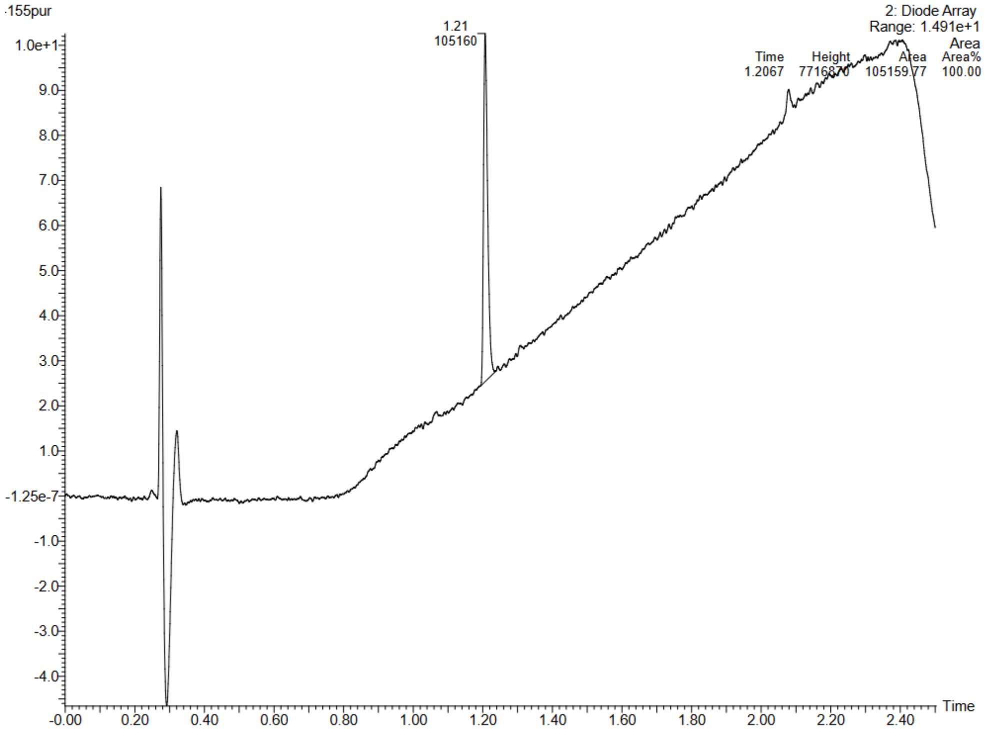


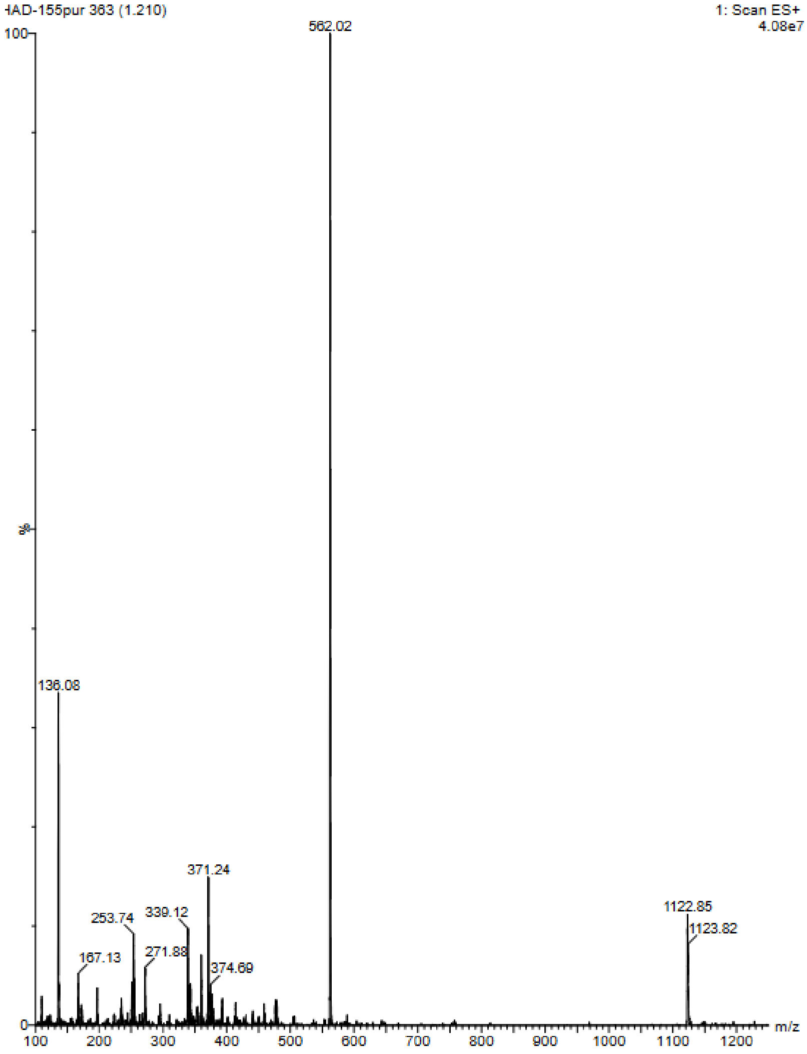

Compound 16: D-R-V-Y-I-H-P-Ser(Bzl)

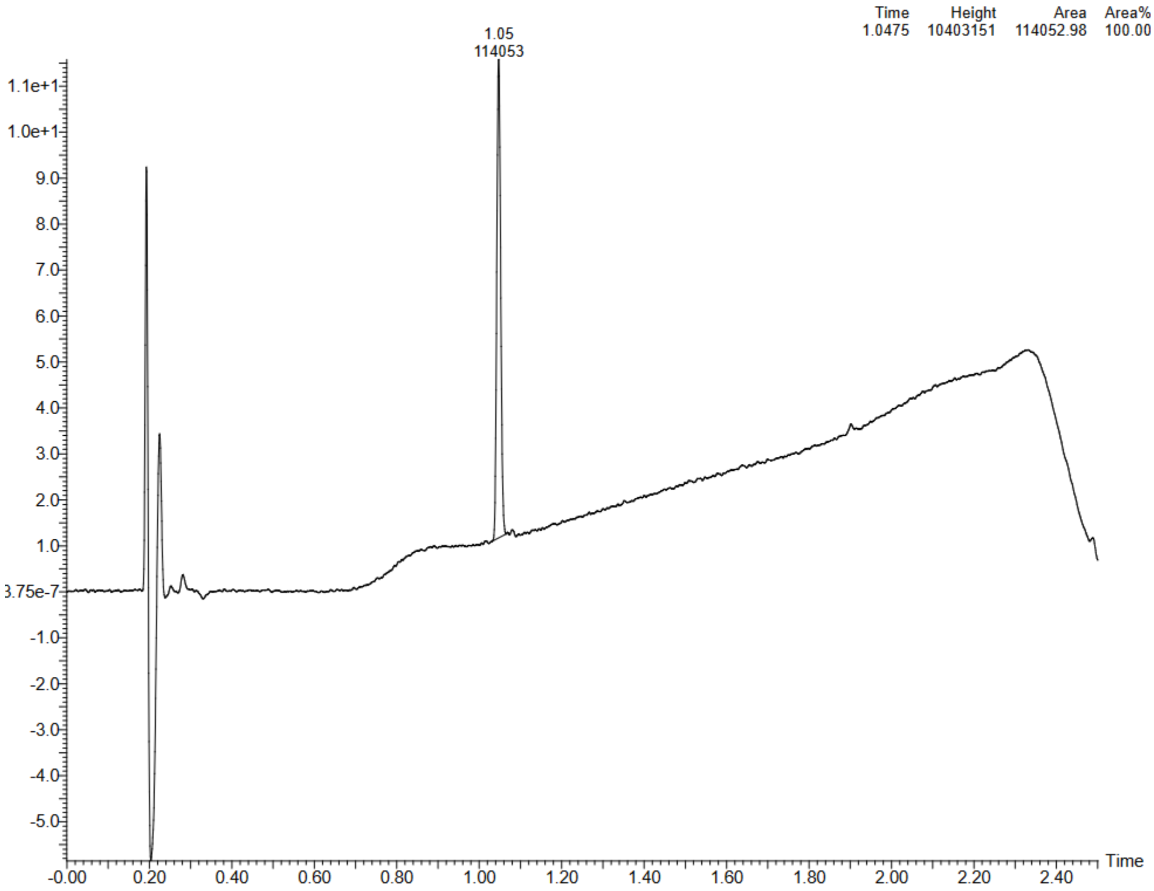


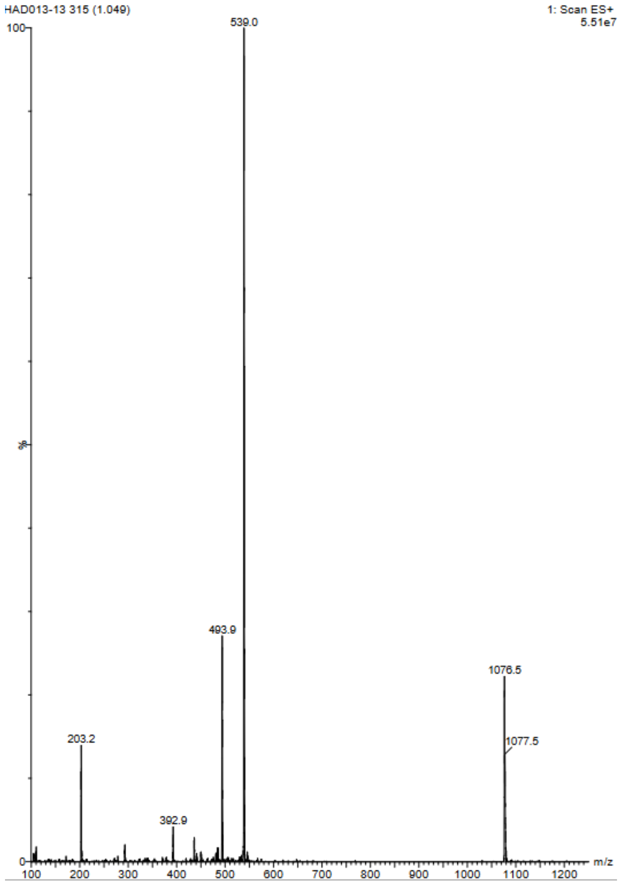

Compound 17: D-R-V-Y-I-H-P-4Pyridyl

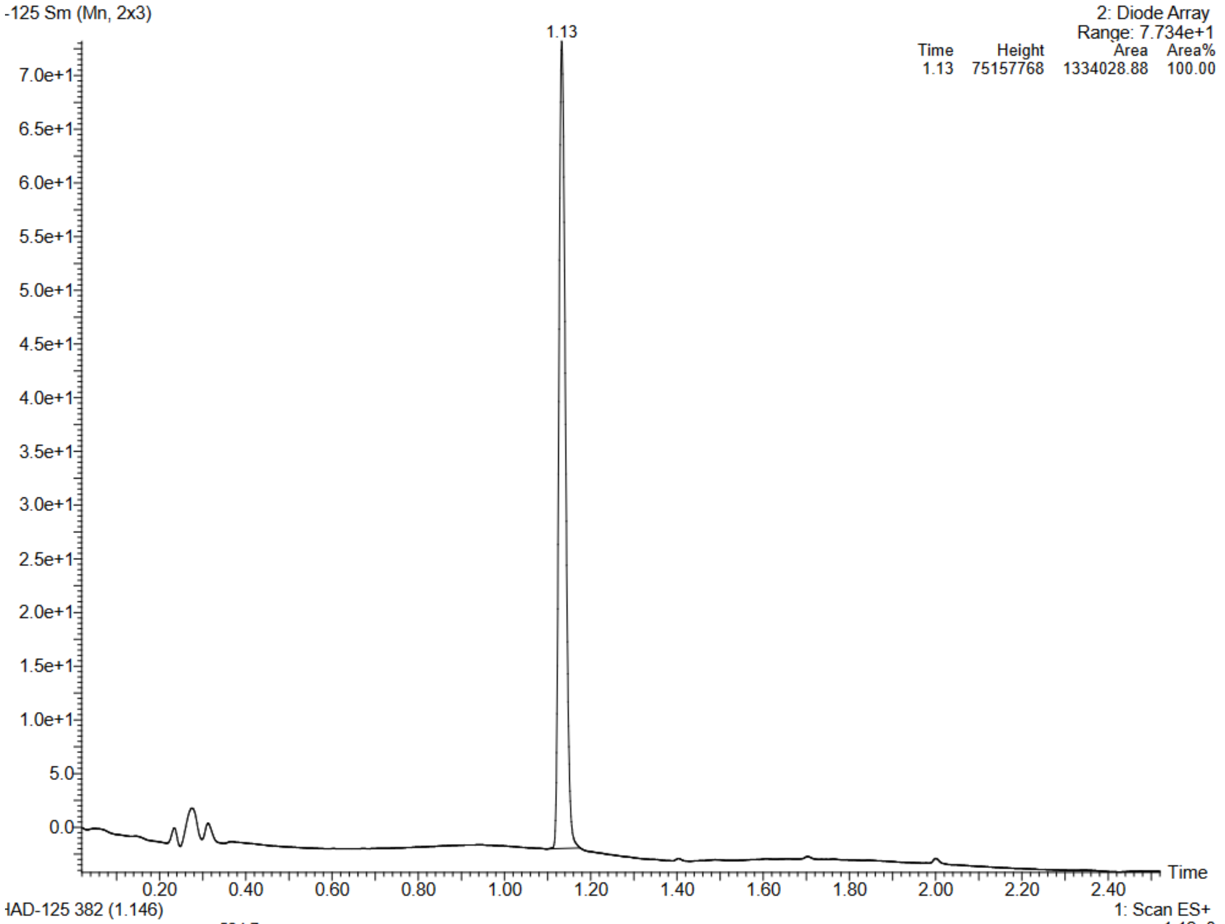


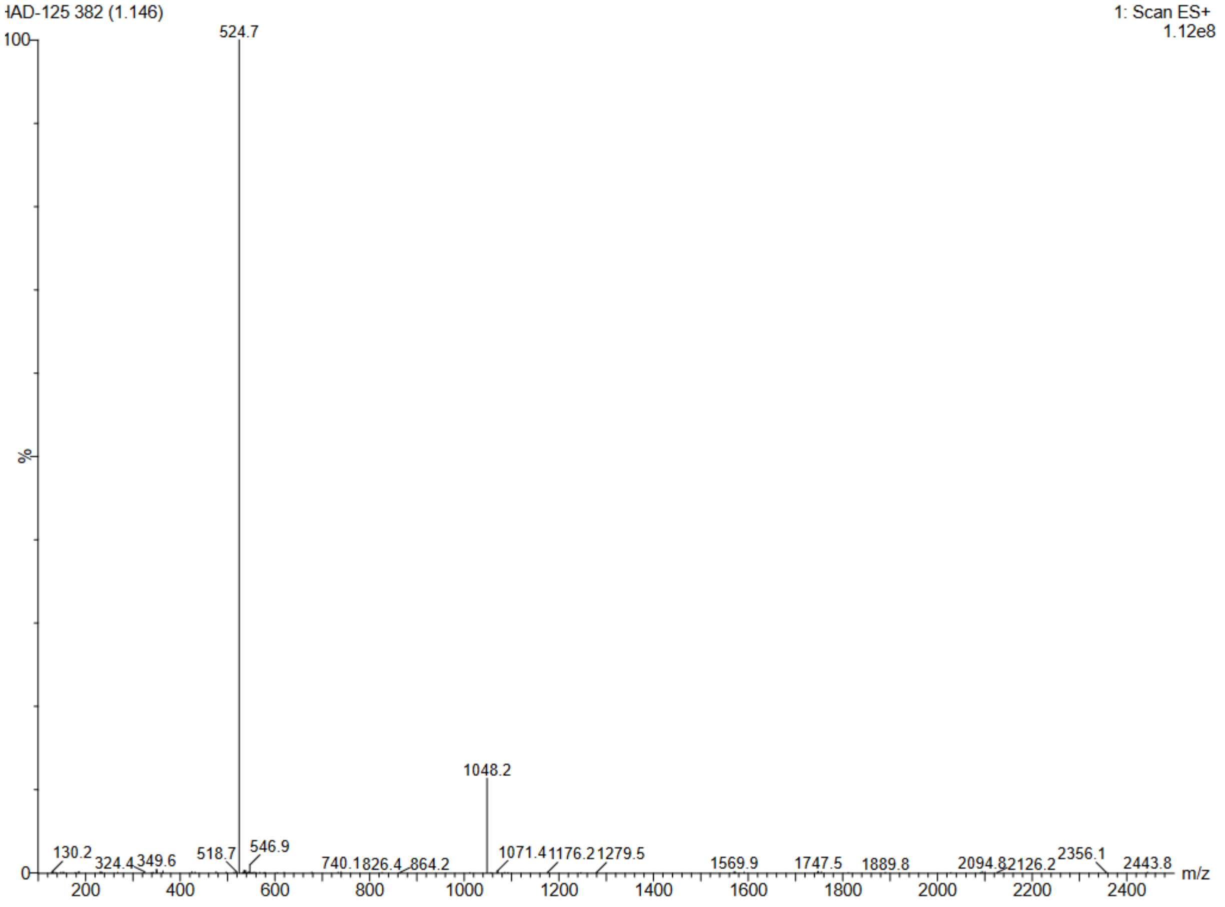

Compound 18 : D-R-V-Y-I-H-P-3Pyridyl

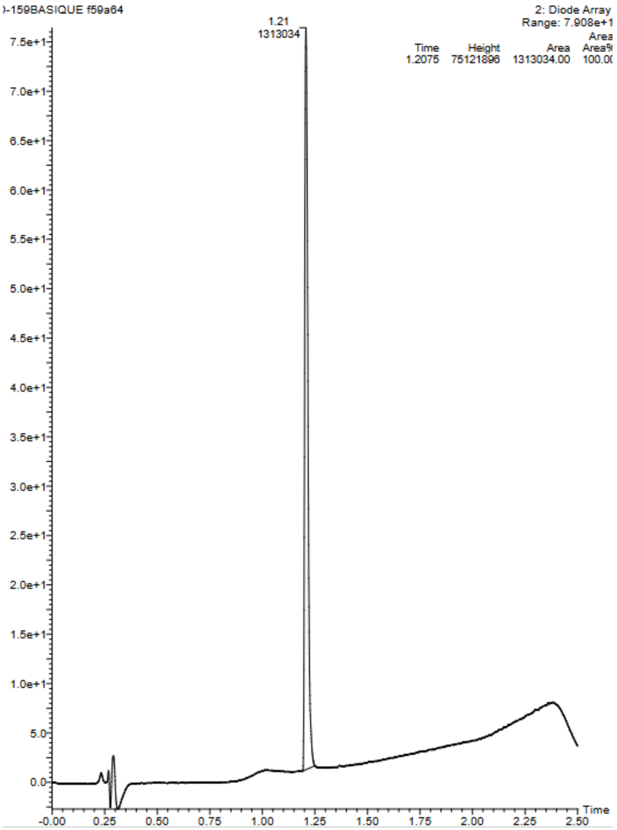


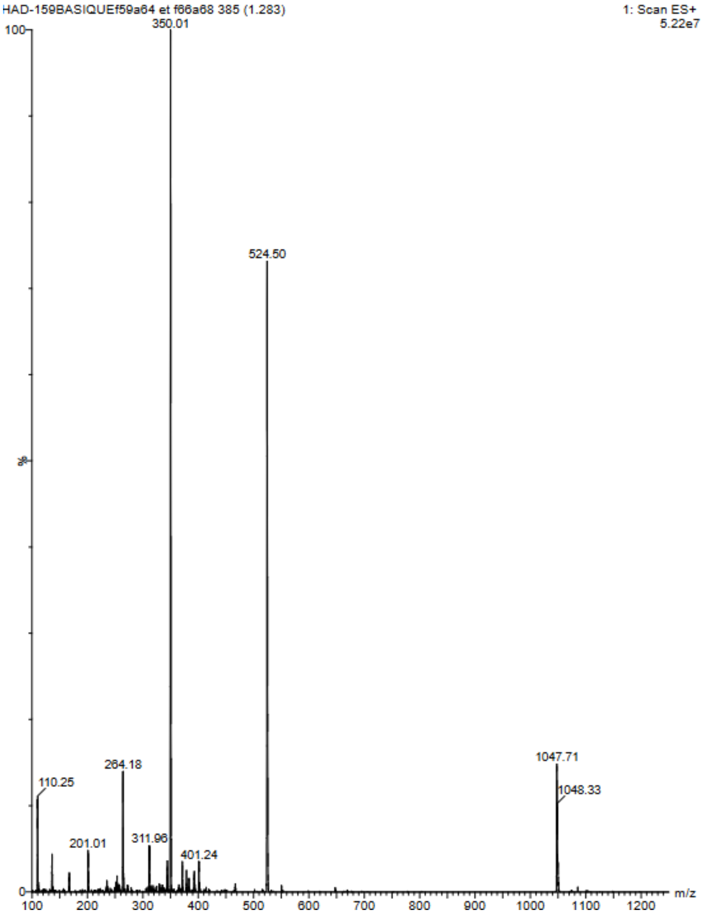

Compound 19: D-R-V-Y-I-H-P-2Pyridyl

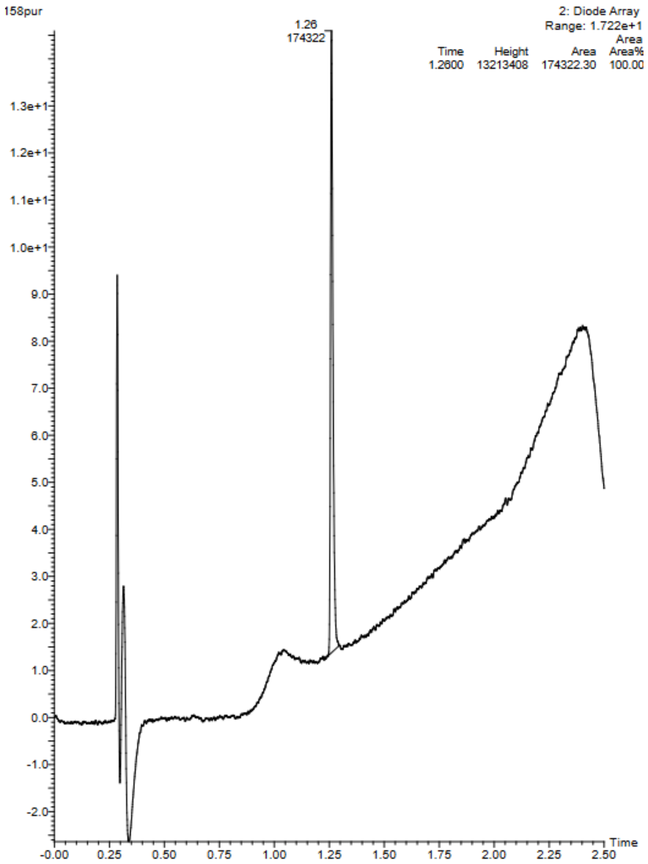


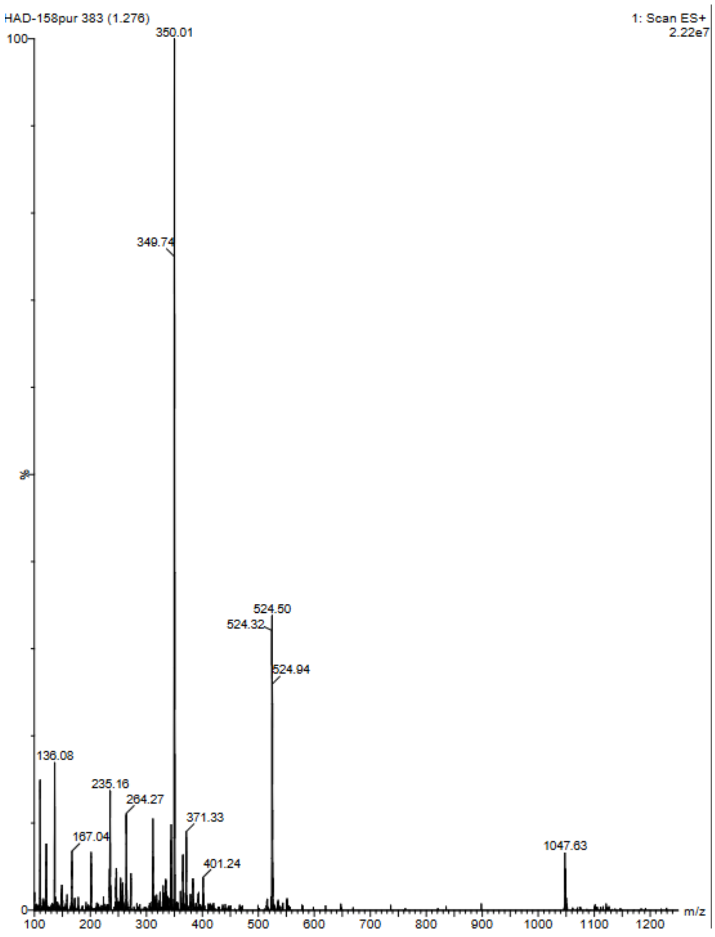

Compound 20 : D-R-V-Y-I-H-P-4ThzAla

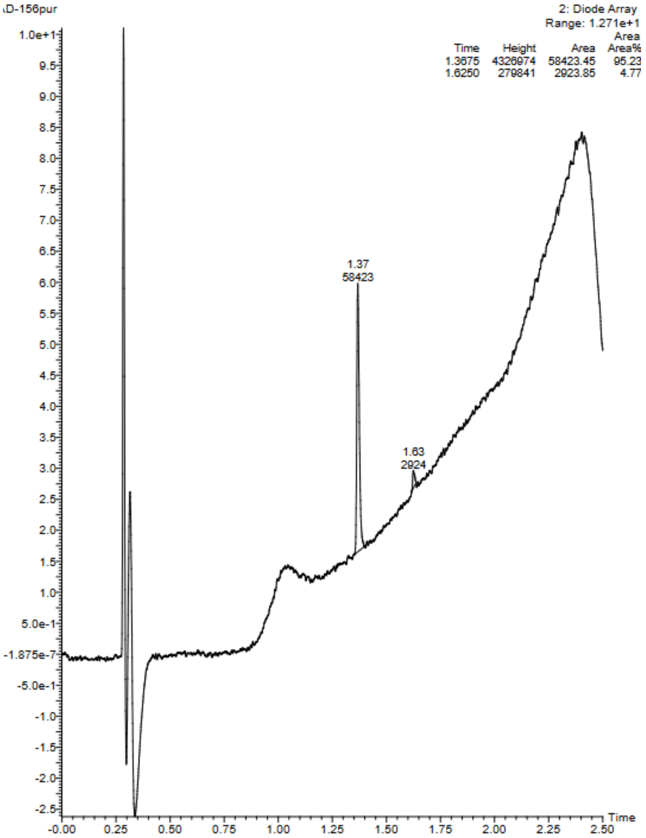


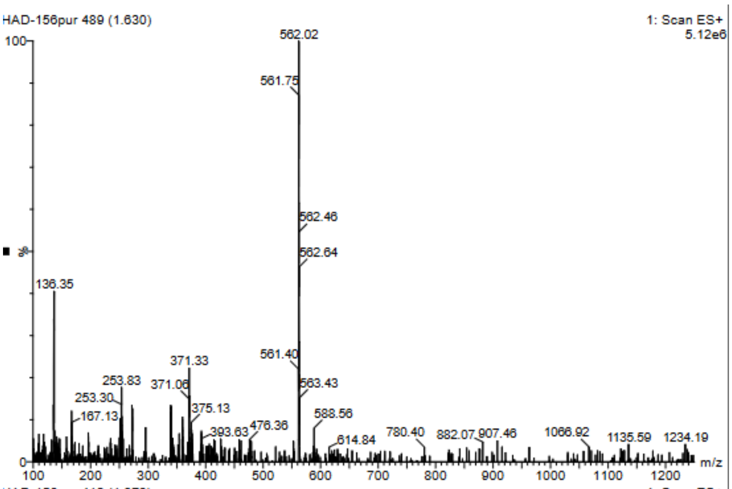


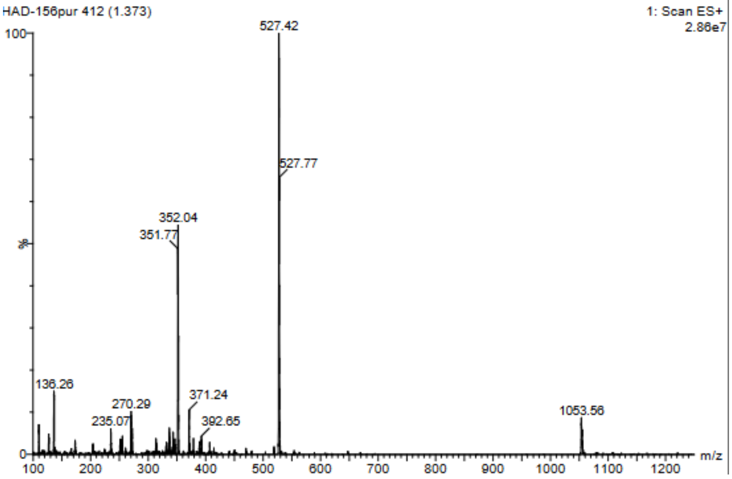

Compound 21: D-R-V-Y-I-H-P-W

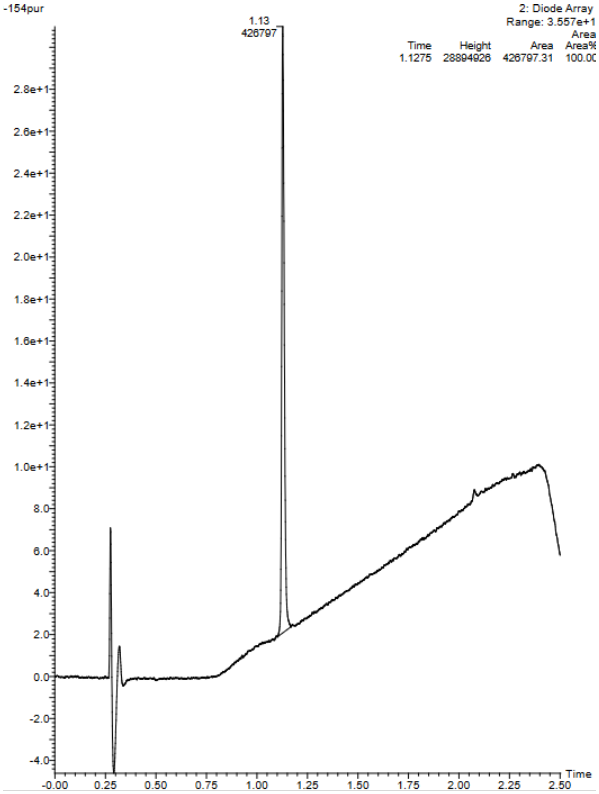


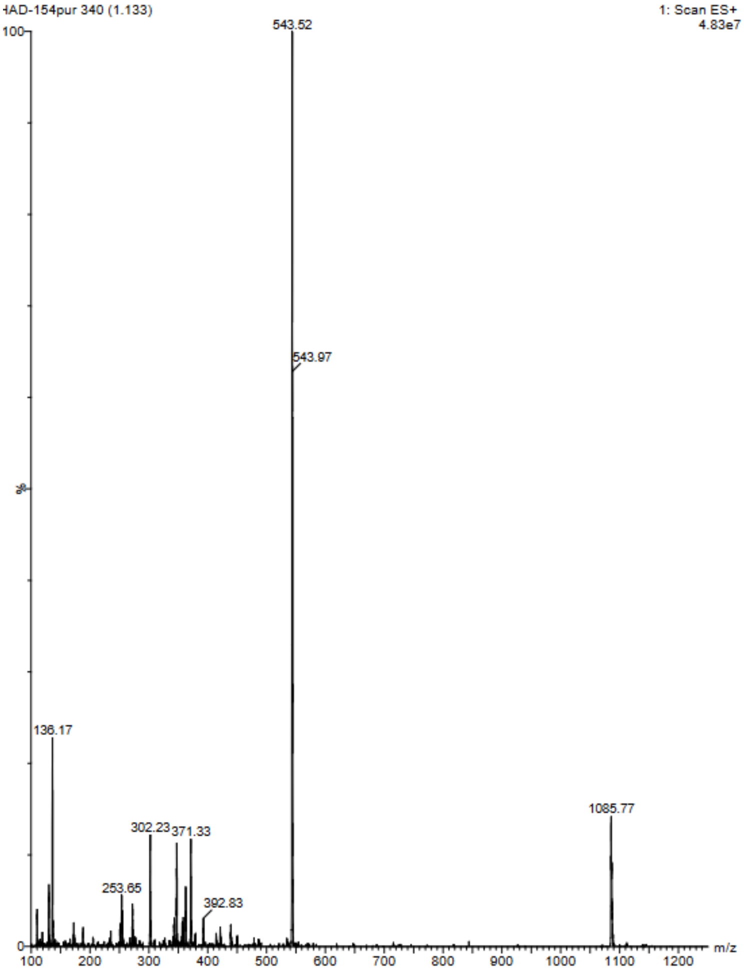

Compound 22a : D-R-V-Y-I-H-P-F(2NO2)

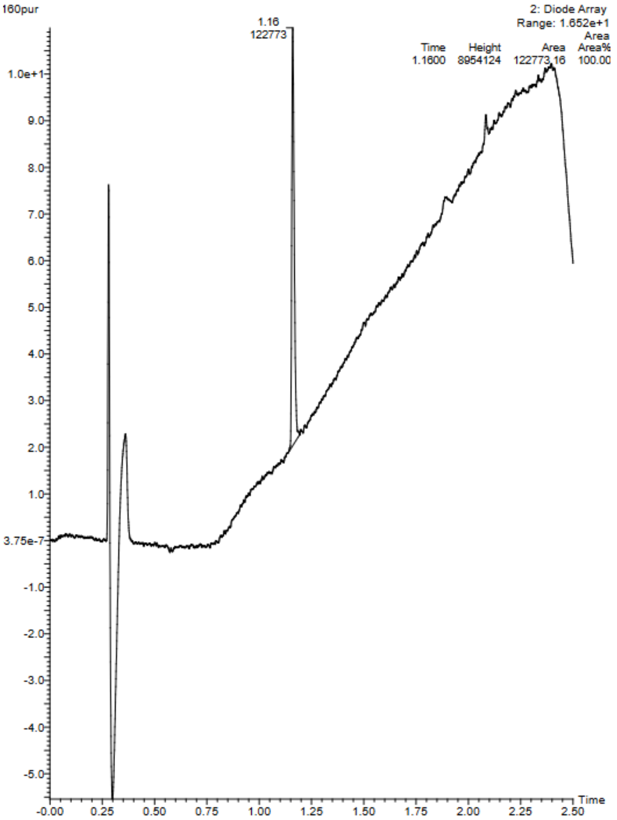


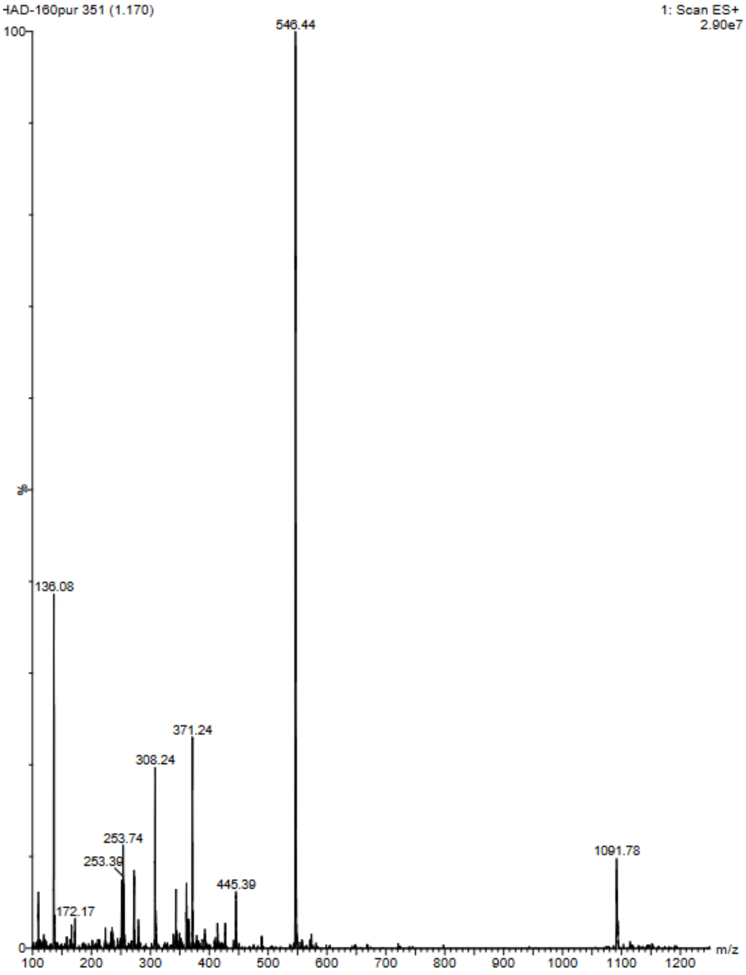

Compound 23a:D-R-V-Y-I-H-P-F(2CN)

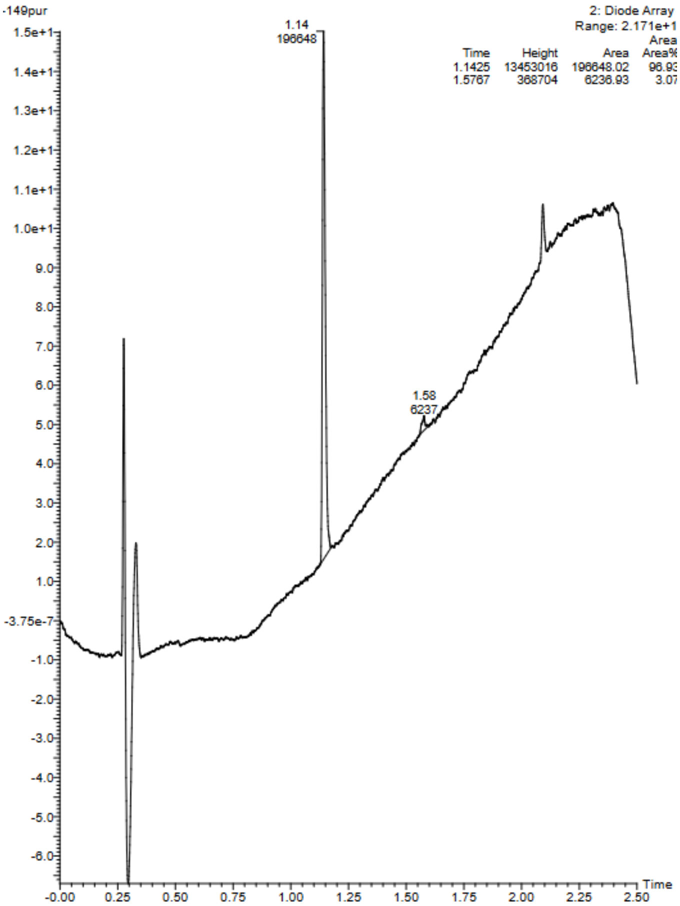


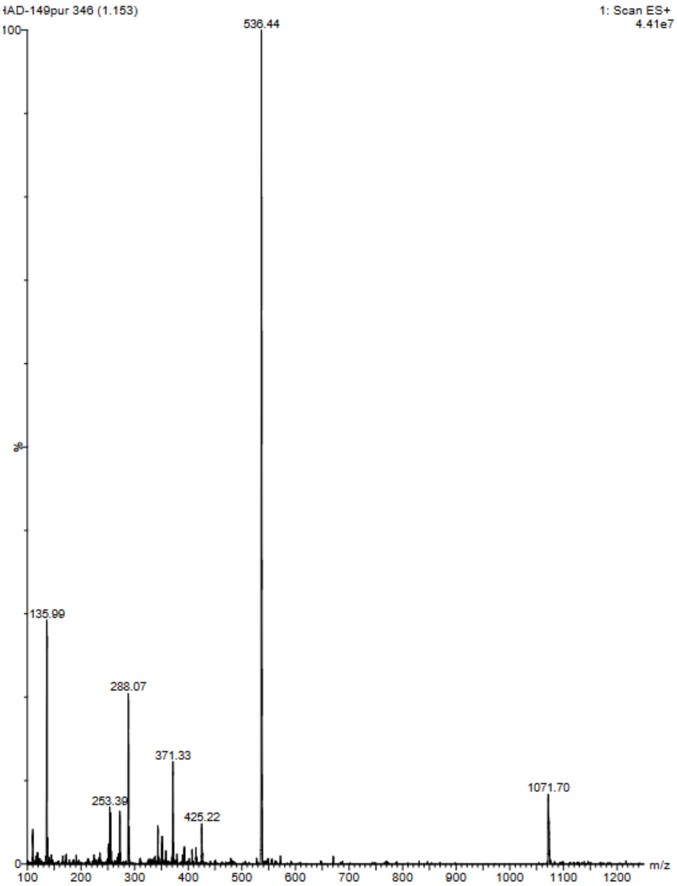

Compound 24a:D-R-V-Y-I-H-P-F(2OH)

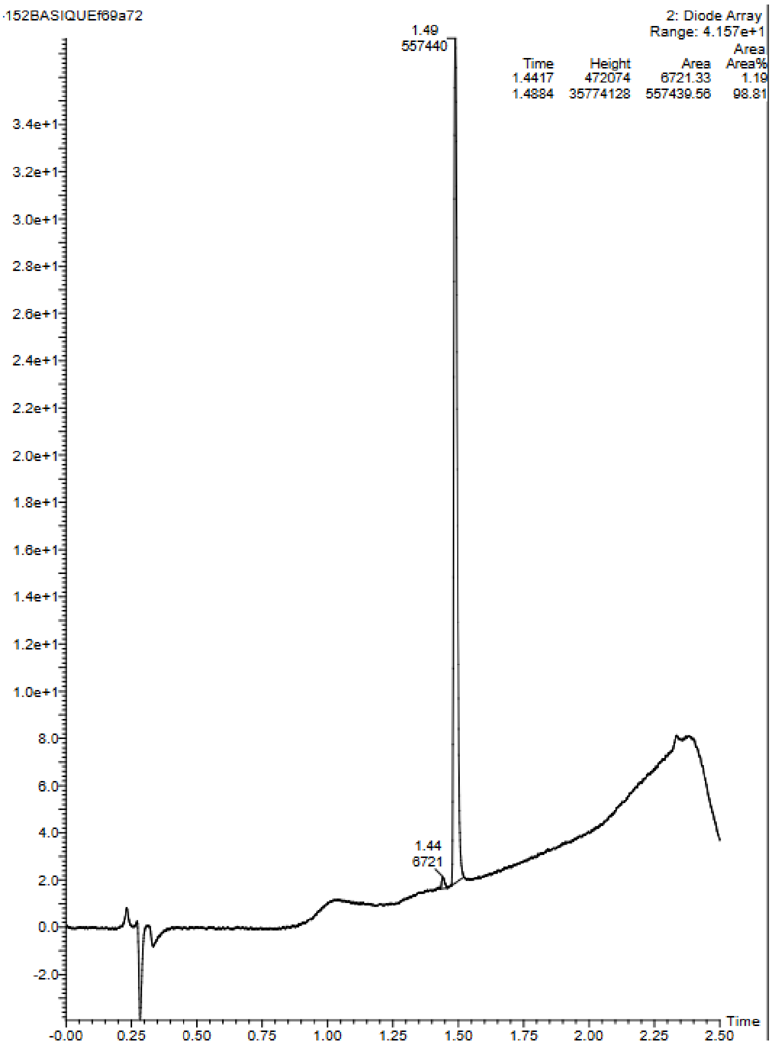


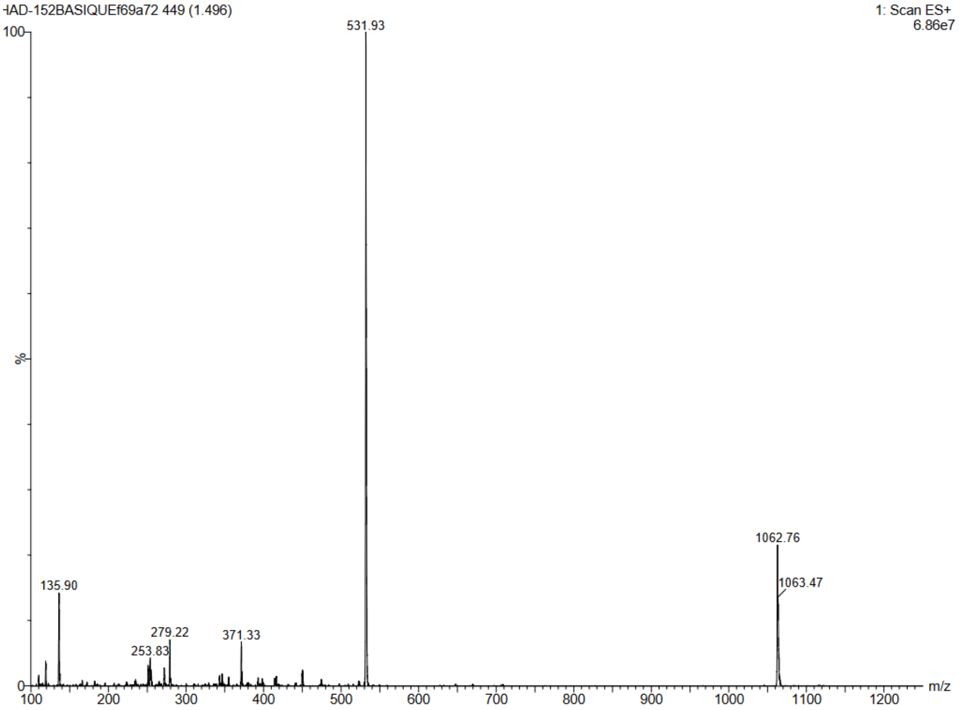

Compound 25a:D-R-V-Y-I-H-P-F(2Me)

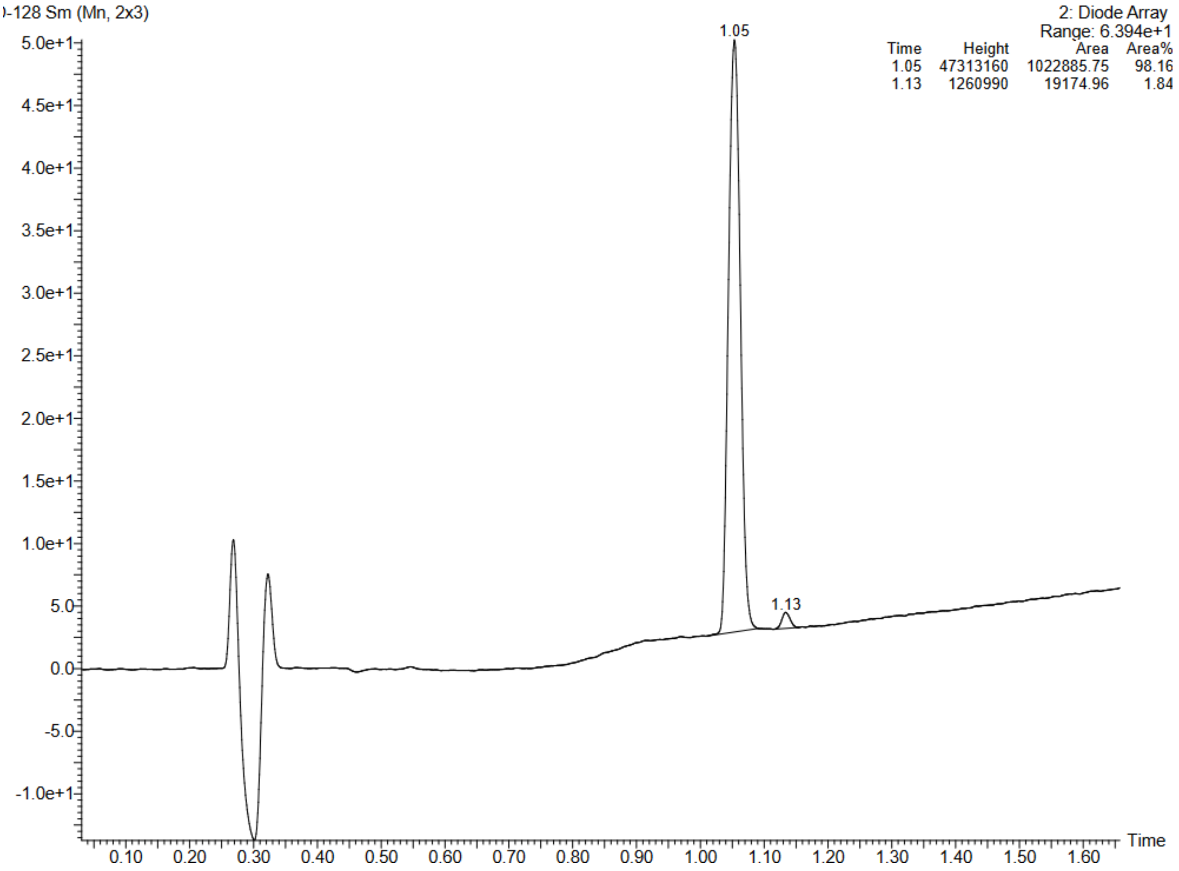


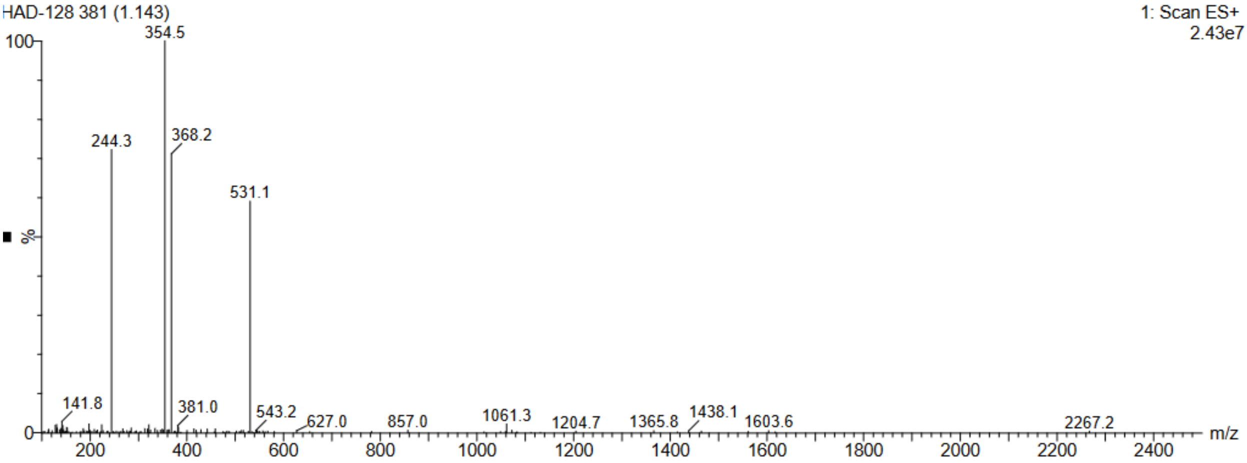


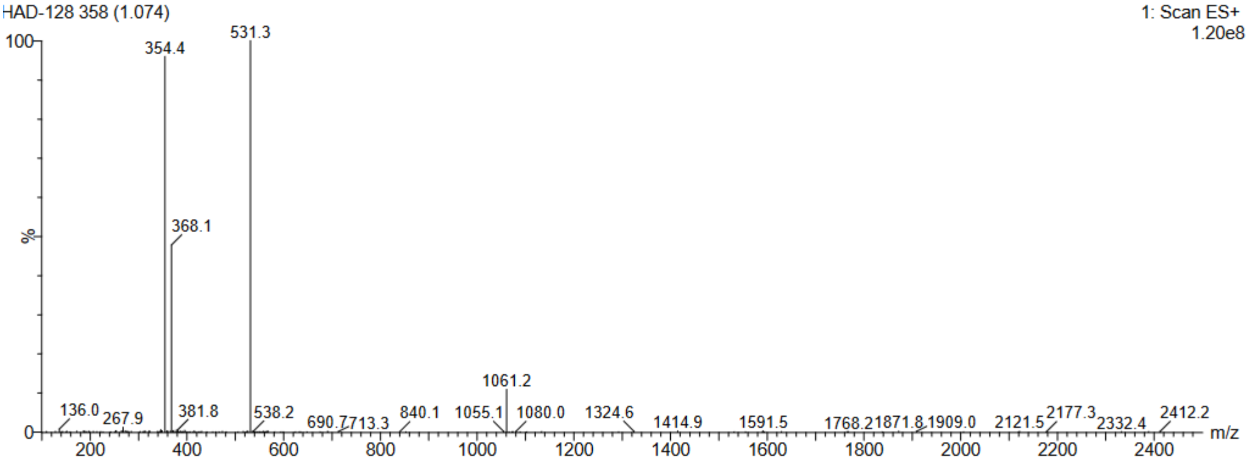

Compound 26a:D-R-V-Y-I-H-P-F(2CF3)

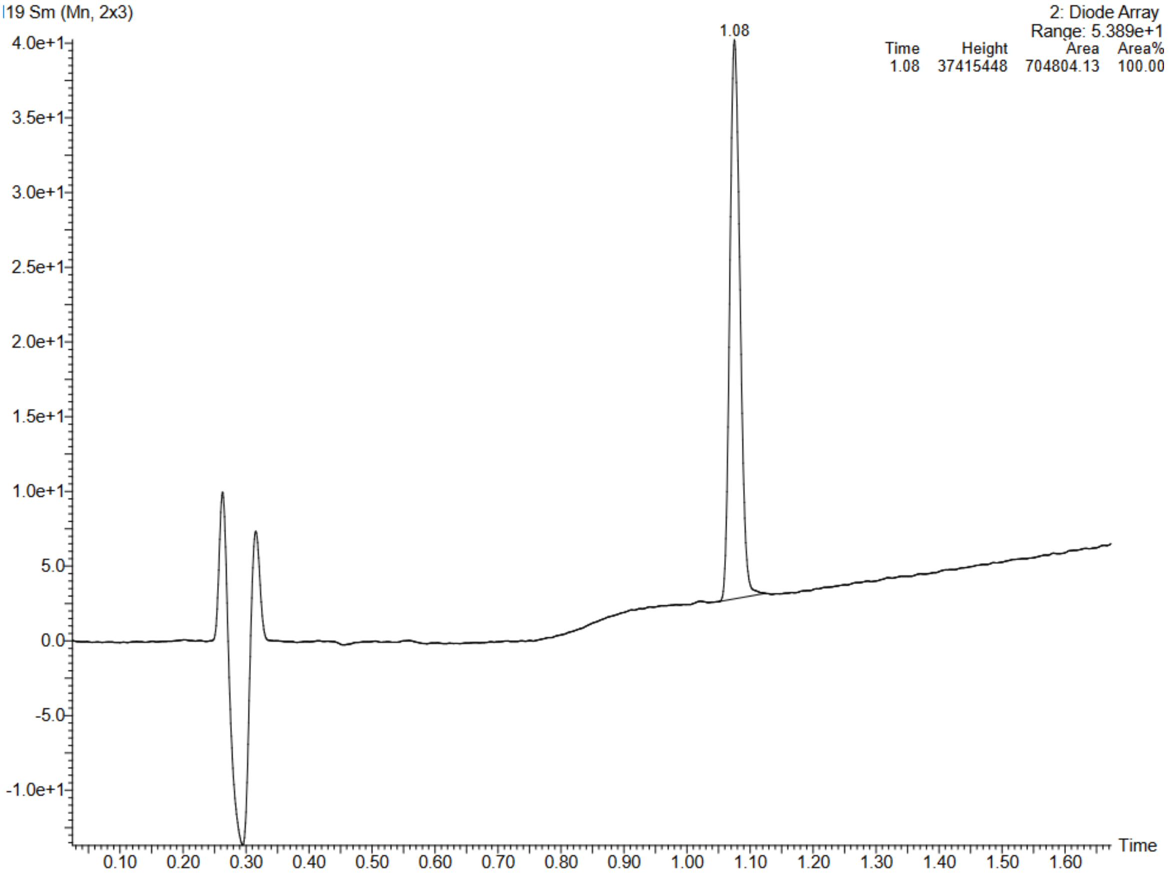


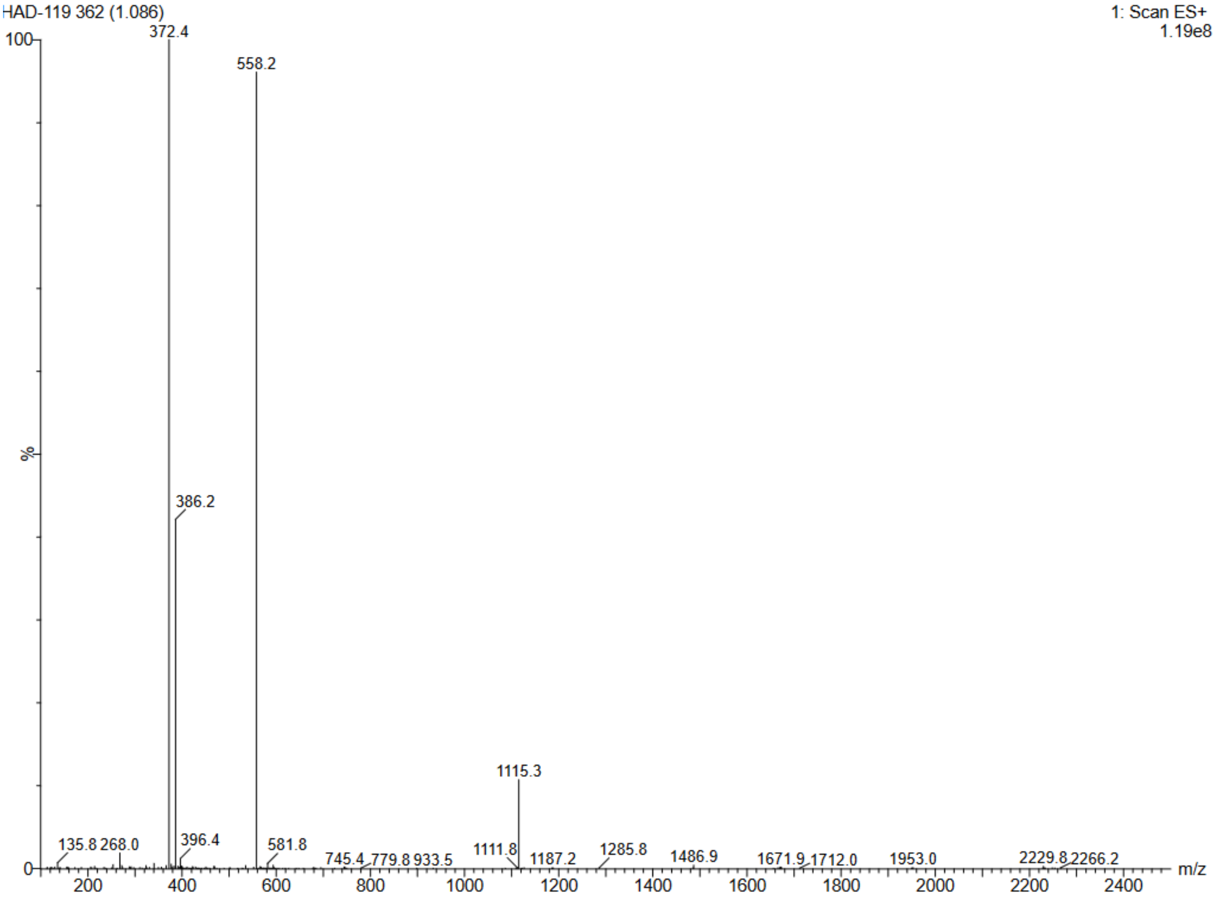

Compound 27a:D-R-V-Y-I-H-P-F(2I)

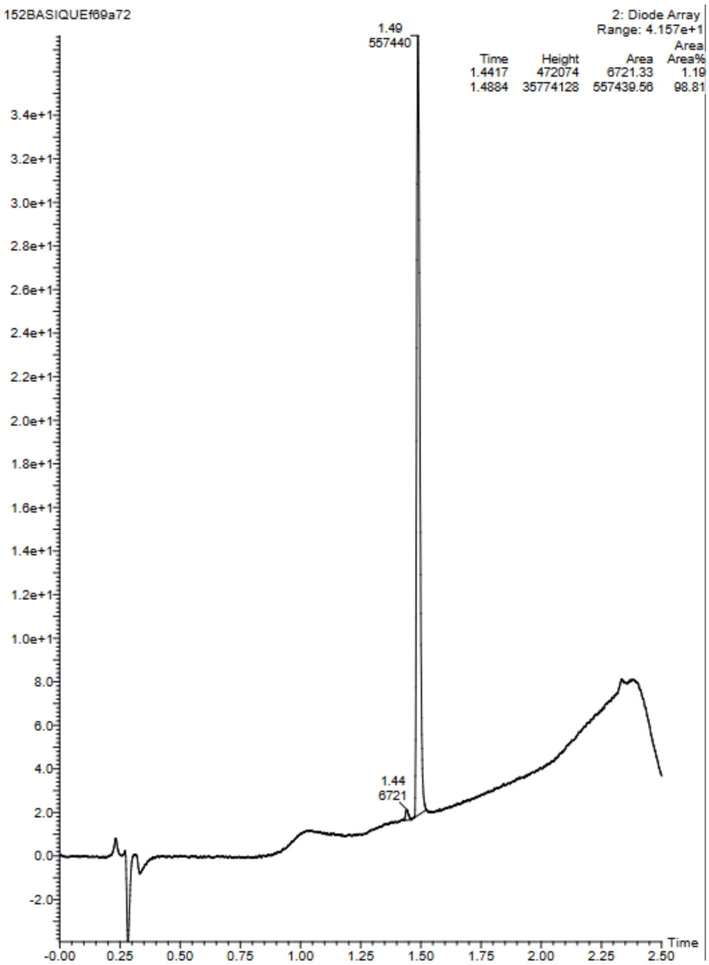


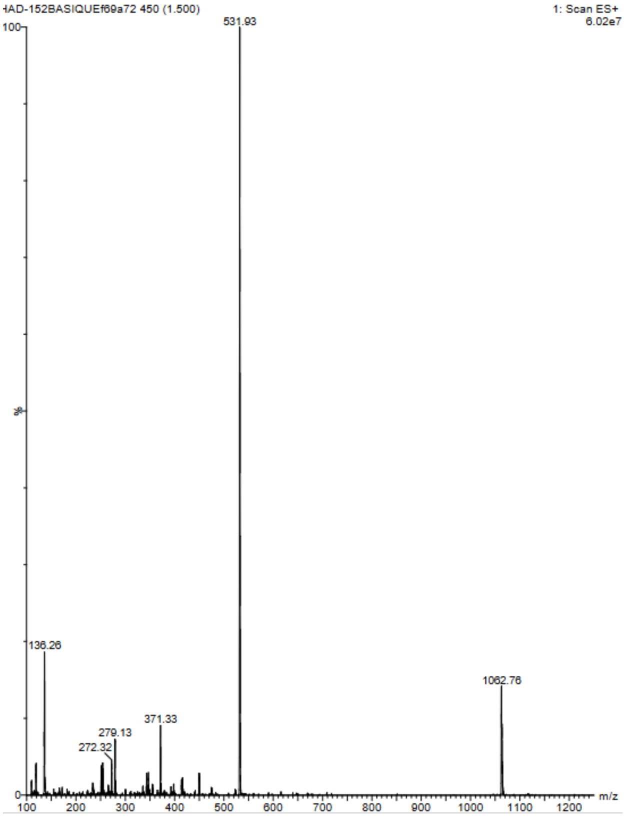

Compound 28a:D-R-V-Y-I-H-P-F(2Br)

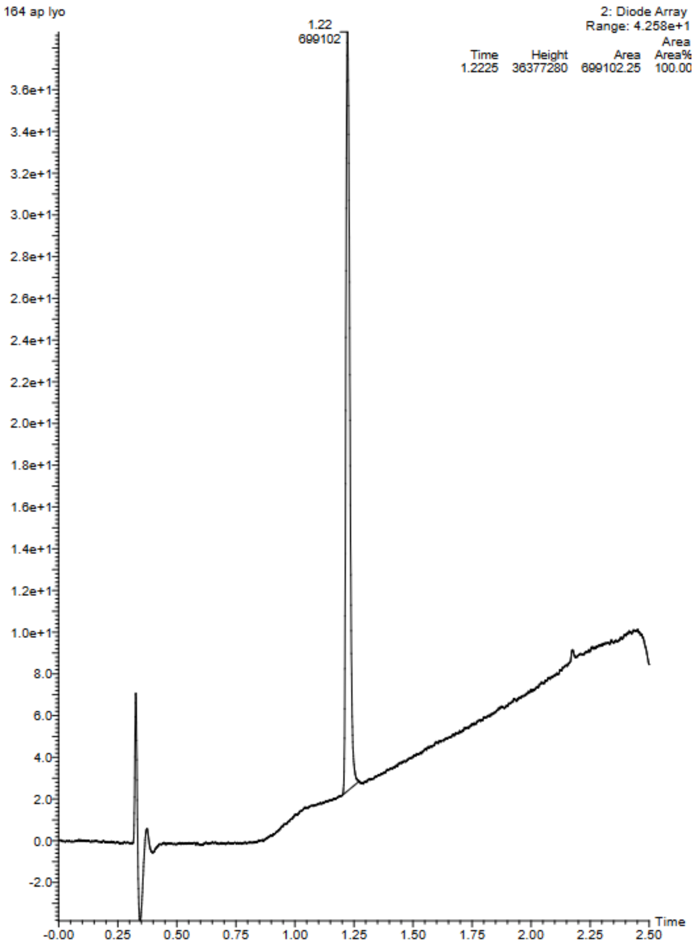


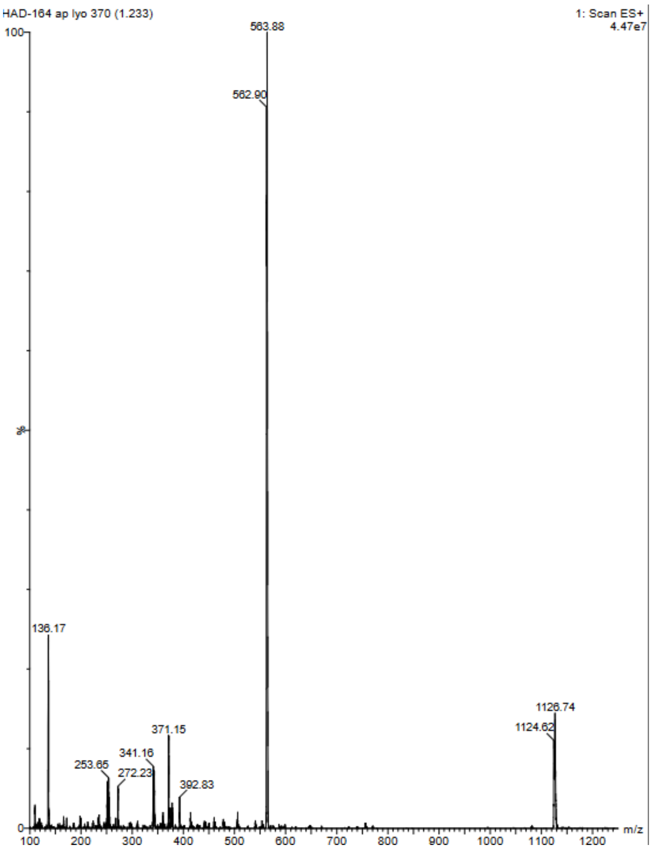

Compound 29a:D-R-V-Y-I-H-P-F(2Cl)

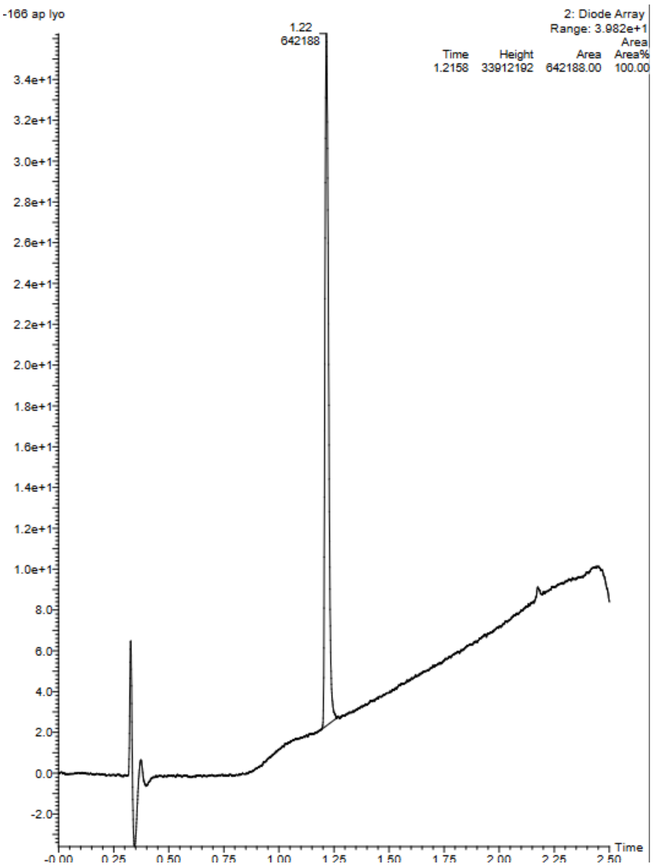


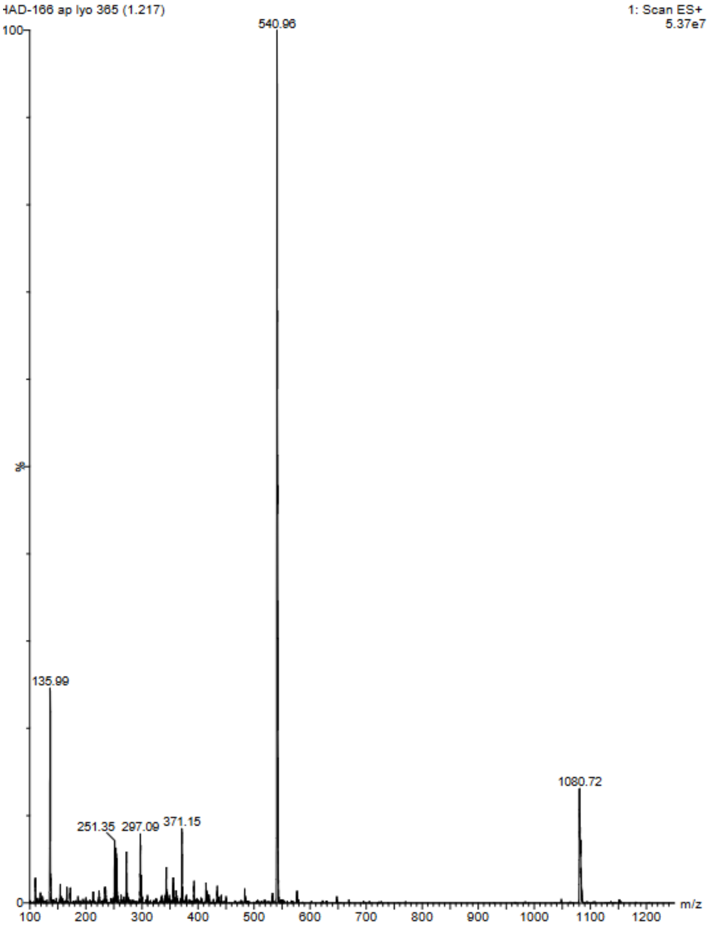

Compound 30a:D-R-V-Y-I-H-P-F(2F)

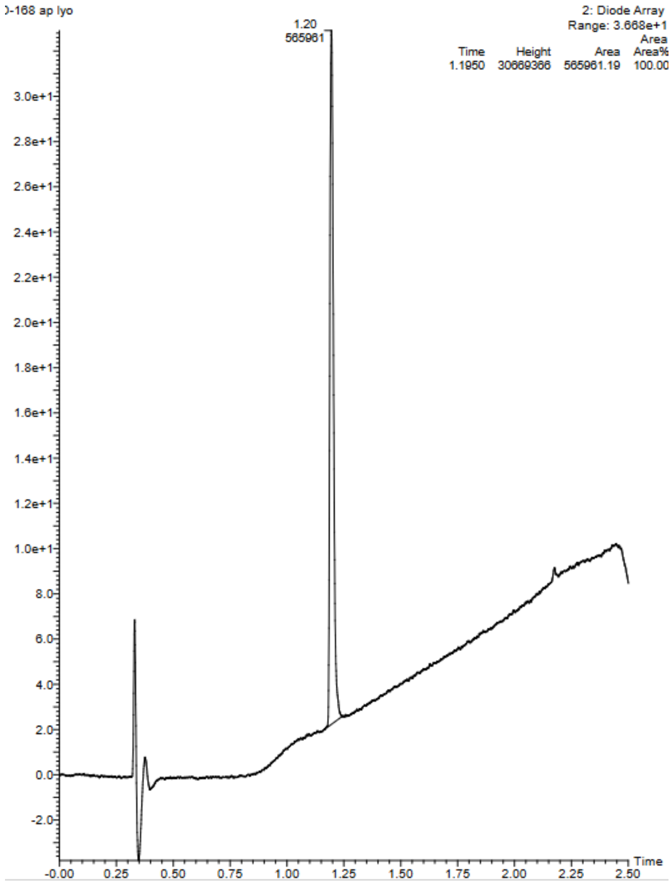


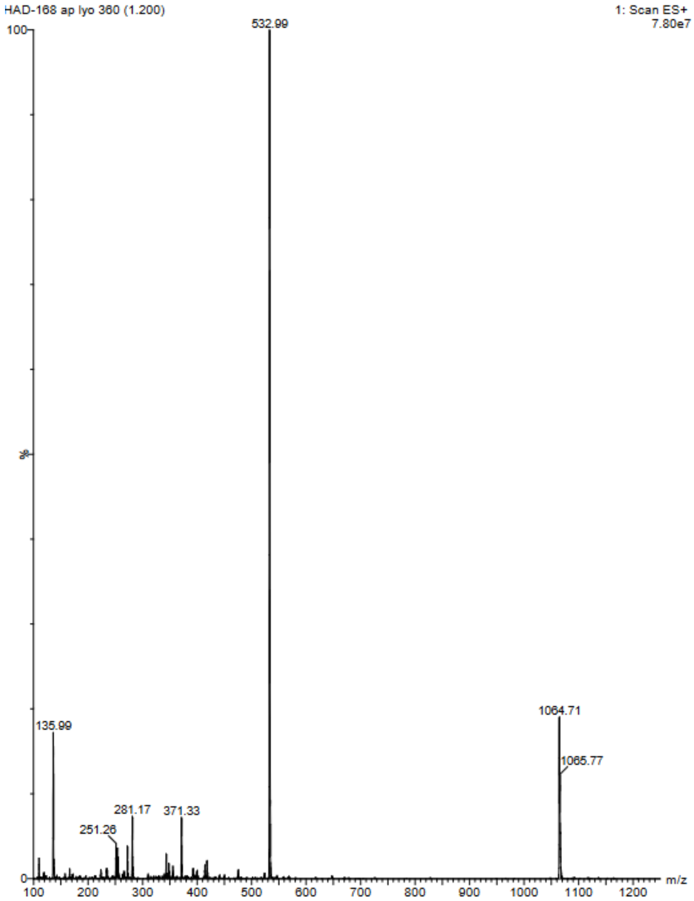

Compound 22b: D-R-V-Y-I-H-P-F(3NO2)

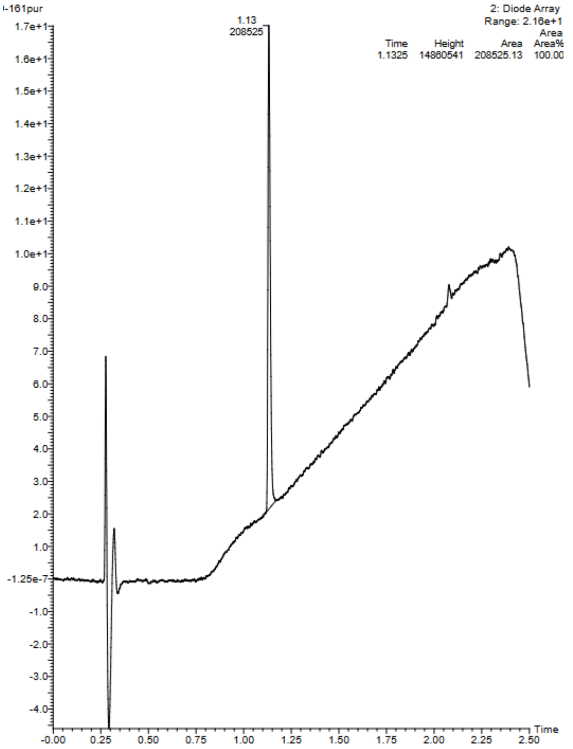


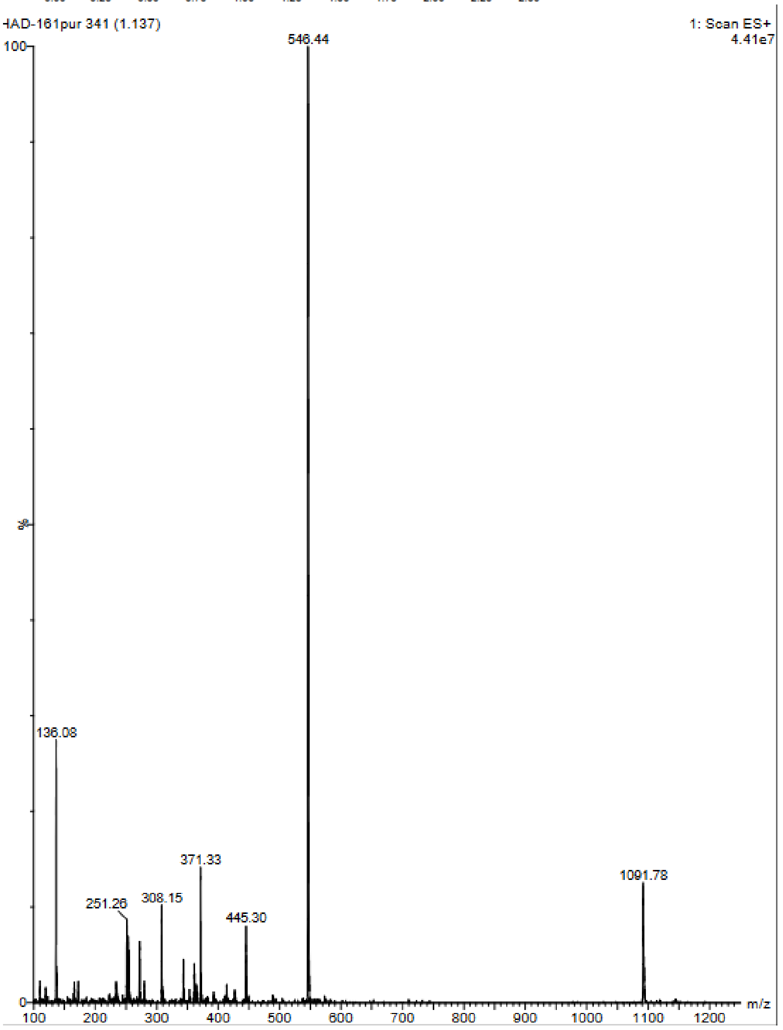

Compound 23b: D-R-V-Y-I-H-P-F(3CN)

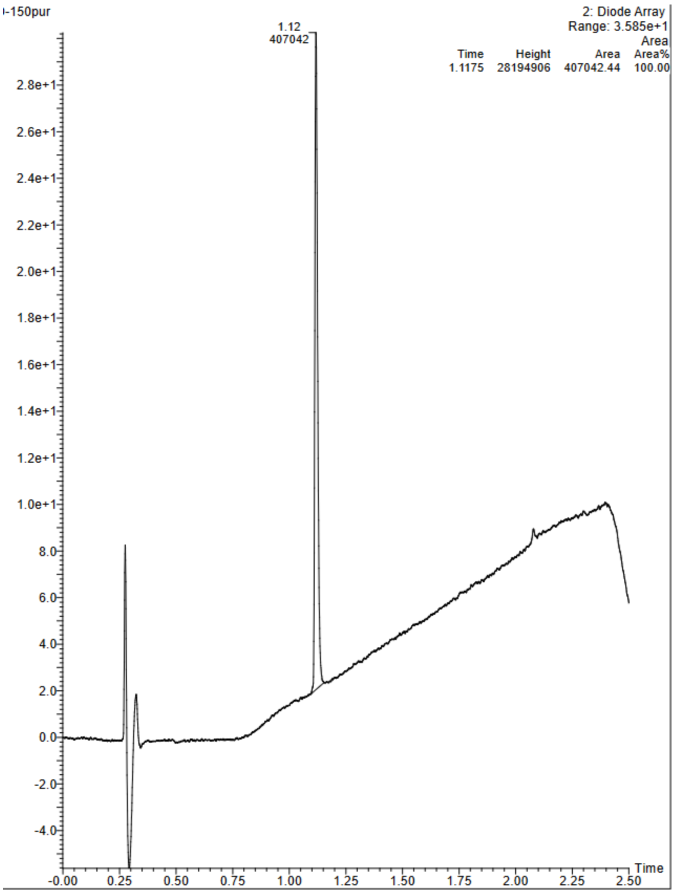


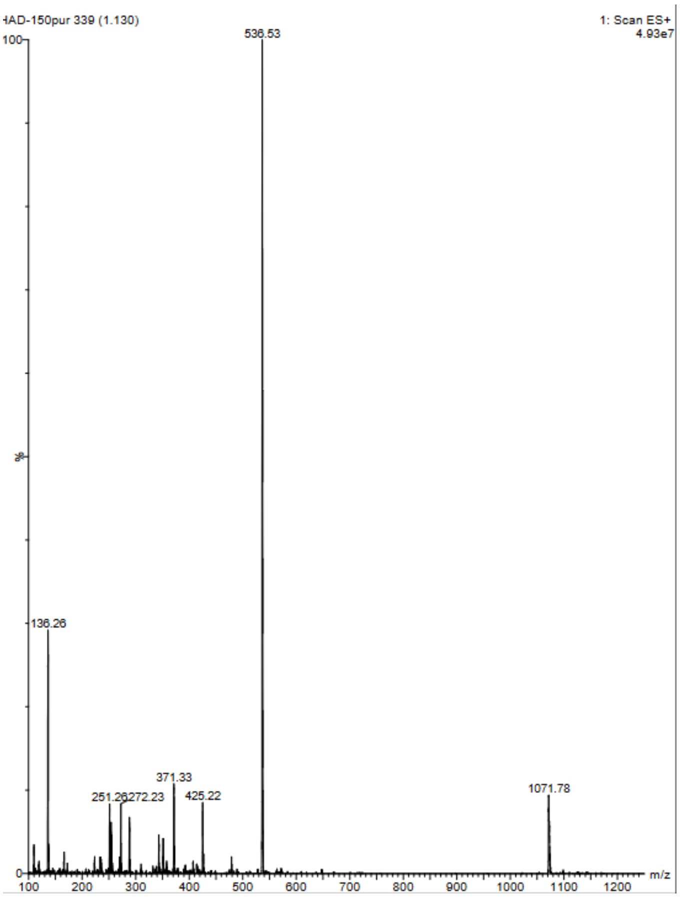

Compound 24b: D-R-V-Y-I-H-P-F(3OH)

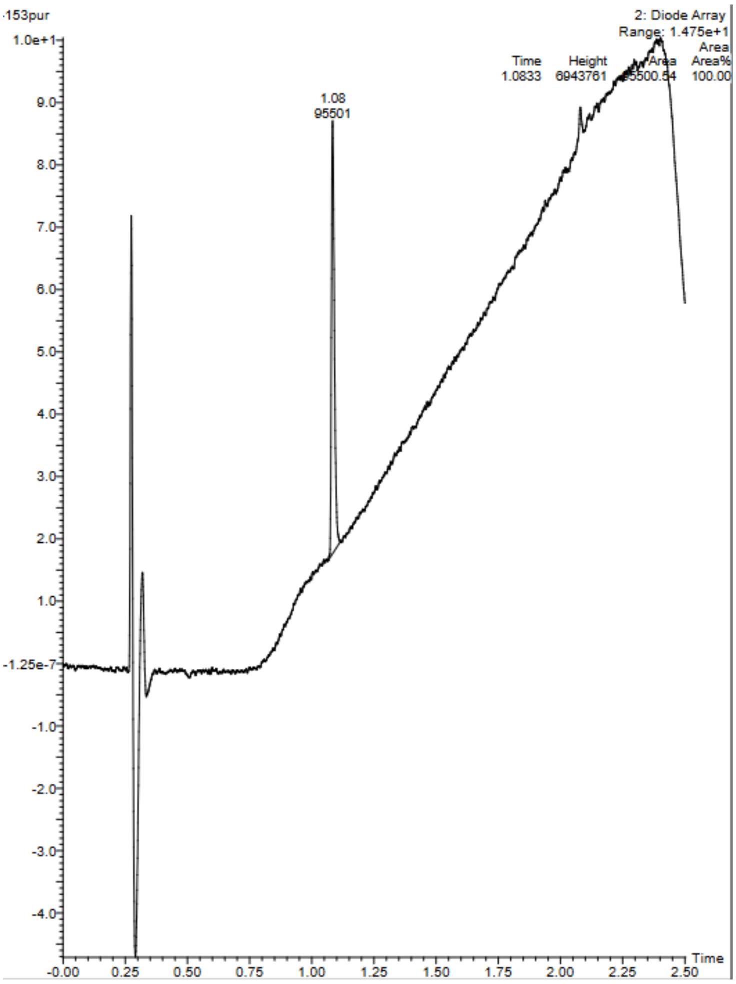


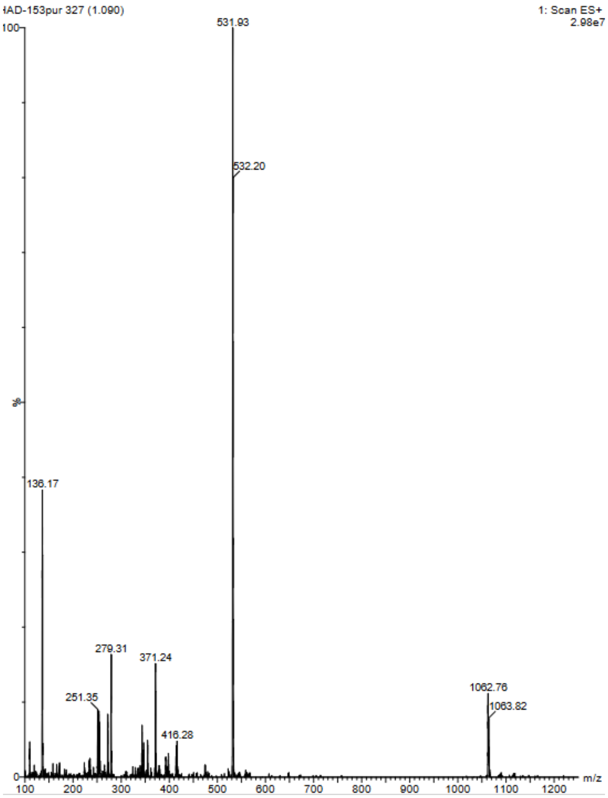

Compound 25b: D-R-V-Y-I-H-P-F(3Me)

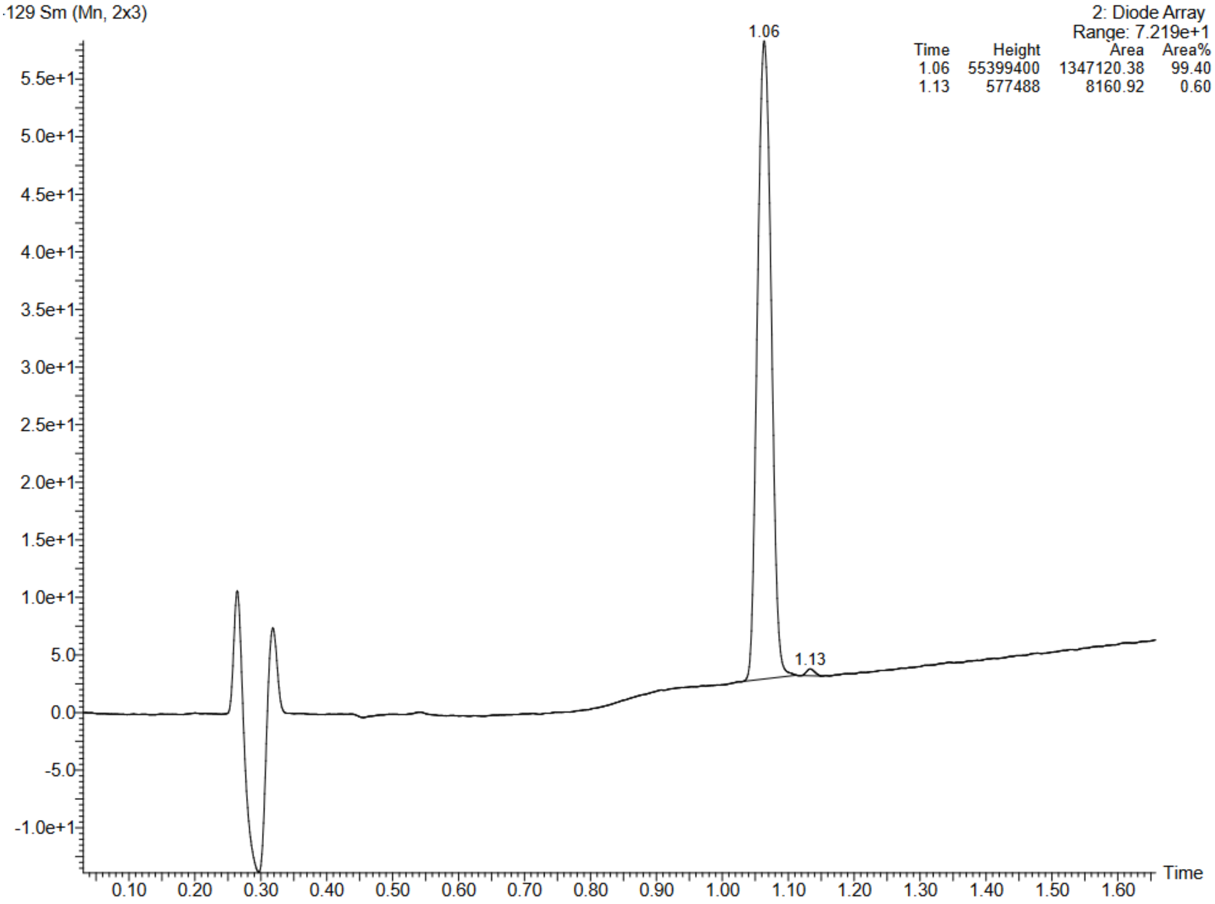


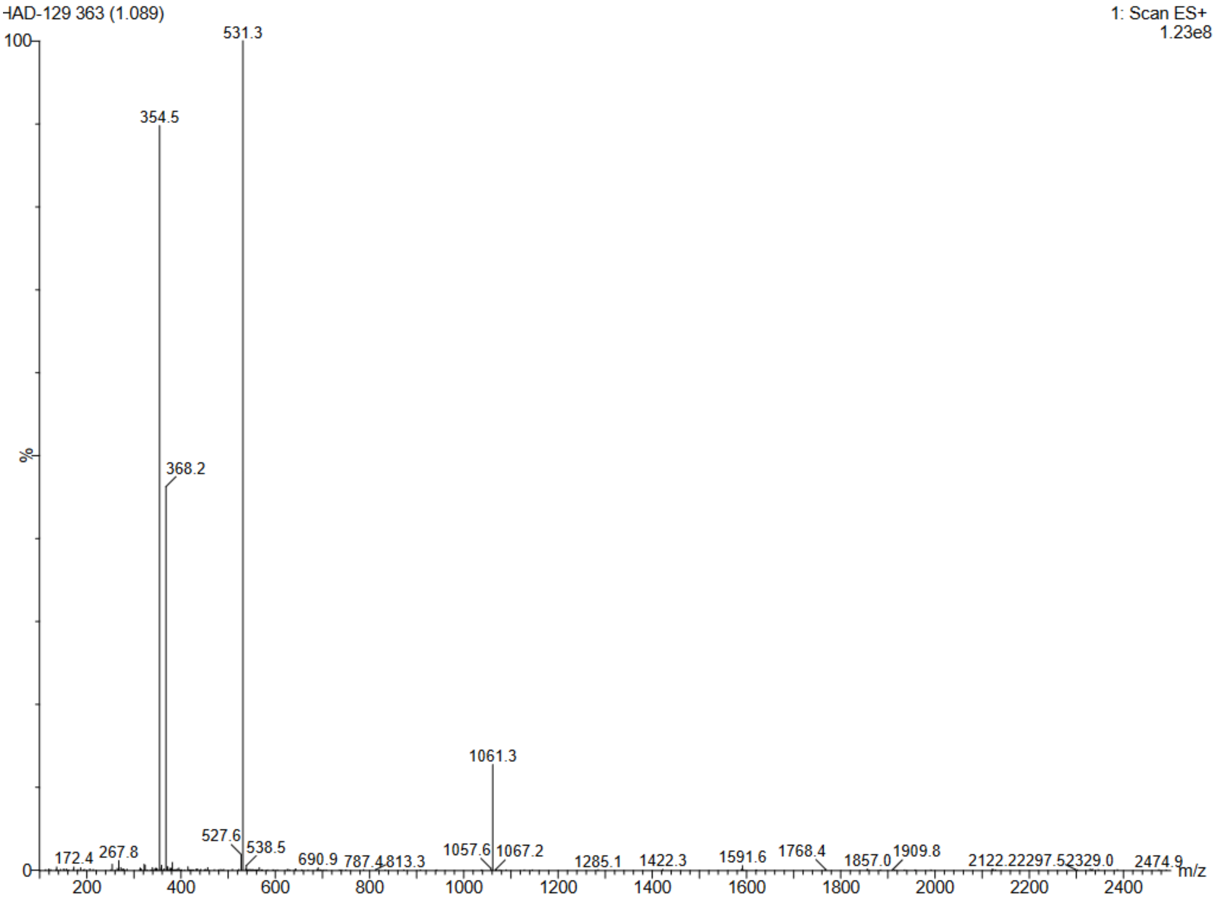

Compound 26b: D-R-V-Y-I-H-P-F(3CF3)

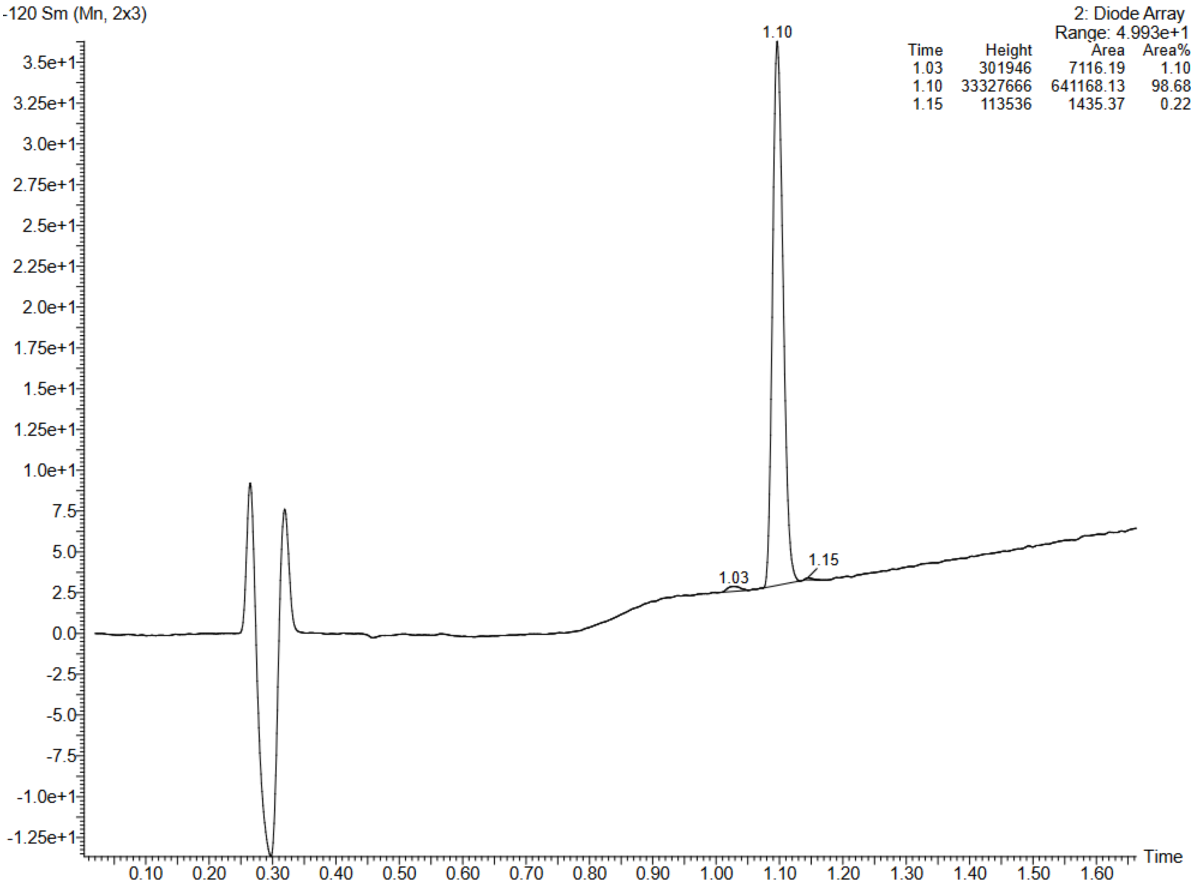


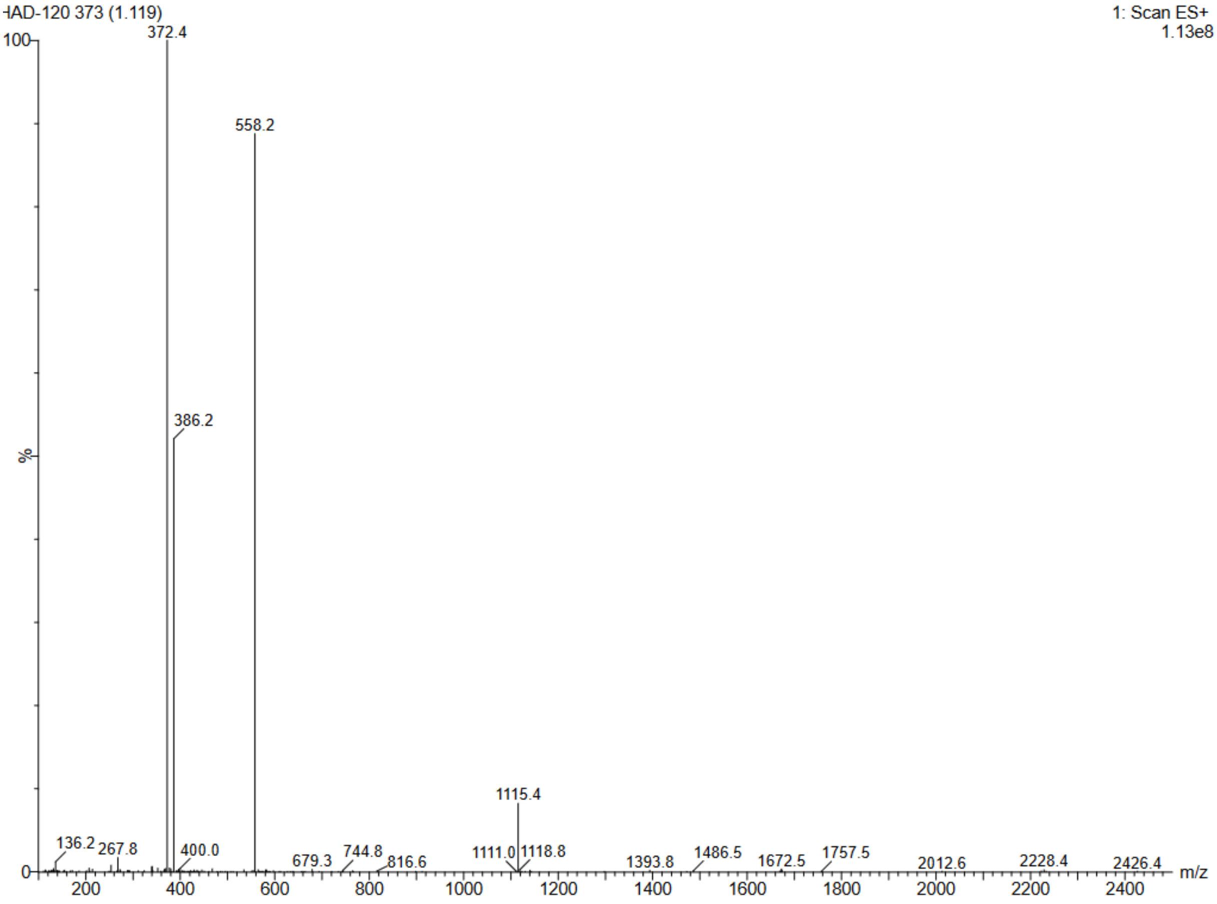

Compound 27b: D-R-V-Y-I-H-P-F(3I)

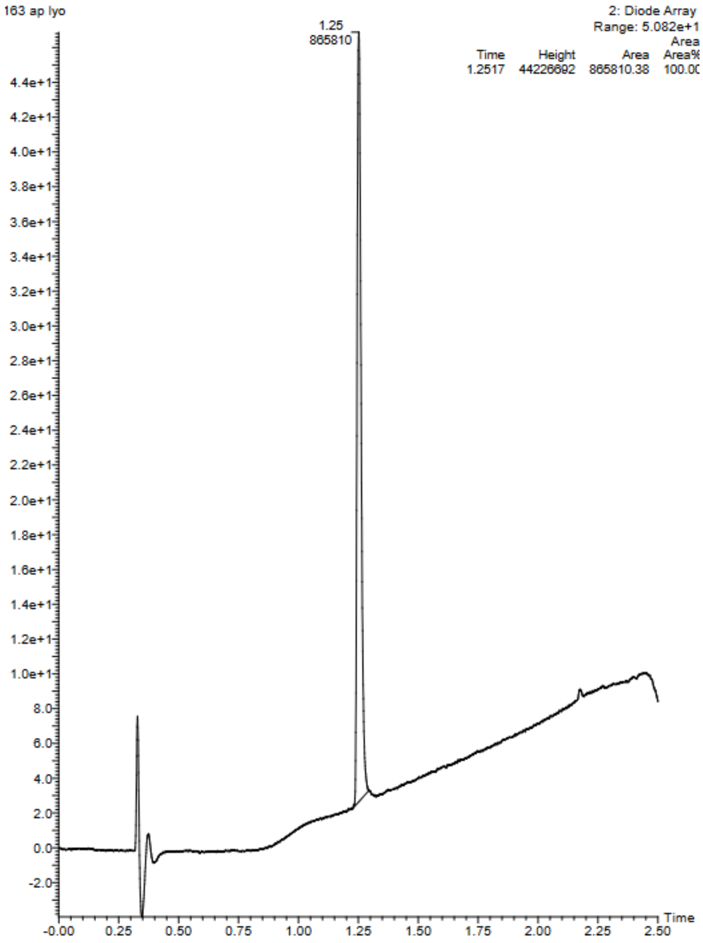


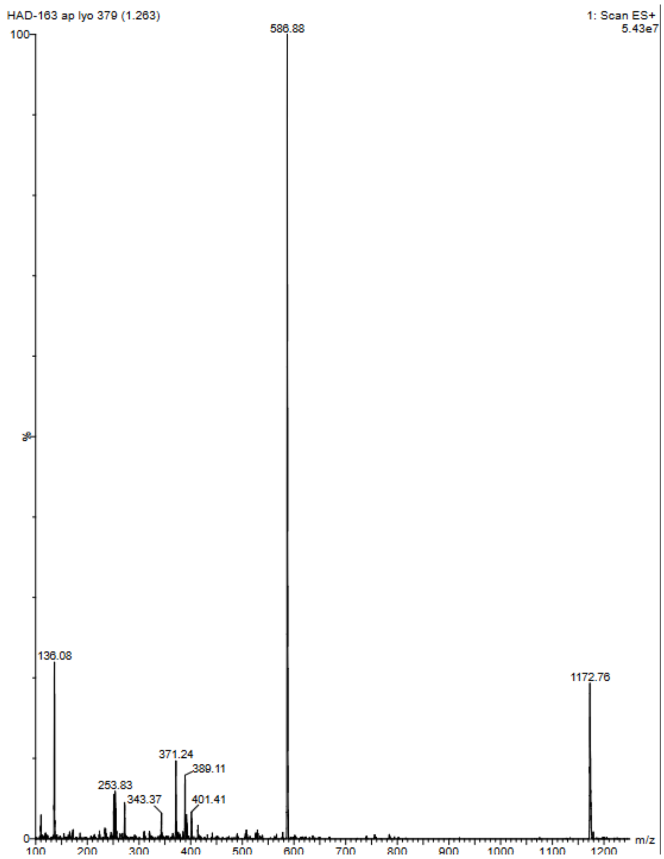

Compound 28b: D-R-V-Y-I-H-P-F(3Br)

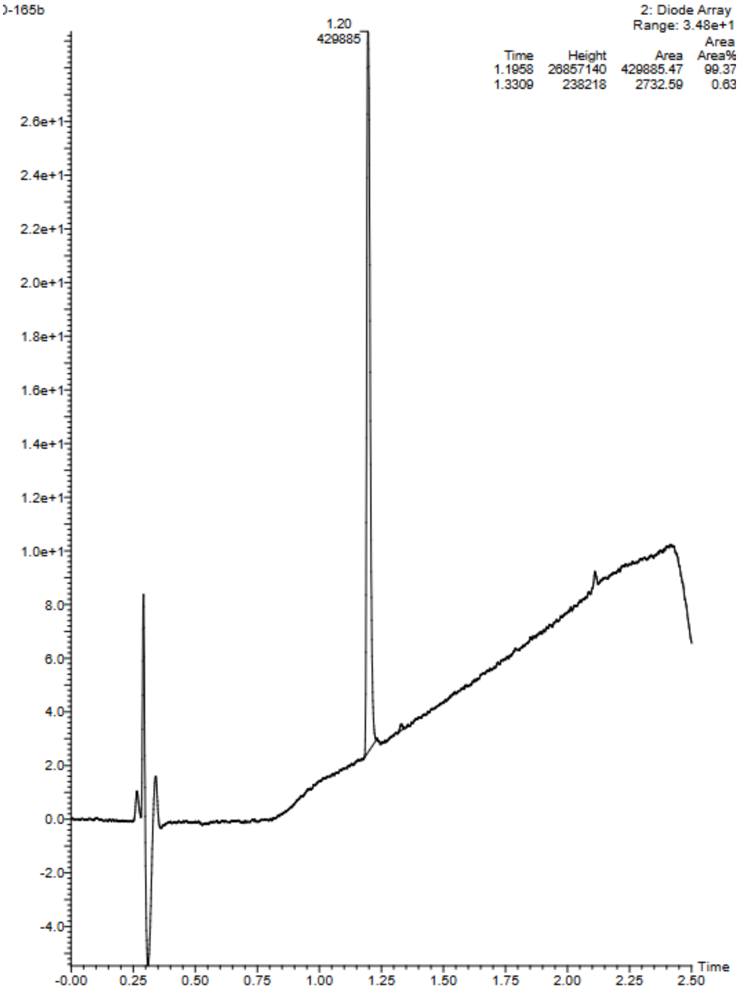


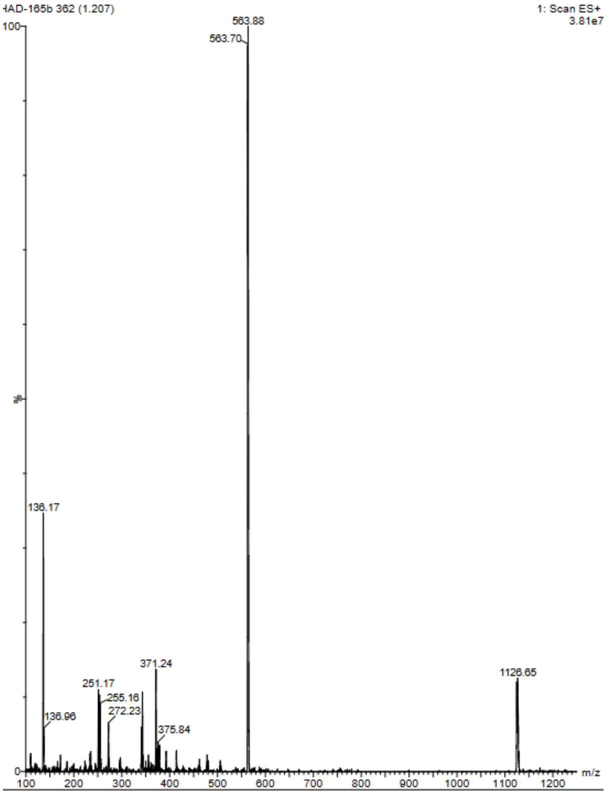

Compound 29b: D-R-V-Y-I-H-P-F(3Cl)

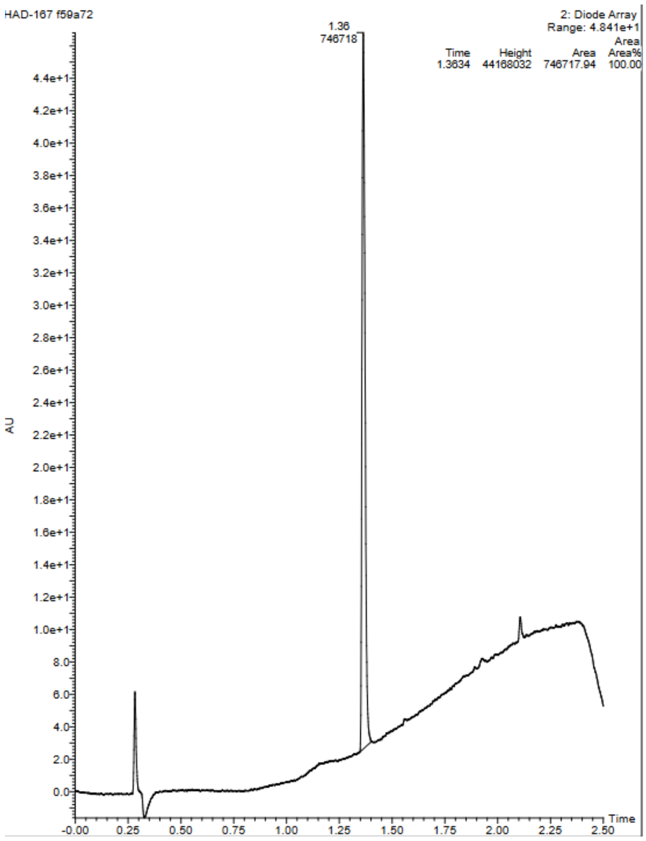


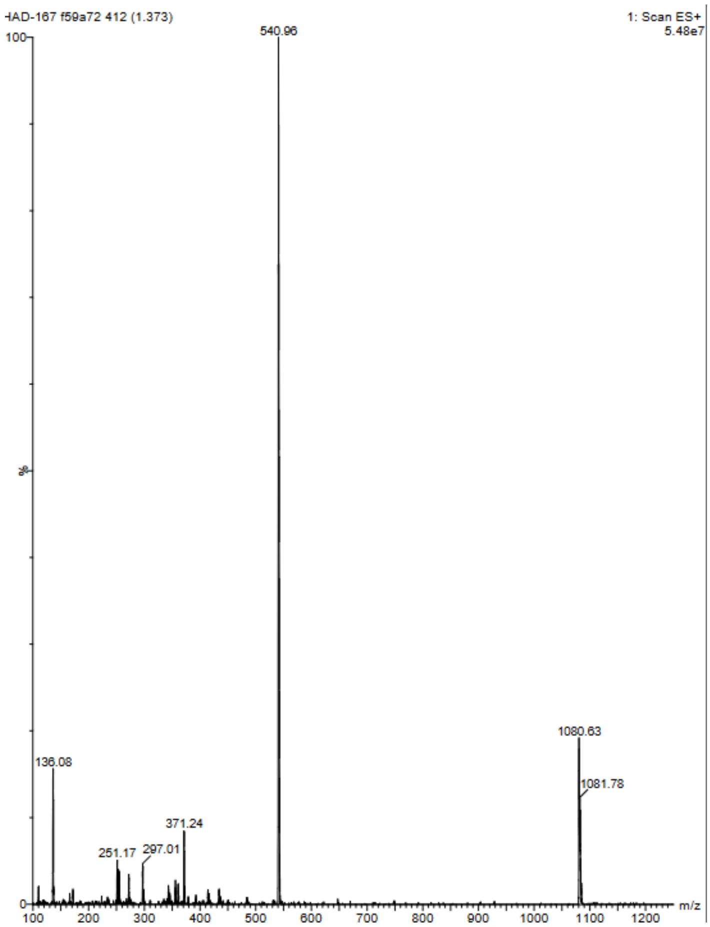

Compound 30b: D-R-V-Y-I-H-P-F(3F)

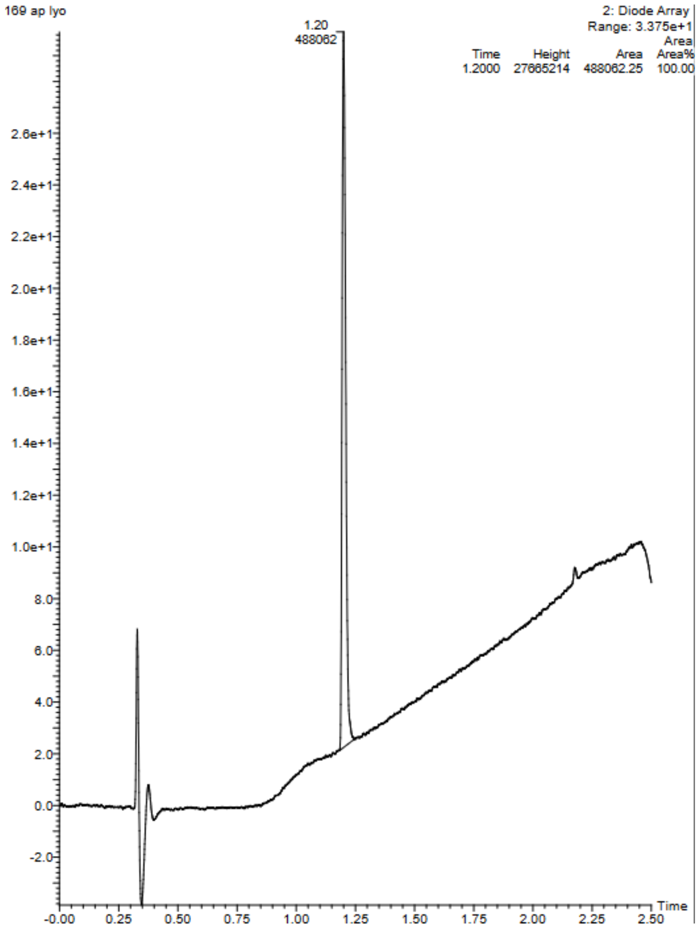


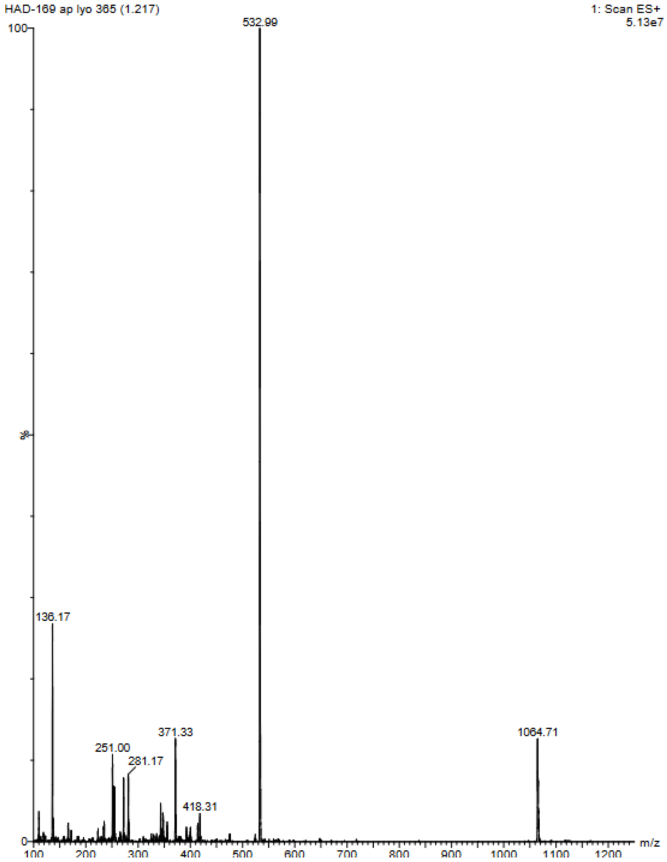

Compound 22c:D-R-V-Y-I-H-P-F(4NO2)

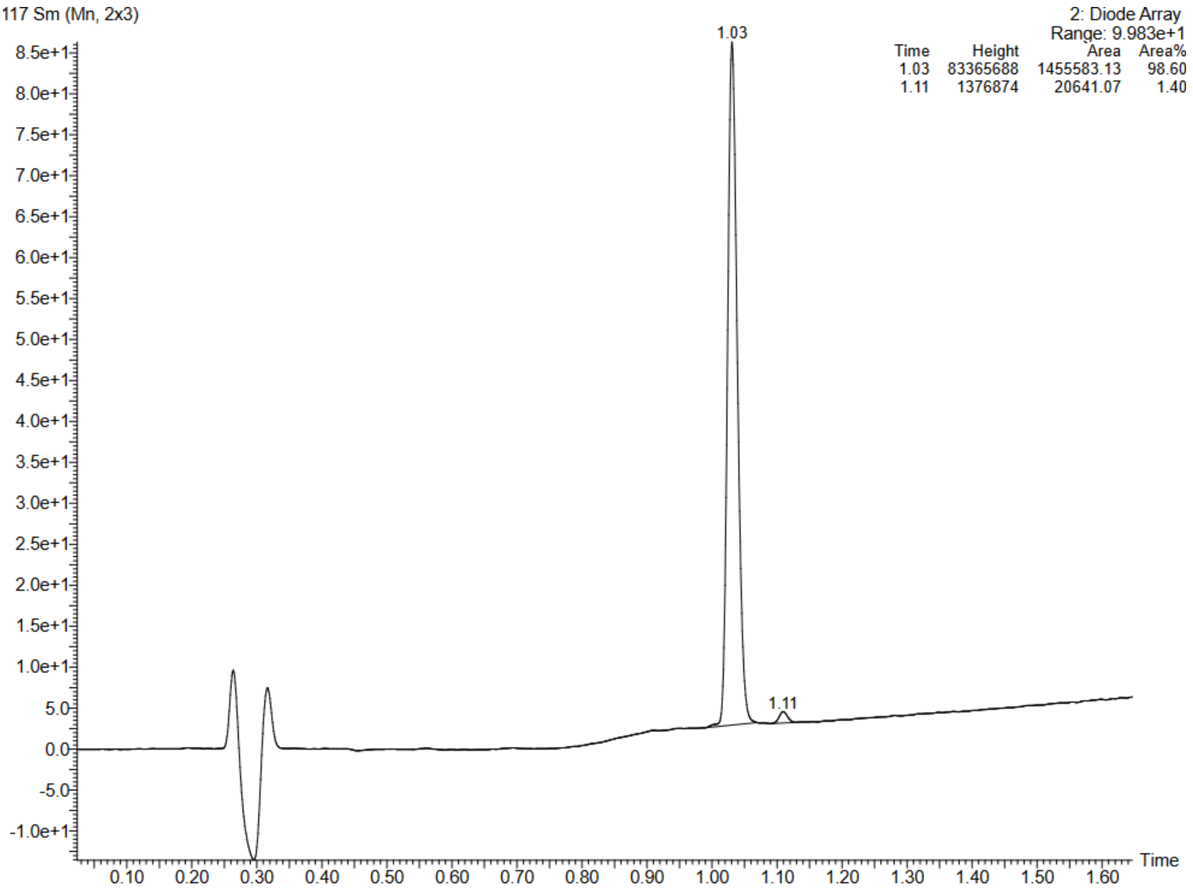


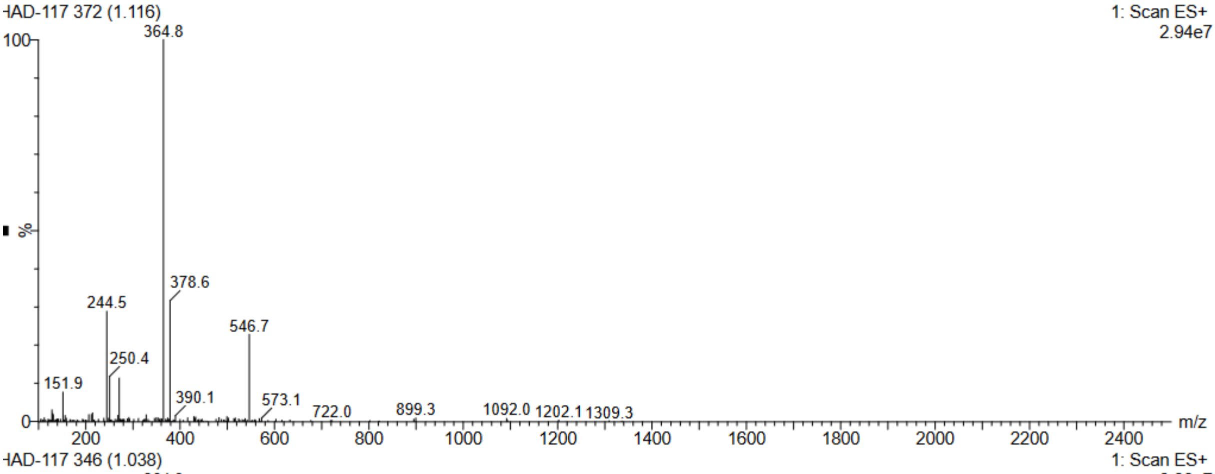


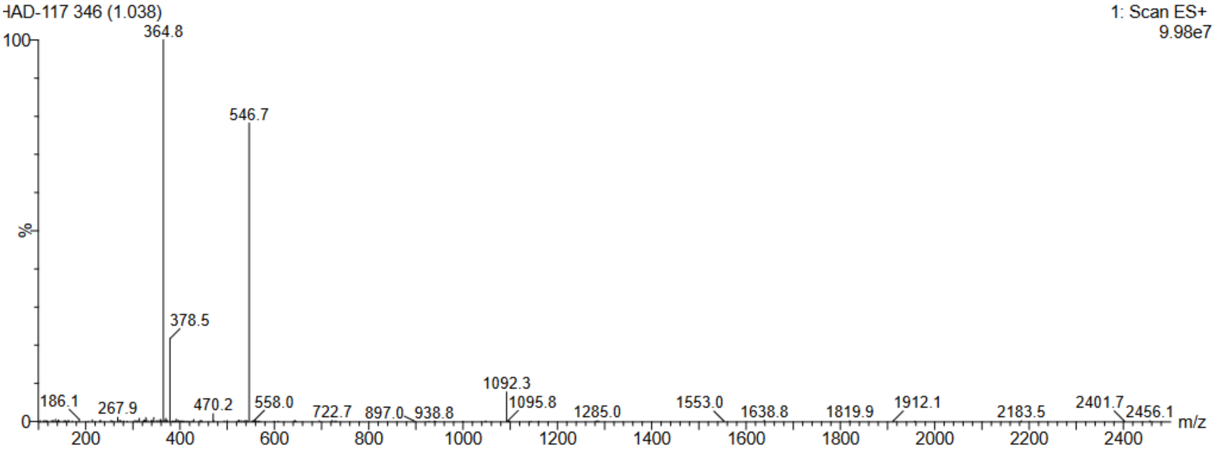

Compound 23c:D-R-V-Y-I-H-P-F(4CN)

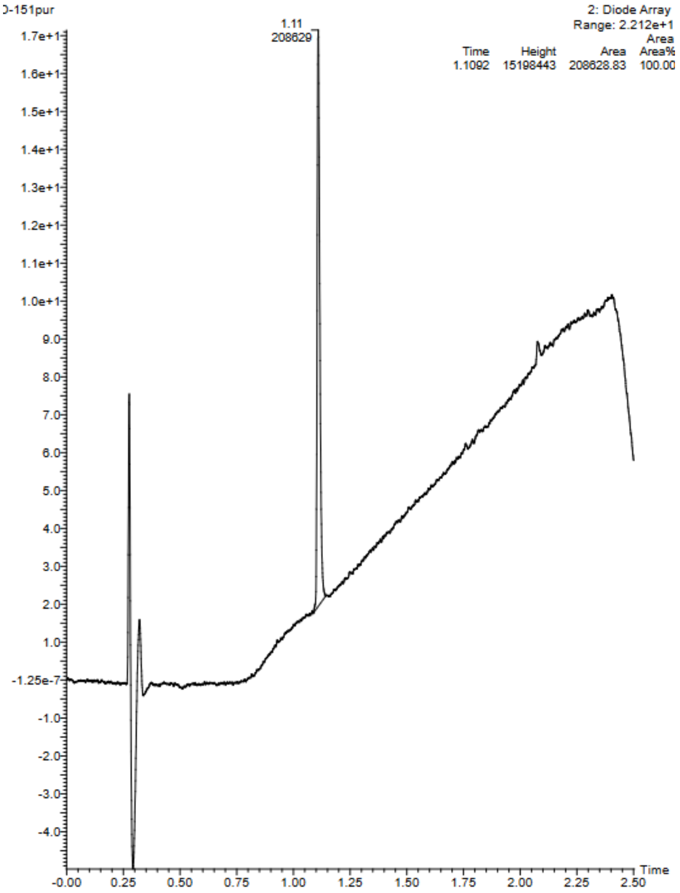


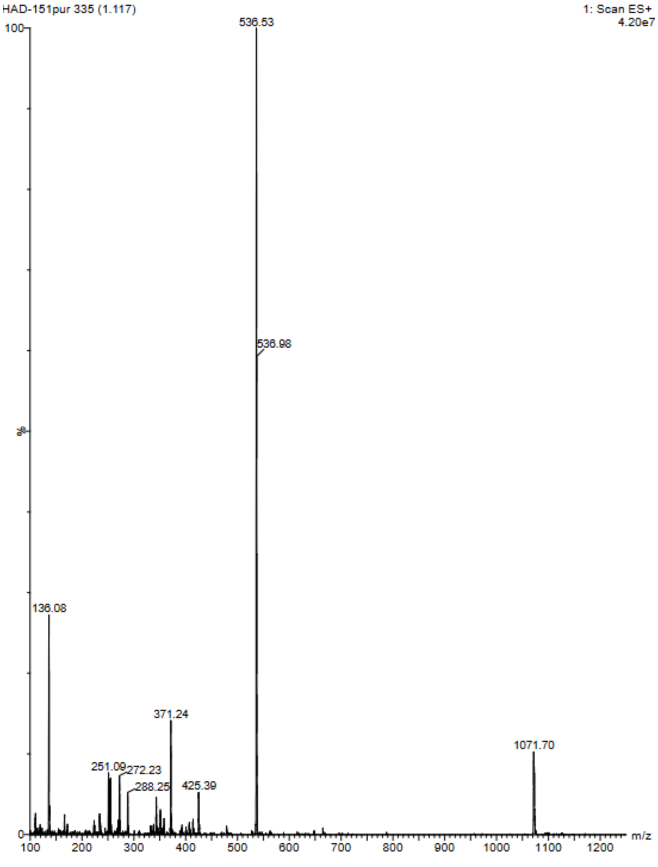

Compound 24c:D-R-V-Y-I-H-P-F(4OH)

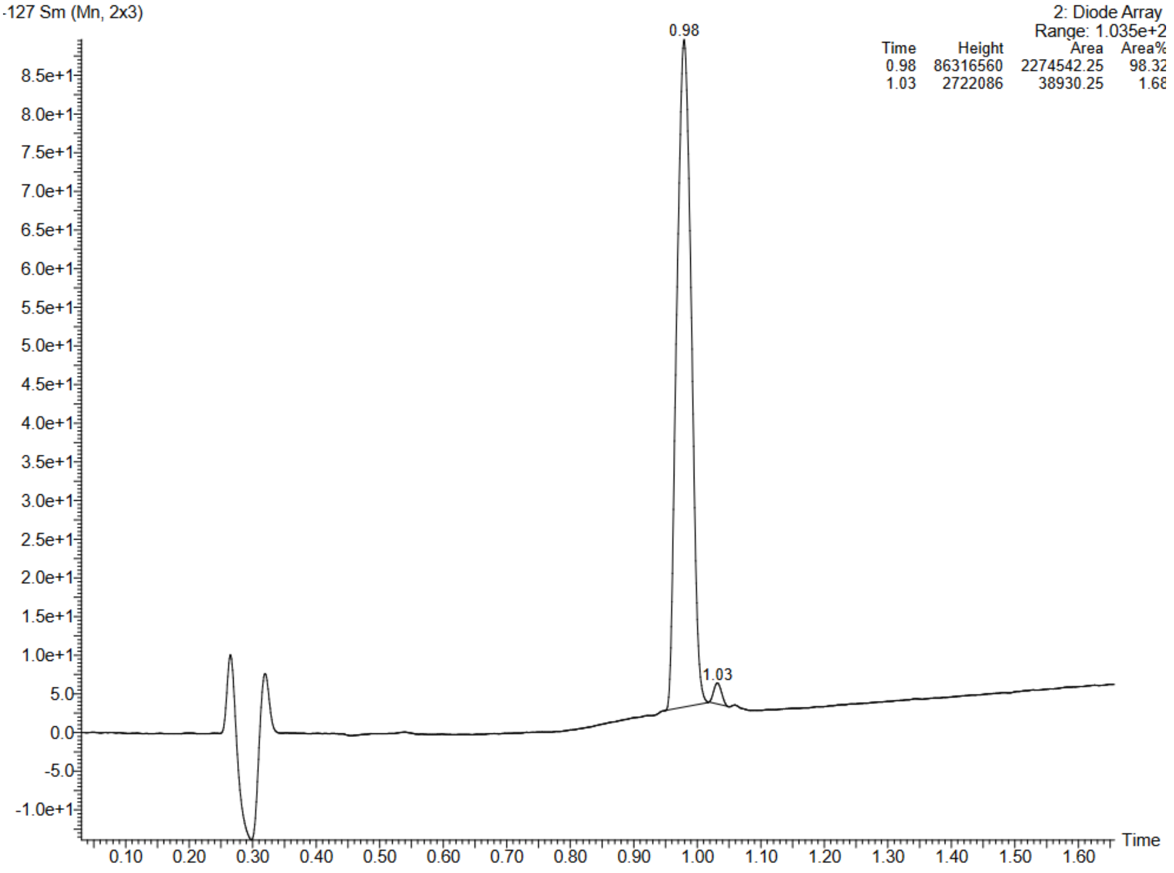


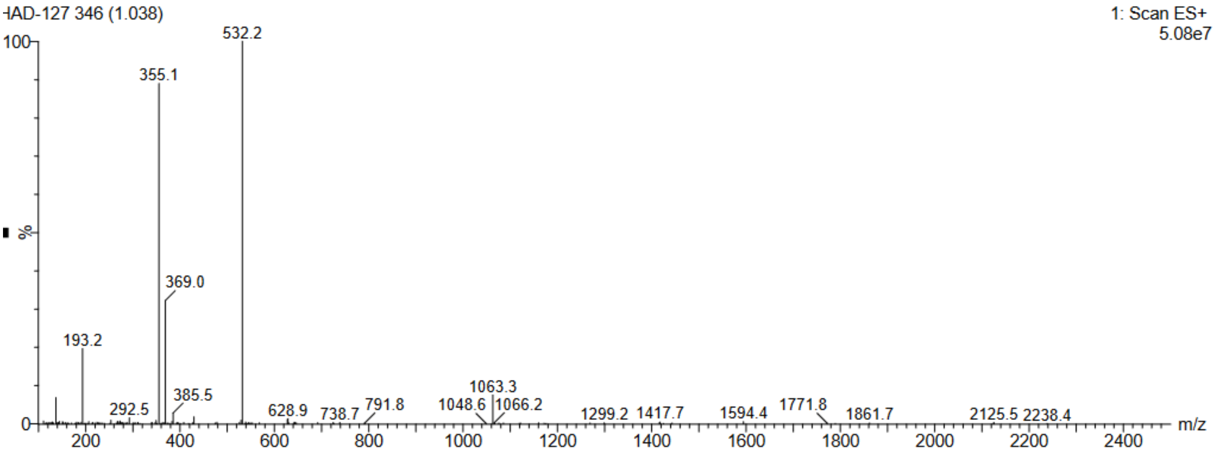


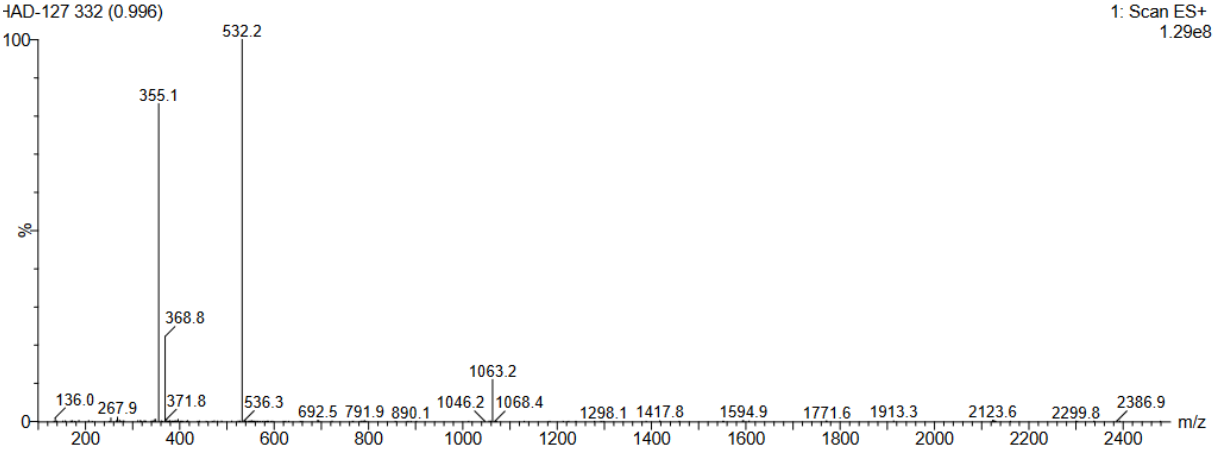

Compound 25c:D-R-V-Y-I-H-P-F(4Me)

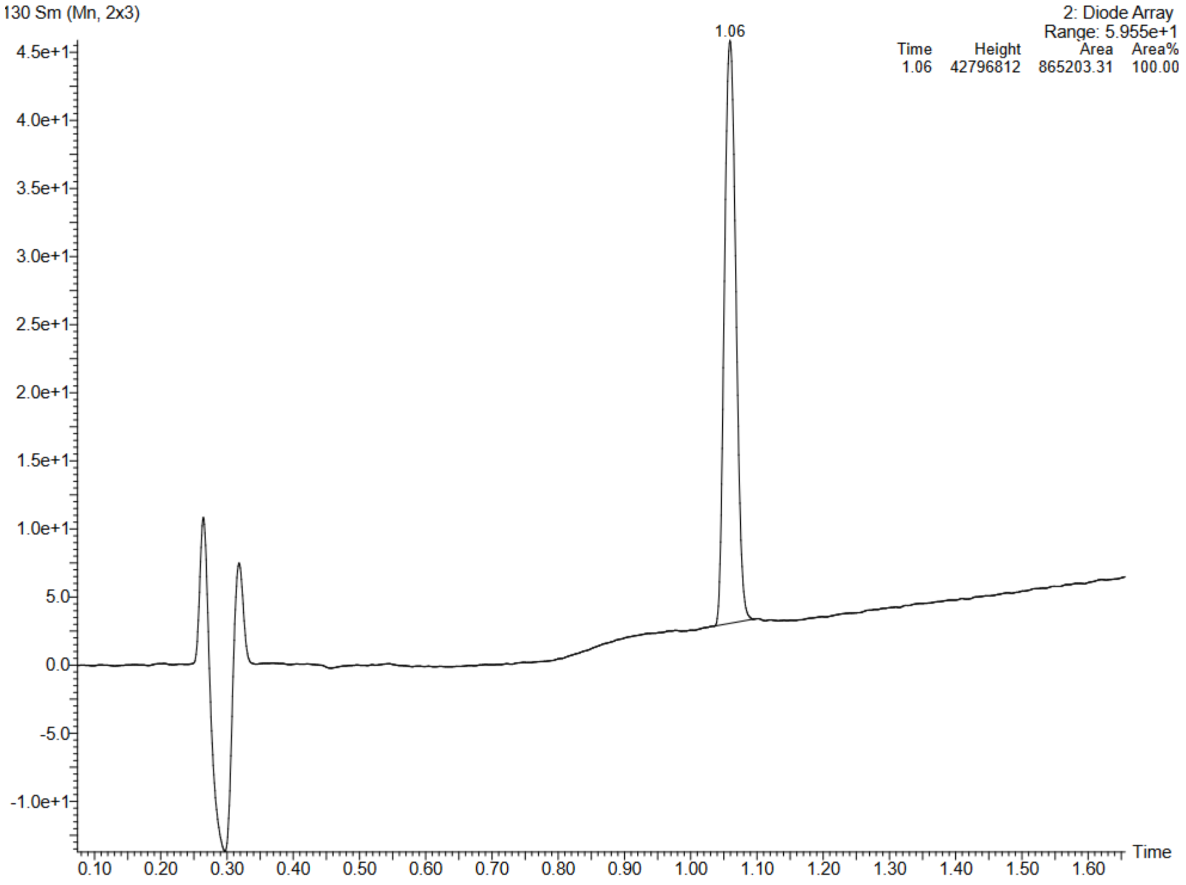


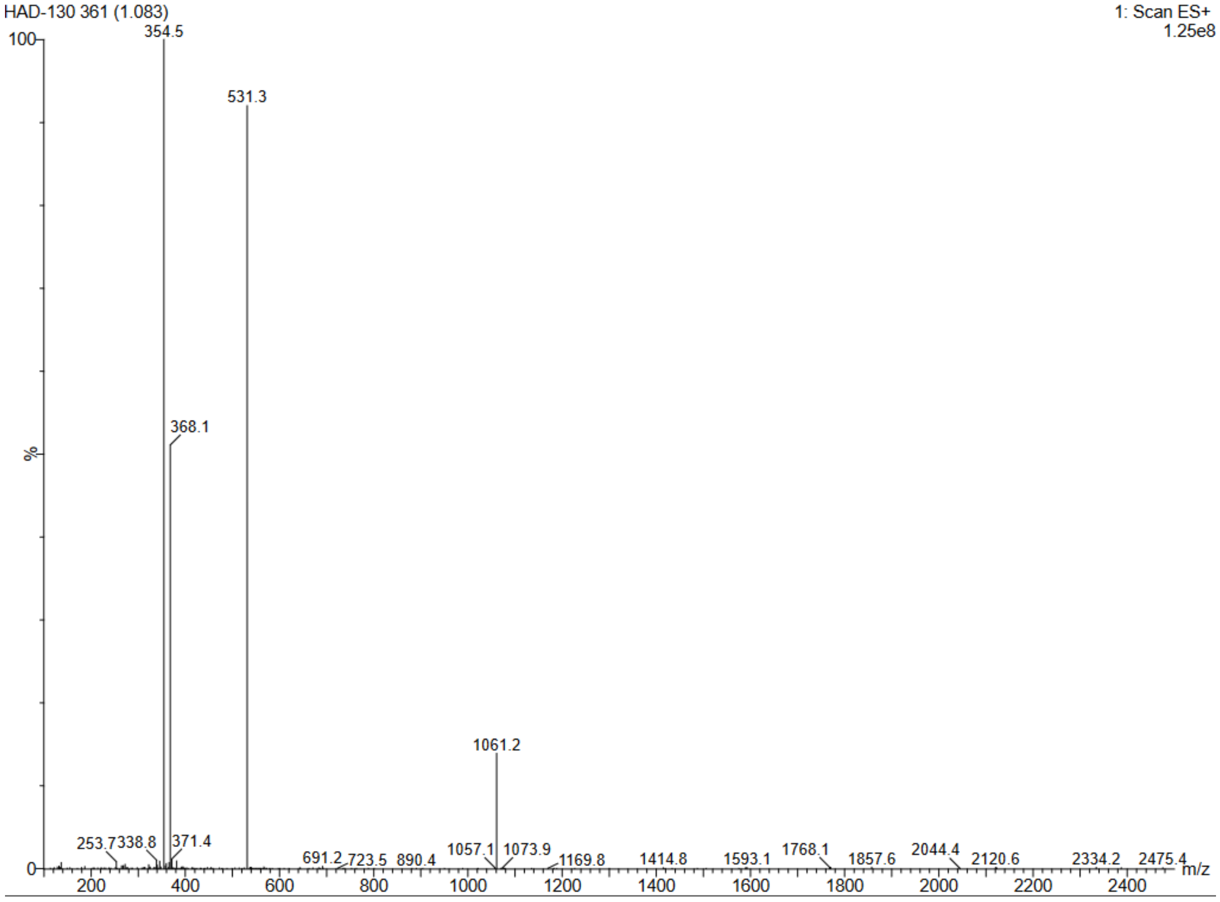

Compound 26c:D-R-V-Y-I-H-P-F(4CF3)

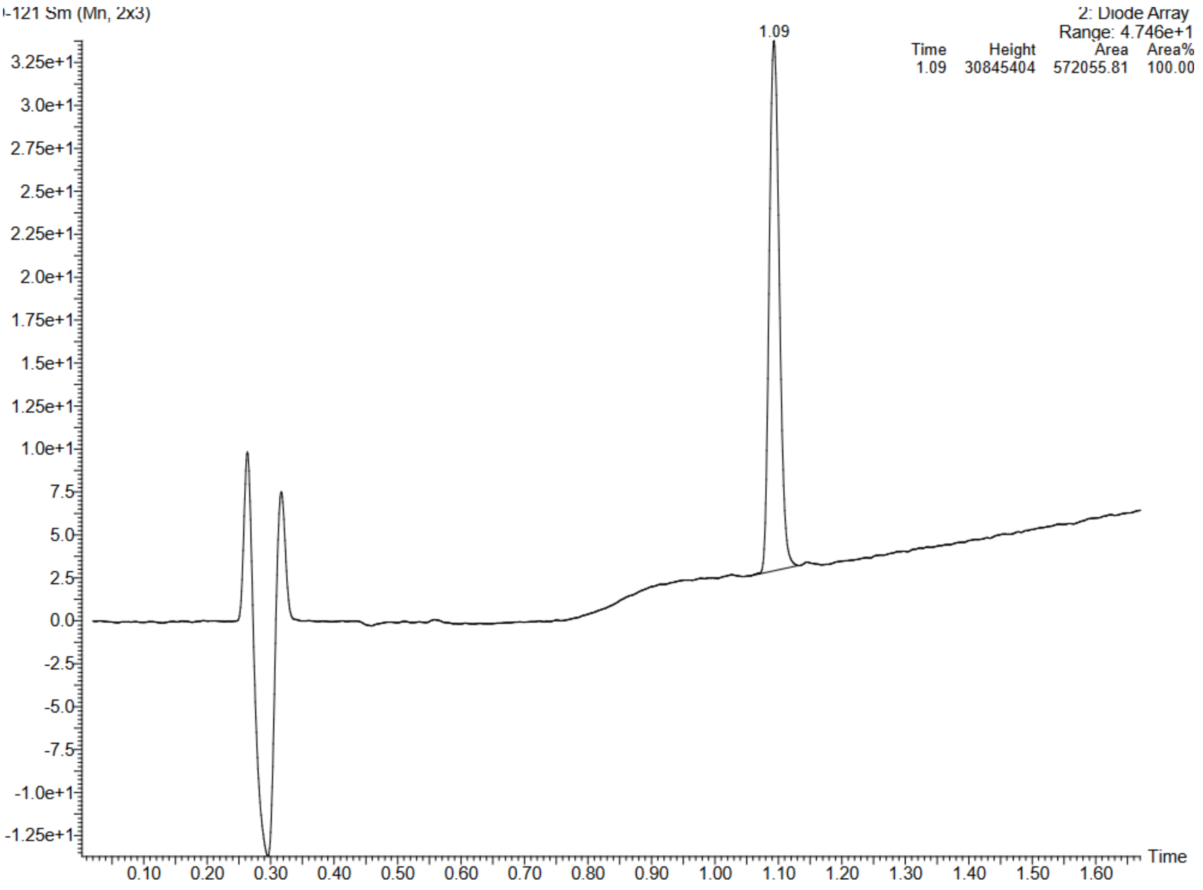


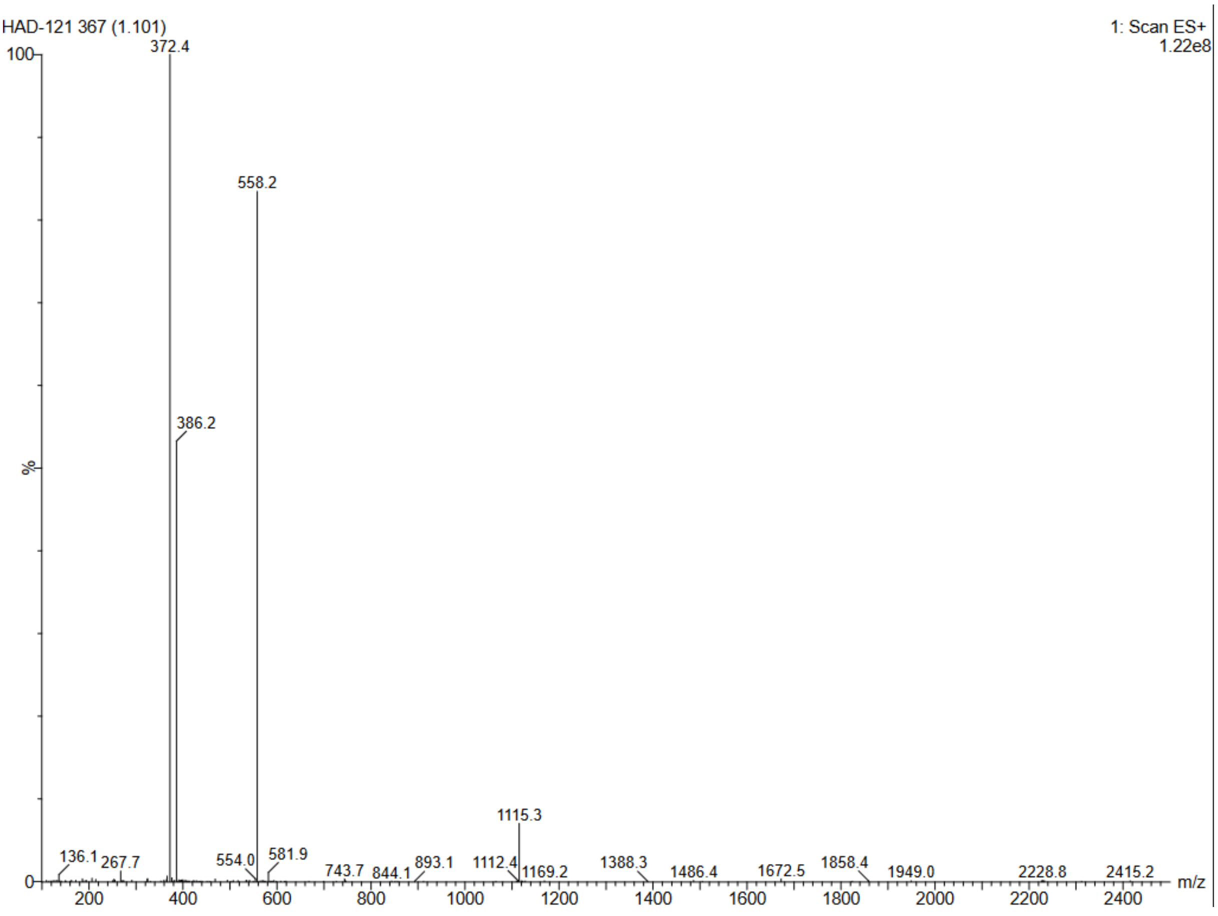

Compound 27c:D-R-V-Y-I-H-P-F(4I)

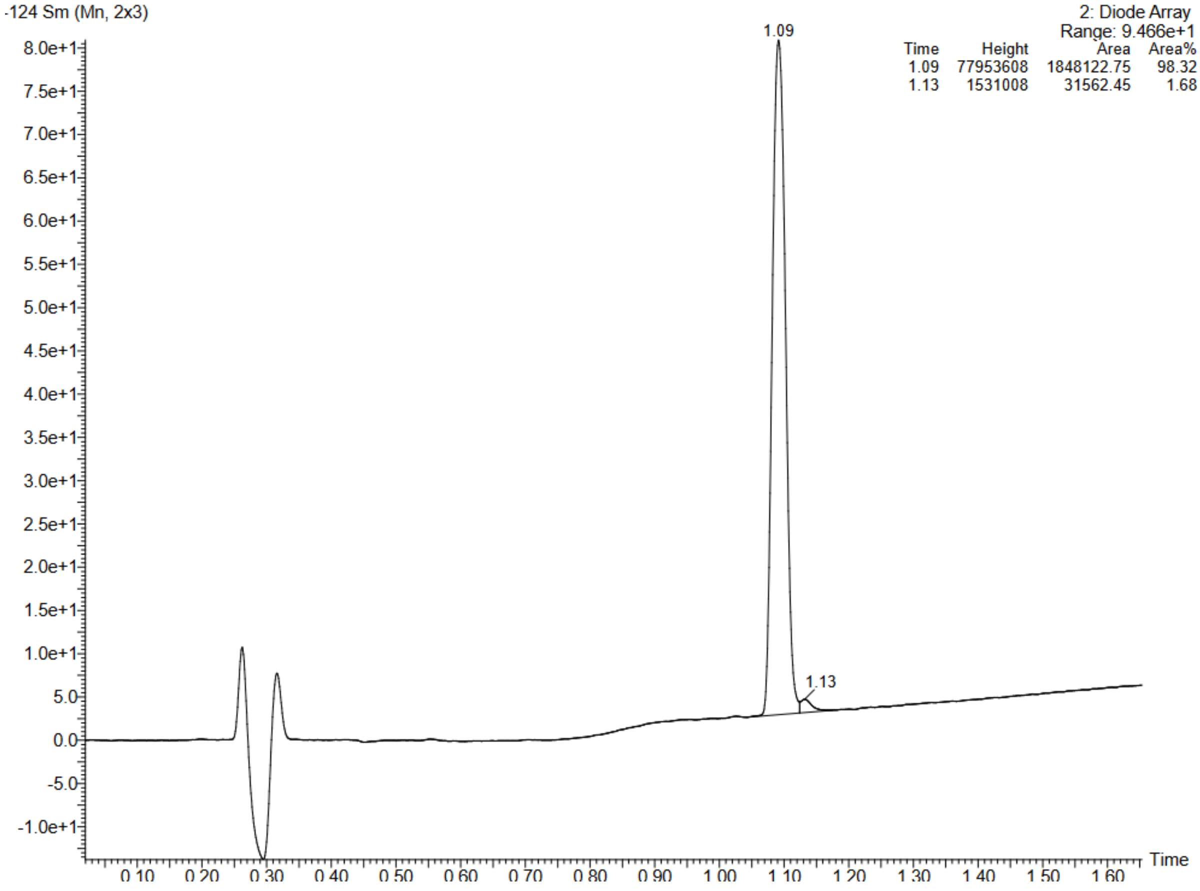


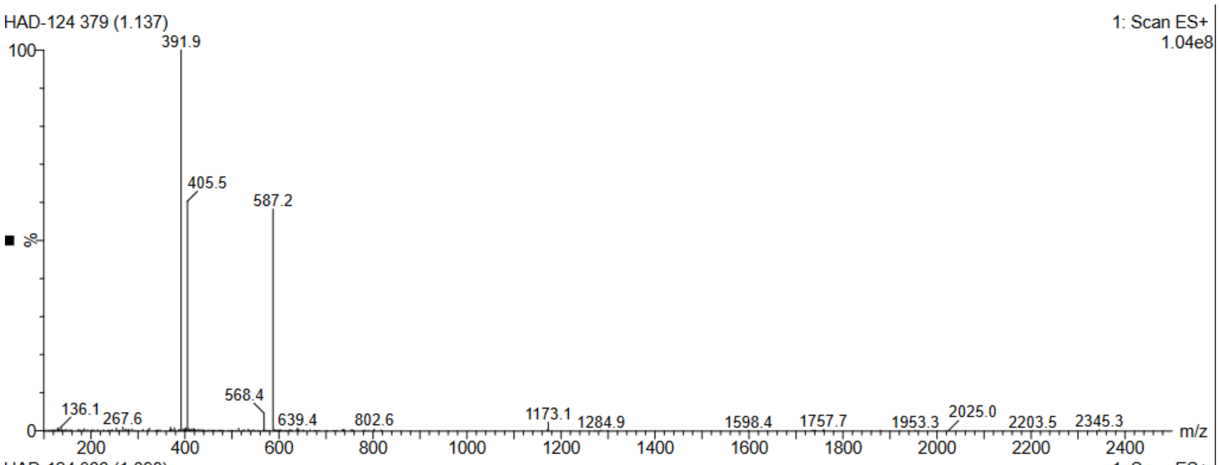


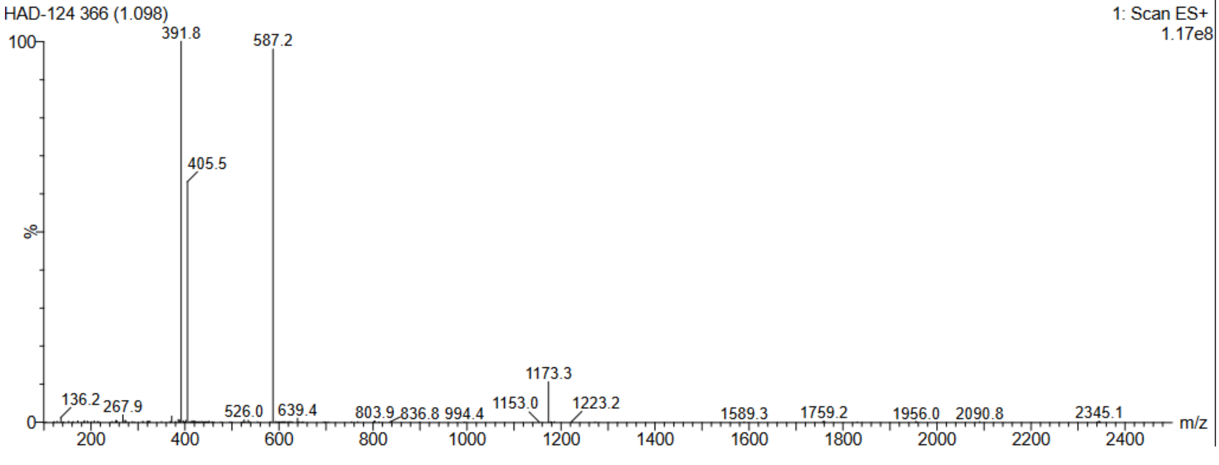

Compound 28c:D-R-V-Y-I-H-P-F(4Br)

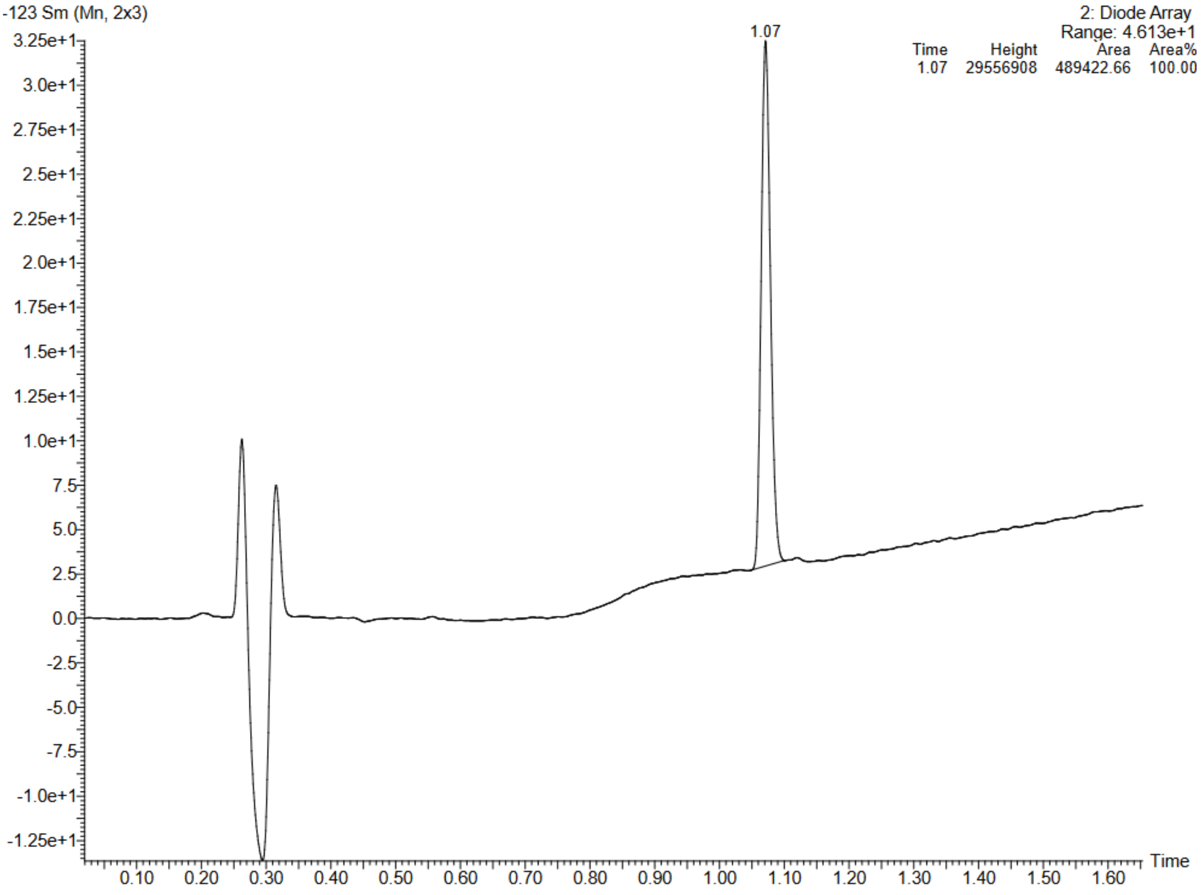


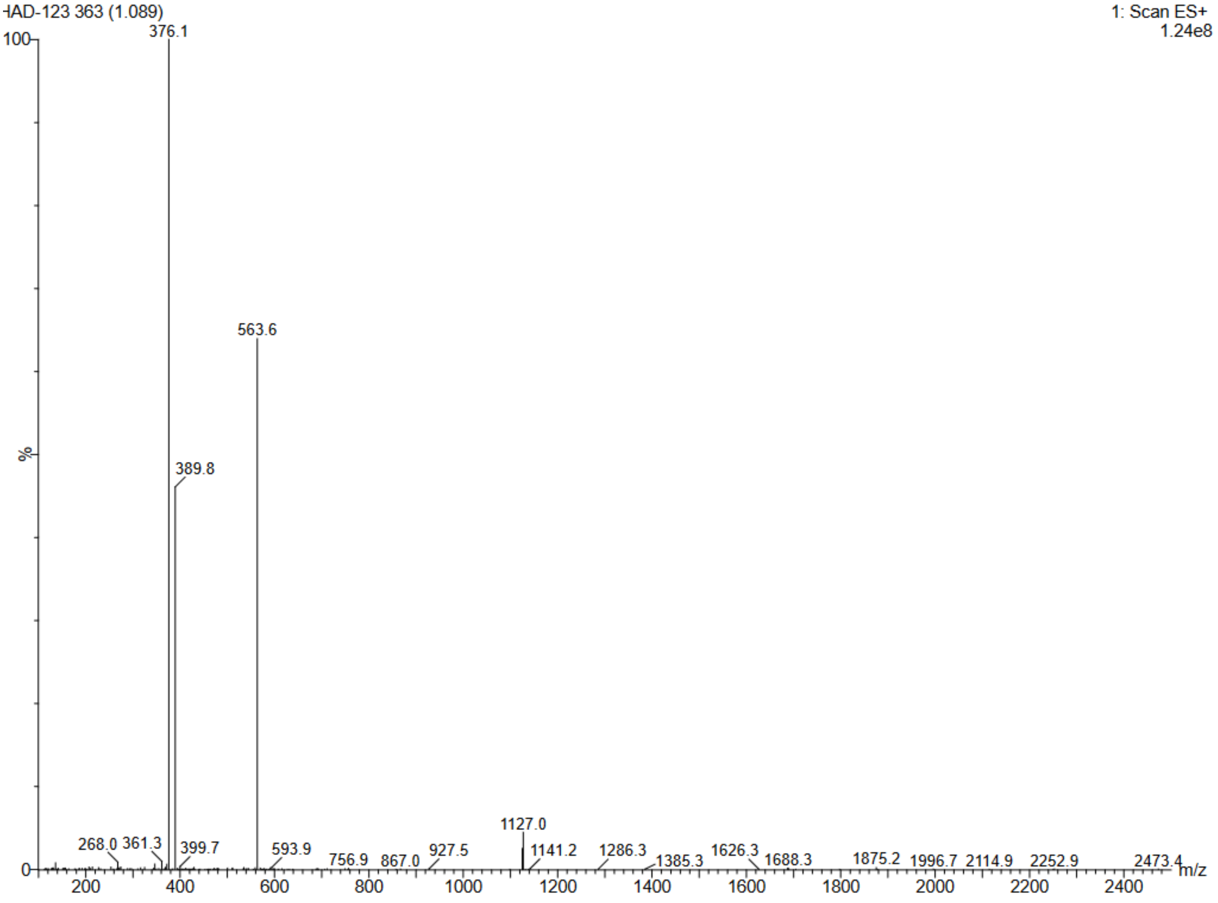

Compound 29c : D-R-V-Y-I-H-P-F(4Cl)

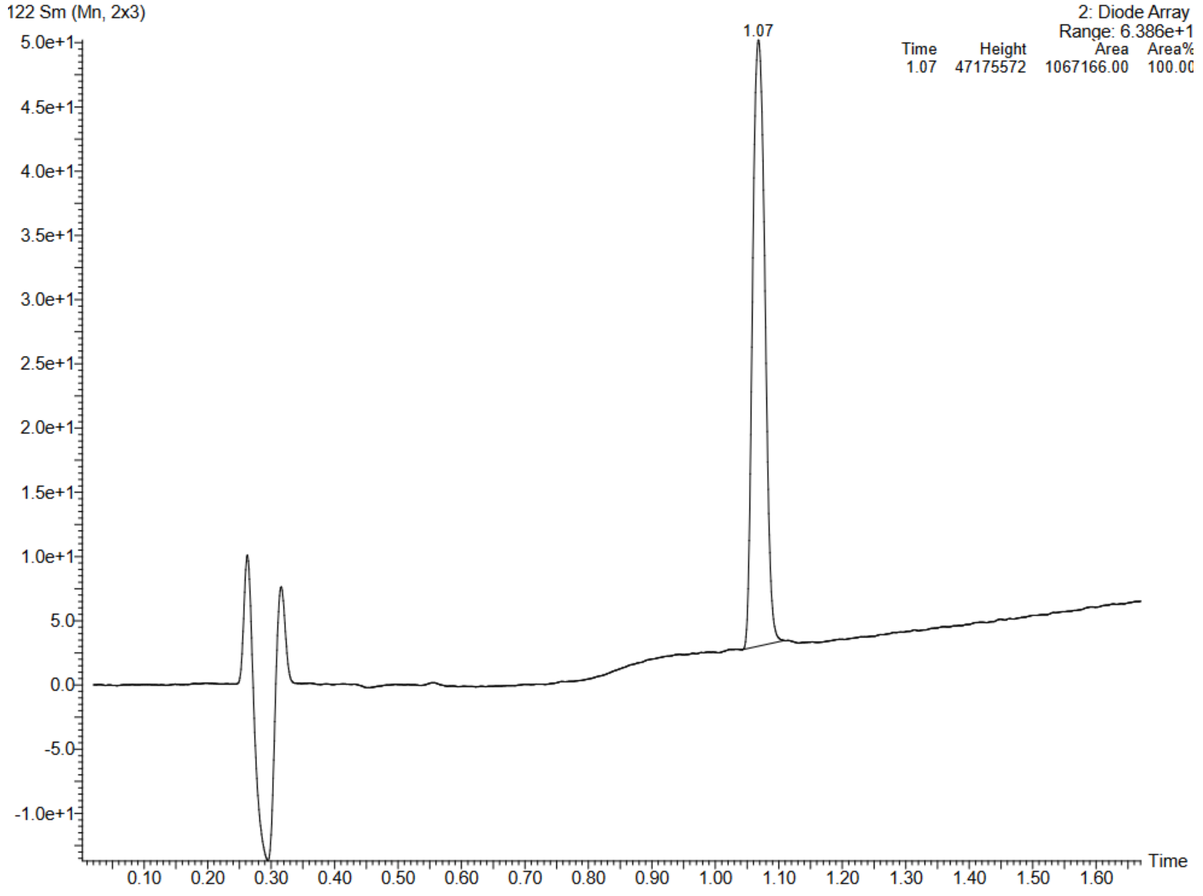


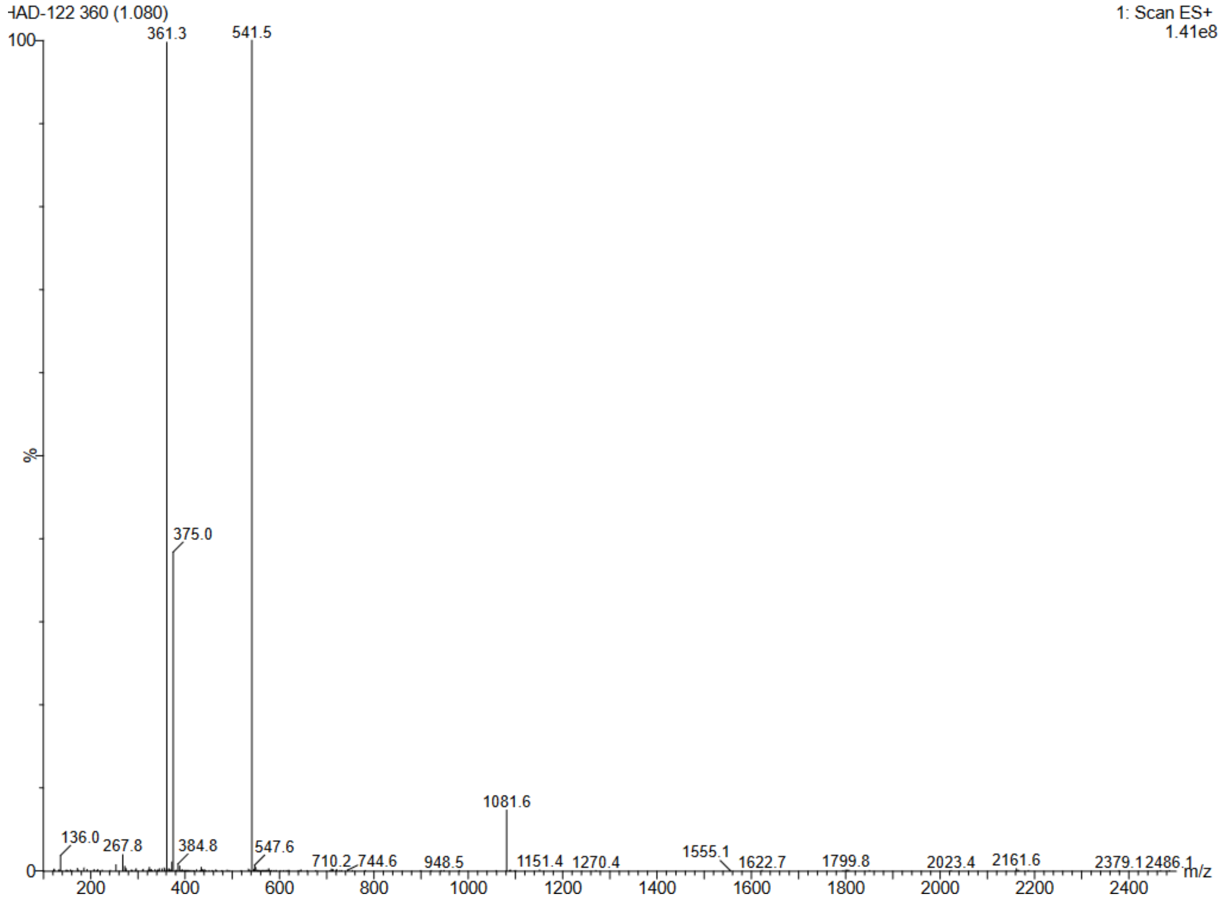

Compound 30c : D-R-V-Y-I-H-P-F(4F)

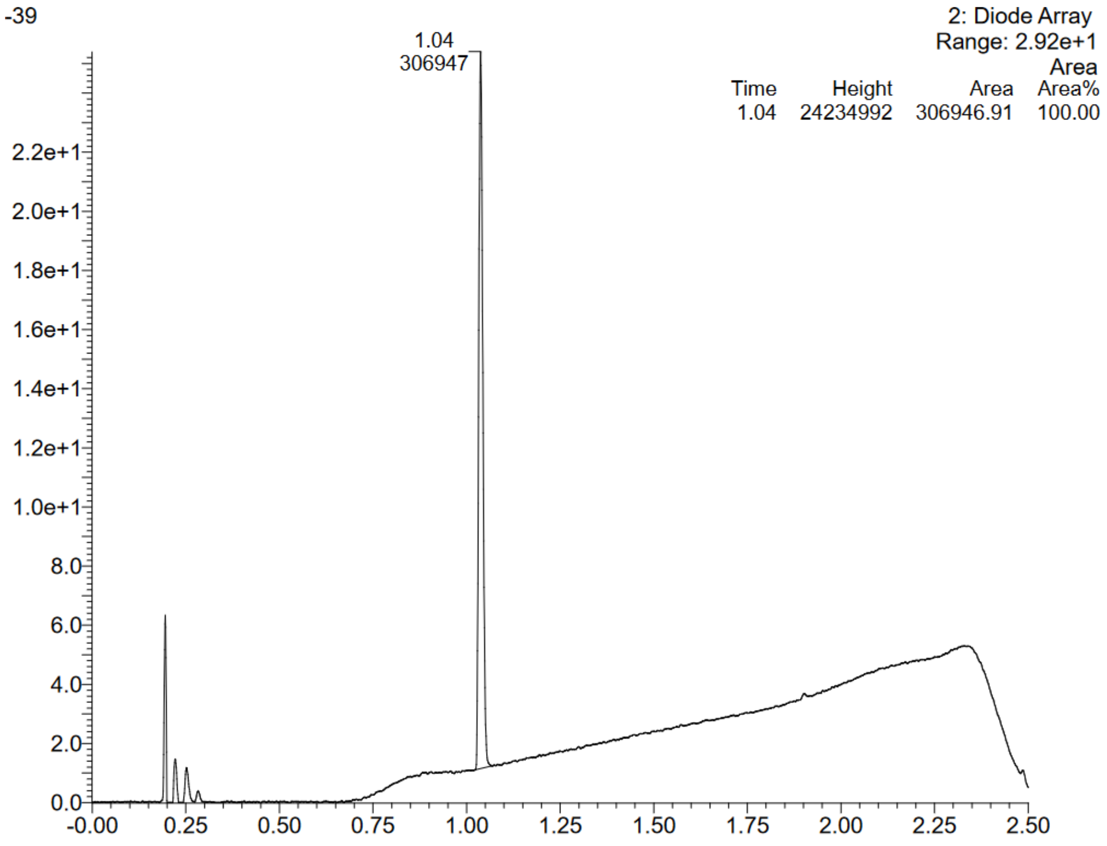


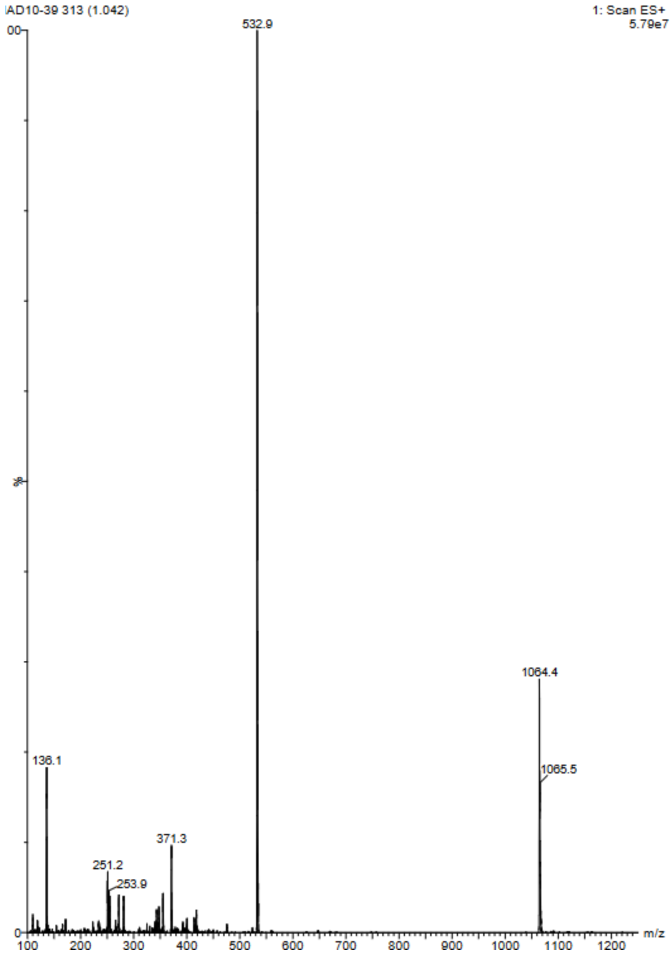
